# Segmented wavetrains and sites of reversal in the mouse seminiferous tubules

**DOI:** 10.64898/2026.02.06.703668

**Authors:** Kei Sugihara, Ayuki Sekisaka, Toshiyuki Ogawa, Takashi Miura

**Author notes:** Corresponding author *Email address:* (Kei Sugihara).

## Abstract

Mammalian spermatogenesis occurs in the seminiferous tubules, which exhibit unique spatiotemporal differentiation patterns known as cellular association patterns. In mice, these patterns can be regarded as one-dimensional wavetrains that consistently propagate inward from both ends, resulting in one or more “sites of reversal.” Segmented wavetrain pattern, in which the wave propagation direction spatially switches, was observed in our previous three-species reaction-diffusion model for interspecific species difference in spermatogenic waves (Kawamura et al., 2021). However, the biological mechanisms of the formation of sites of reversal and of this directional bias, as well as the principle of pattern formation, remain unknown. Here, we refined our previous model to match the actual biological spatiotemporal scale and examined its dynamics through extensive numerical simulations. The modified model frequently generated segmented wavetrain patterns, corresponding to the sites of reversal, but without directional bias. We systematically examined possible biological mechanisms for the bias and found that tubule elongation, especially near the rete testis, most effectively accounts for the bias among the tested. Extensive simulations revealed that the segmented pattern is numerically stable, emerges more frequently in longer domains, and shows an exponential segment size distribution with a lower limit for the stably existing segment length. These explorations imply that locally emerged unidirectional wavetrains serve as building blocks to generate the stable segmented wavetrains through their interactions.

**Highlights:**

- Segmented wavetrains reflect sites of reversal in seminiferous tubules.
- Segmented patterns frequently emerge but show no inherent directional bias.
- Tubule elongation may contribute to inward propagation near the rete testis.
- Segmented wavetrains are numerically stable and more frequent in longer domains.
- Interactions of local unidirectional wavetrains generate stable segmented structures.

## 1. Introduction

### 1.1. The structure of mouse seminiferous tubules

Mammalian sperm are generated in a tube-like structure called the seminiferous tubules (Fig. 1a, (2015)). In the mouse testis, these tubules are 200-300 *µ*m in diameter, U-shaped, and both ends are connected to the rete testis (Fig. 1a). Each seminiferous tubule consists of specialized large epithelial cells called Sertoli cells (Fig. 1b). The germ cells, which eventually develop into sperm cells, are embedded between Sertoli cells. Stem cells are located at the outermost region of the seminiferous tubules. As the differentiation proceeds, the cells gradually move to the inner region. At the end of differentiation, mature sperm are released into the lumen and transported to the rete testis by the coordinated flow inside the tubules (Fleck et al., 2021).

**Figure 1:**
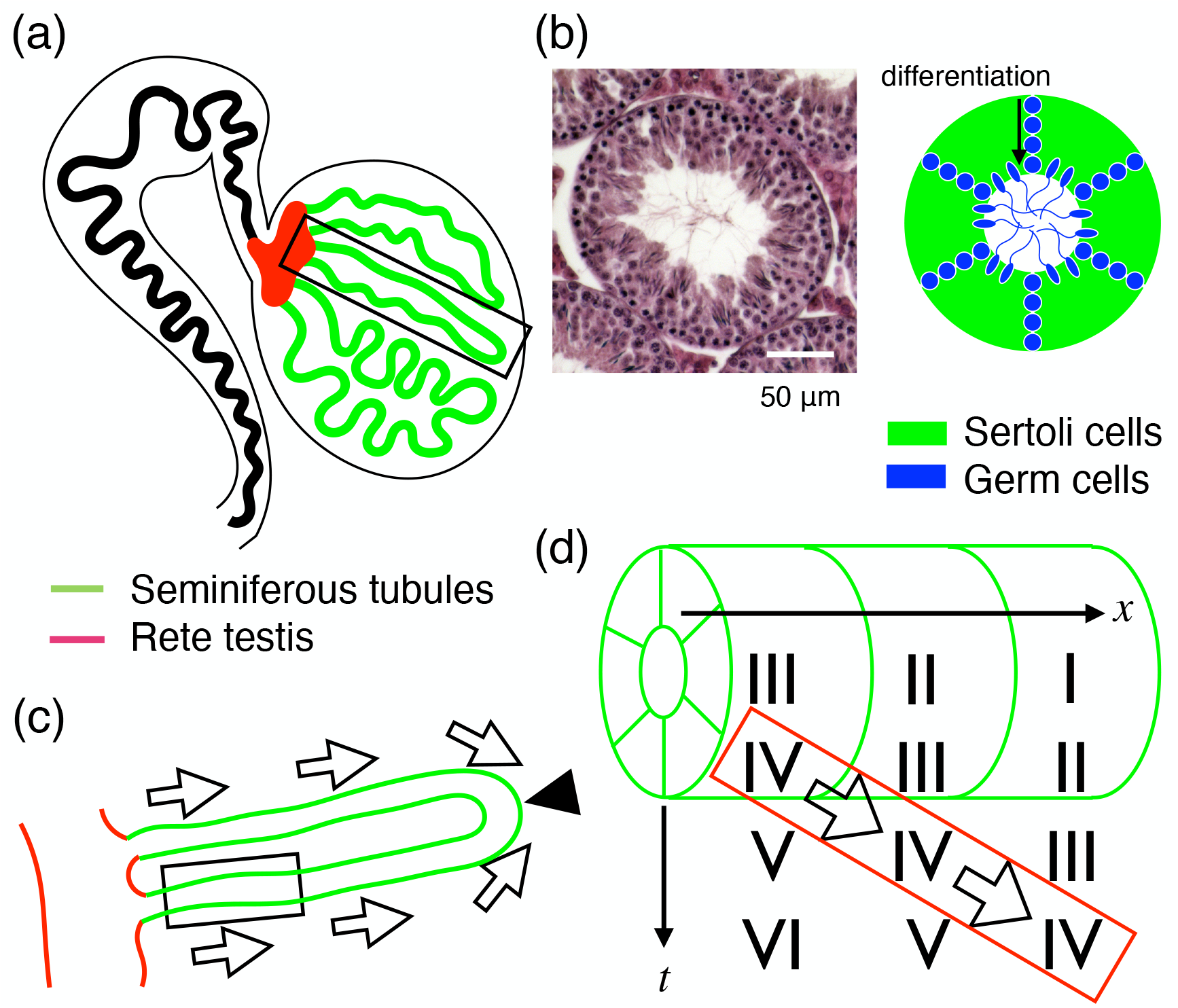
Background of this study. (a) Gross anatomy of the testis. Sperm are generated from germ cells in seminiferous tubules (green), which are transferred to the epididymis via the rete testis (red) and the efferent ducts. (b) Histological structure of the seminiferous tubule. Left: a cross-section of mouse seminiferous tubules (hematoxylin and eosin staining). Right: Schematic representation. The tubule consists of large Sertoli cells (green), and germ cells (blue) are embedded in the epithelial sheet of Sertoli cells. (c) A seminiferous tubule is a U-shaped epithelial tube. The cellular association pattern moves from both edges (inward direction), and as a result, at least one site of reversal always exists (black arrowhead). (d) Each location of the tubules is characterized by the differentiation state pattern of germ cells. Inside the tubules, the alignment of germ cells shows a specific pattern, and the pattern was classified into twelve stages (Roman numerals). As the stage progresses locally at each location, the longitudinal alignment of the stage moves unidirectionally (white arrow).

### 1.2. The cellular association pattern and its characteristics

The differentiation pattern of the sperm exhibits unique spatiotemporal dynamics, known as the cellular association pattern (Fig. 1d, Ross and Pawlina (2015)). Locally, the differentiation state at each point in the tubules can be classified into twelve different stages in mice (Stage I-XII) (Meistrich and Hess, 2013). The stages progress over time in each location and repeat cyclically, which is called the seminiferous epithelium cycle. The differentiation pattern is not spatially independent, showing some spatiotemporal order. Stages are arranged sequentially inside the seminiferous tubule (Fig. 1d). As a result, the periodic pattern of stages moves locally in one direction (Fig. 1d, a red box). We note that these wavetrain patterns are defined for the stage of the seminiferous epithelium and its embedded germ cell differentiation, not the mature sperm distribution released into the lumen.

Based on detailed three-dimensional observations (Nakata et al., 2015, 2017), several known characteristics of cellular association patterns in mouse seminiferous tubules have been identified. Firstly, the direction of pattern progression is always from the edge of the U-shaped region (Fig. 1c, white arrows). As a result, there is always at least one site of reversal of the cellular association pattern (Fig. 1c, black arrowhead). In addition, there is stochasticity in the cellular association pattern, and sometimes more than one site of reversal was observed (Nakata et al., 2017). Despite previous experimental observations, the mechanisms that generate sites of reversal and directional bias in wave propagation remain unknown.

### 1.3. Mathematical models that generate wavetrain patterns

Such wavetrain patterns can be generated in a variety of mathematical systems showing Turing-wave instability. This instability can occur when the zero wavenumber, spatially uniform mode is stable, but finite, non-zero wavenumber modes become unstable, with complex eigenvalues. This type of model is mentioned in the original Turing model paper (Turing, 1952). To implement these characteristics, we require three or more variables in the reaction-diffusion framework when nonlocal terms are excluded. It has also been reported that the complex Swift-Hohenberg equation can generate this type of pattern (Sakaguchi, 1997).

### 1.4. Mathematical modeling of wavetrain formation in the seminiferous tubules

To understand wavetrain patterns in the seminiferous tubules, we previously proposed a three-species reaction-diffusion model (Eq. (1) with *µ* = 0, Fig. 3a,c, (2021)). We adopted a phenomenological approach because little is known about the molecular mechanisms regulating spermatogenic waves except for diffusive retinoic acid signaling. We constructed one of the minimal reaction-diffusion models without nonlocal terms, which exhibits a wave bifurcation. In this bifurcation, a Hopf bifurcation occurs at a non-zero wavenumber mode before the spatially uniform mode becomes unstable (Ogawa, 2007). In our previous work, we demonstrated that the interspecific difference between helical wavetrains in humans and longitudinal ones in mice can be attributed to the relationship between the generated pattern size and the circumferential domain size.

### 1.5. Segmented pattern in wavetrains

In addition to these two patterns, we frequently observed segmented wavetrain patterns in long domains, where the direction of the wavetrain propagation switched at one or more spatial positions (Fig. 6, Supplementary Movie 6). While similar behaviors were also reported in the complex Swift-Hohenberg system (Sakaguchi, 1997), the nature of such segmented patterns remained unclear.

The numerical simulations in the previous study were performed under generalized settings of domain size, timescale, and boundary conditions, which may not accurately reflect the biological scale of mouse seminiferous tubules. Although we speculated this pattern may correspond to the sites of reversal *in vivo*, its biological relevance has not yet been verified. This study discussed only the pattern geometry, not the direction of wave propagation. Furthermore, the principle of segmented wavetrain pattern formation remains unknown, as does whether they correspond to the actual solutions of the system.

### 1.6. Research purposes

In the present study, we focused on two central questions: (1) whether the model can reproduce the sites of reversal and inward directional bias near the rete testis at biologically relevant scales, and (2) what principle underlies the emergence of segmented wavetrain patterns. To address these questions, we first modified our previous model to better reflect the actual spatiotemporal dynamics in the mouse seminiferous tubules. Then, we quantified the frequency and directional bias in the model and explored possible additional mechanisms contributing to the directional bias. Finally, through extensive numerical simulations, we explored the principles governing the emergence and stability of segmented wavetrain patterns.

## 2. Model and methods

### 2.1. Model definition

We employed a three-species reaction–diffusion system, which exhibits Turing–wave instability, as introduced in our previous study (Kawamura et al., 2021):

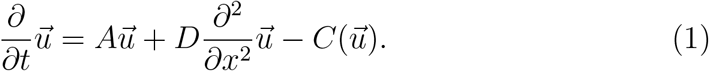

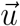, *A*, and *D* denote a three-species vector 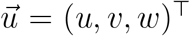, 3#x00D7;3 reaction and diffusion term matrices, respectively. Biologically, *u, v*, and *w* can be interpreted as deviations from the equilibrium concentration of hypothetical molecules that control spermatogenic staging rather than germ or sperm cell density. We assumed that each stage corresponds to the combination of the level of each molecule, for example, low *u*, low *v*, and high *w*. The species *w* without a diffusion term can be interpreted as an intracellular molecule that does not diffuse outside the cell (e.g., a transcription factor). We emphasize that a two-species reaction-diffusion system without nonlocal terms cannot exhibit wavetrains. Hence, a three-species system is one of the simplest possible models in the reaction-diffusion framework. 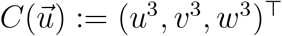 is a cubic term to prevent diverging to infinity. As a phenomenological model, we chose this nonlinear term as one of the simplest possible forms. Linear stability analysis and weakly nonlinear analysis were discussed for the original model in detail in Kawamura et al. (2021).

Unless otherwise specified, we utilized the following parameter sets:

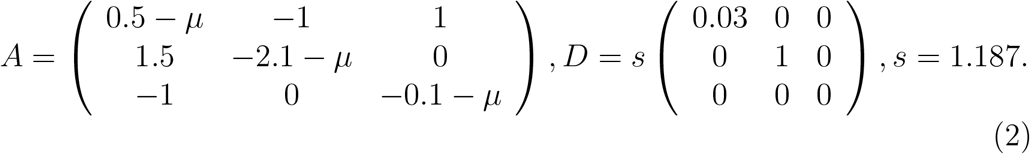

In this paper, we introduced the control parameter *µ* to regulate the growth speed of the wave. *s* is the spatial scaling factor to regulate the wavelength of the emerging pattern, which is proportional to 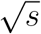. We set this specific value of *s* so that the most unstable wavelength 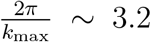. For initial conditions, white noises or predetermined waves for *u, v*, and *w* were used. We performed numerical simulations in one-dimensional space with periodic boundary conditions or zero-flux (Neumann) conditions unless stated otherwise. Numerical simulations were performed using finite-difference methods and implemented in C. Reaction terms were calculated using the fourth-order Runge-Kutta method. Diffusion terms were calculated using the explicit central difference method. The initial white noise was generated using Mersenne Twister (Matsumoto and Nishimura, 1998).

In our model, linear stability analysis shows that the most unstable wavelength is 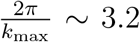. Weakly nonlinear analysis with *µ* = 0 shows that the typical wave speed *c* is *c* ∼ 0.51 for *k* = *k*_max_, yielding a typical time period of around *T* ∼ 6.2. In the mouse, the average length of the complete waves at postnatal day 90 was around 1.8 cm (Nakata et al., 2017). The duration of a cycle is widely known as 8.6 days (Oakberg, 1956). By introducing linear variable transformation for *t*^′^ := *c*_*t*_*t* and *x*^′^ := *c*_*x*_*x*, we can rewrite our system as

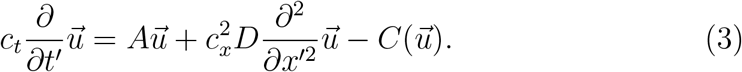

We can choose the values of *c*_*t*_ and *c*_*x*_ so that one typical wavelength corresponds to 1.8 cm and one period corresponds to 8.6 days. Hereafter, we use these *x*^′^ and *t*^′^ as mere *x* and *t*, respectively.

### 2.2. Estimation of wave velocity from numerical simulations

To analyze segmented patterns, we numerically estimated wave velocity with sign *c* from the numerical simulation results. If a specific segment of *u* moves with the velocity *c, u* can be expressed as *u*(*x, t*) = *f* (*x* − *ct*) using a univariate function *f* and thus

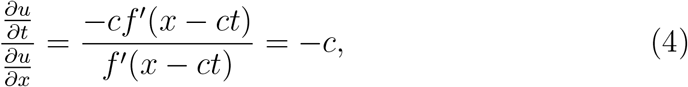

when *f*^′^(*x* − *ct*) ≠ 0. Therefore, we numerically estimate the wave velocity *c*_*e*_ as

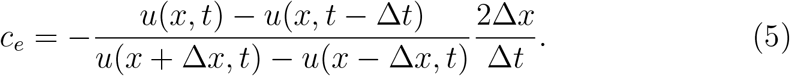

Note that a central and backward difference scheme is applied in the spatial and temporal directions, respectively.

### 2.3. Quantification of segment size distribution

For the quantification of segment size distribution, local wave velocity *c*_*e*_ was estimated for each location *x* at a time step (Fig. 2a,b). Here, we assume each segment has the same wavenumber and thus the same absolute value of wave velocity. Because wave velocity estimation is expected to be inaccurate where *f*^′^ ∼ 0, there were extremely large absolute values and large fluctuations along the *x* axis (Fig. 2b). To ameliorate this irregularity, we took only the sign of *c*_*e*_ (Fig. 2c) and applied the Gaussian filter along *x* direction (Fig. 2d). We note that the resulting function was not entirely smooth because the smoothing was applied to the discontinuous function between -1 and 1. We defined the segment size as the size of the region where the smoothed sign of *c*_*e*_ is continuously the same.

**Figure 2:**
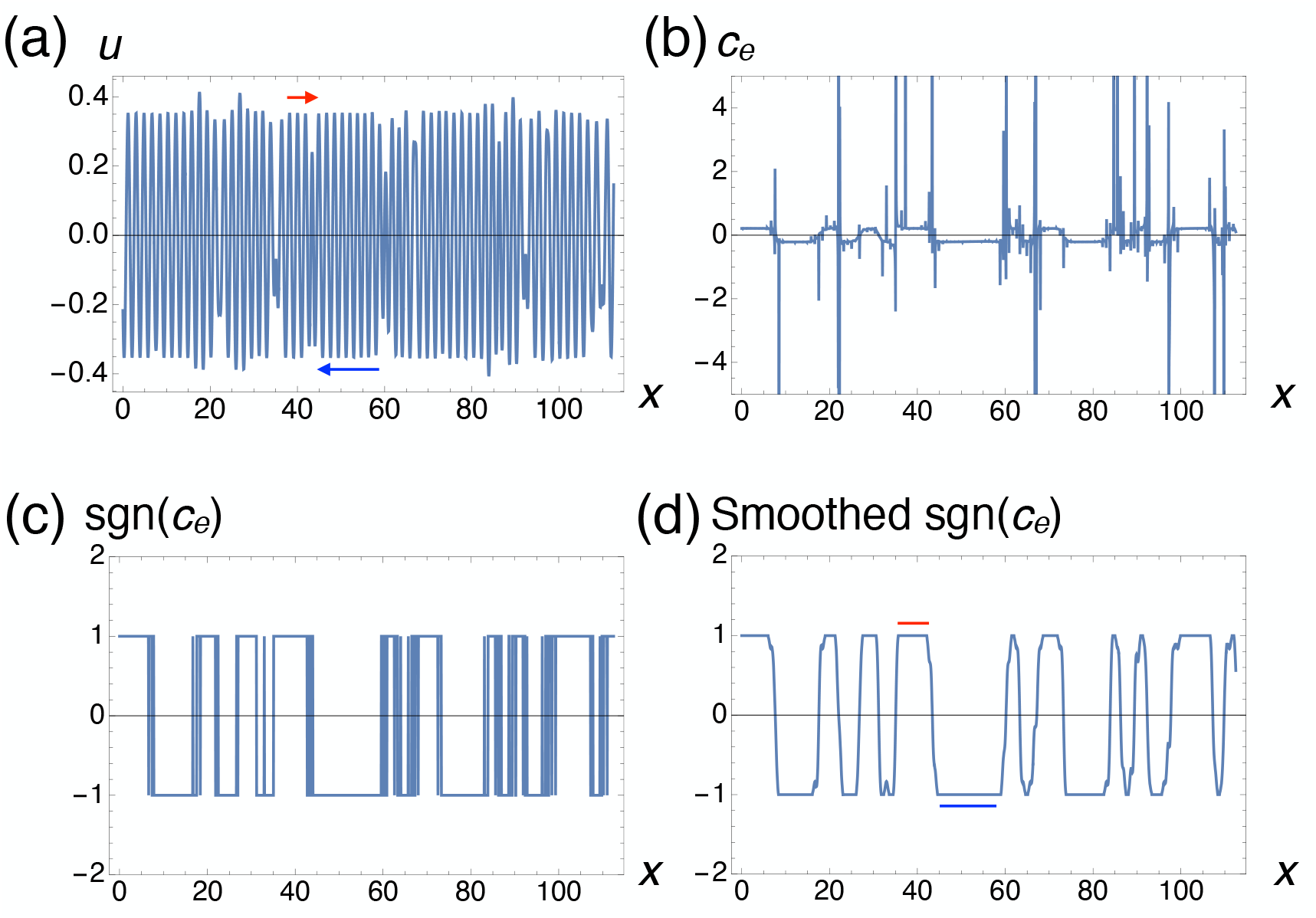
Numerical methods to identify the segments. (a) A representative original numerical simulation result for *u*. Blue and red arrows are examples of segments with positive and negative wave directions, respectively, as identified in (d). (b) Local wave velocity *c*_*e*_ estimated in numerical simulations. (c) Sign of *c*_*e*_, sgn(*c*_*e*_). (d) Gaussian filtered sgn(*c*_*e*_). Blue and red lines are the intervals of the same positive and negative sign values corresponding to (a).

### 2.4. Experimental procedures

The procedures are based on Kawamura et al. (2021). Briefly, adult male mice euthanized by skilled cervical dislocation were dissected. Testes were fixed in Bouin’s fixative, and their paraffin-embedded specimens were made into 10-µm-thick sections. The sections were stained with hematoxylin and eosin, and observed using a BZ-X700 microscope (Keyence, Osaka, Japan).

## 3. Results

### 3.1. Emergence of segmented patterns at the biologically relevant spatiotemporal scale

#### 3.1.1. Model modification

We first modified our previous model (Kawamura et al., 2021) to match the actual spatiotemporal scale *in vivo*. In mice, the seminiferous tubules are approximately 6 mm long at birth and grow to about 14 cm by 90 days of age (2017), after which their length remains largely unchanged (Nakano et al., 2021). Thus, the domain length should be set to approximately eight cycles of the wavetrain pattern in the numerical simulations. Since the seminiferous tubules are connected to the anatomically distinct structures at both ends, the boundary conditions were set as zero-flux (Neumann) conditions.

Sexual maturity usually happens by six weeks of age in male mice (Lambert, 2009). When we consider that the pattern formation starts at birth, taken together, the model should show the grown wave patterns after five cycles (corresponding to 43 days old). We also note that their life spans are typically 2-3 years (Garratt et al., 2022), so the simulations for 100 cycles (corresponding to 2.35 years) are sufficient for comparing the dynamics to the actual biological phenomena.

When we performed the numerical simulations of our previous model with the above settings, we found that the patterns did not grow enough at 43 days (Fig. 3a upper, c left), compared to 860 days (Fig. 3a lower, c right). By increasing the initial noise amplitude and the growth speed of the system to the extent that the spatial uniform mode *k* = 0 is stable (*µ* = −0.07), we successfully observed sufficiently developed waves at 43 days (Fig. 3b,d, Supplementary Movie 1).

**Figure 3:**
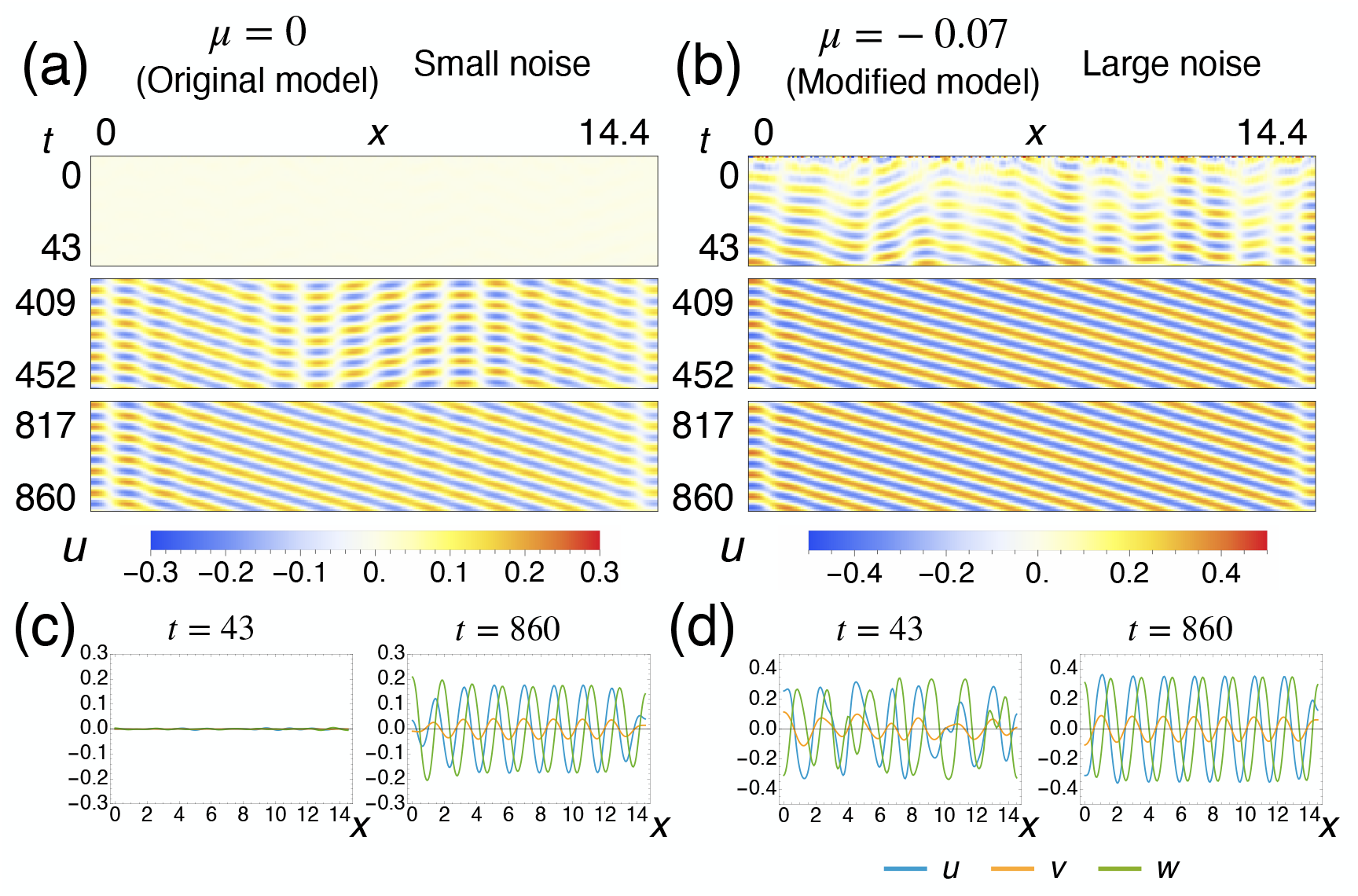
Model modification for faster pattern formation that reflects the actual spatiotemporal scale *in vivo*. Representative numerical simulation results for *u* in the original model (Kawamura et al., 2021) with the initial condition given by small-amplitude white noise (a,c) and the modified model with the initial condition given by larger-amplitude (50×) white noise (b,d). (a,b) *x* − *t* plot with *u* shown in color. Only initial, middle, and final time segments are visualized. (c,d) Representative snapshots for *u, v*, and *w*. Left: *t* = 43, corresponding to 7-week-old mice. Right: *t* = 860, corresponding to two-year-old mice.

In contrast, increasing only the system growth speed has minor effects on the early pattern growth at 43 days (Supplementary Fig. 1a upper, c left, Supplementary Movie 2). Increasing only the initial noise amplitude can speed up the pattern growth, but the standing wave components remain longer (Supplementary Fig. 1b middle). For various control parameters *µ* and initial noise amplitudes, we also quantitatively assessed relative wavetrain growth at *t* = 43 compared to *t* = 860 by evaluating the mean square of *u* (Supplementary Fig. 2a). We confirmed that both the initial noise size and *µ* accelerate the relative growth at *t* = 43. Hereafter, we used these conditions (*µ* = −0.07 with larger initial noise amplitude) unless otherwise stated.

#### 3.1.2. Frequency and directional bias of the segmented pattern

Next, we asked whether the segmented patterns and directional bias in wave propagation can be observed in the biologically relevant settings. We performed numerical simulations from the random white noise at a biologically relevant spatiotemporal scale (*t* = 860 and *L* = 14.4) with zero-flux boundary conditions (Fig. 4a, Supplementary Movie 3). Among 100 simulations, segmented patterns were observed frequently (83 out of 100, Fig. 4b). In some cases, more than two segments can be observed, consistent with some seminiferous tubules with multiple sites of reversal (Nakata et al., 2017).

**Figure 4:**
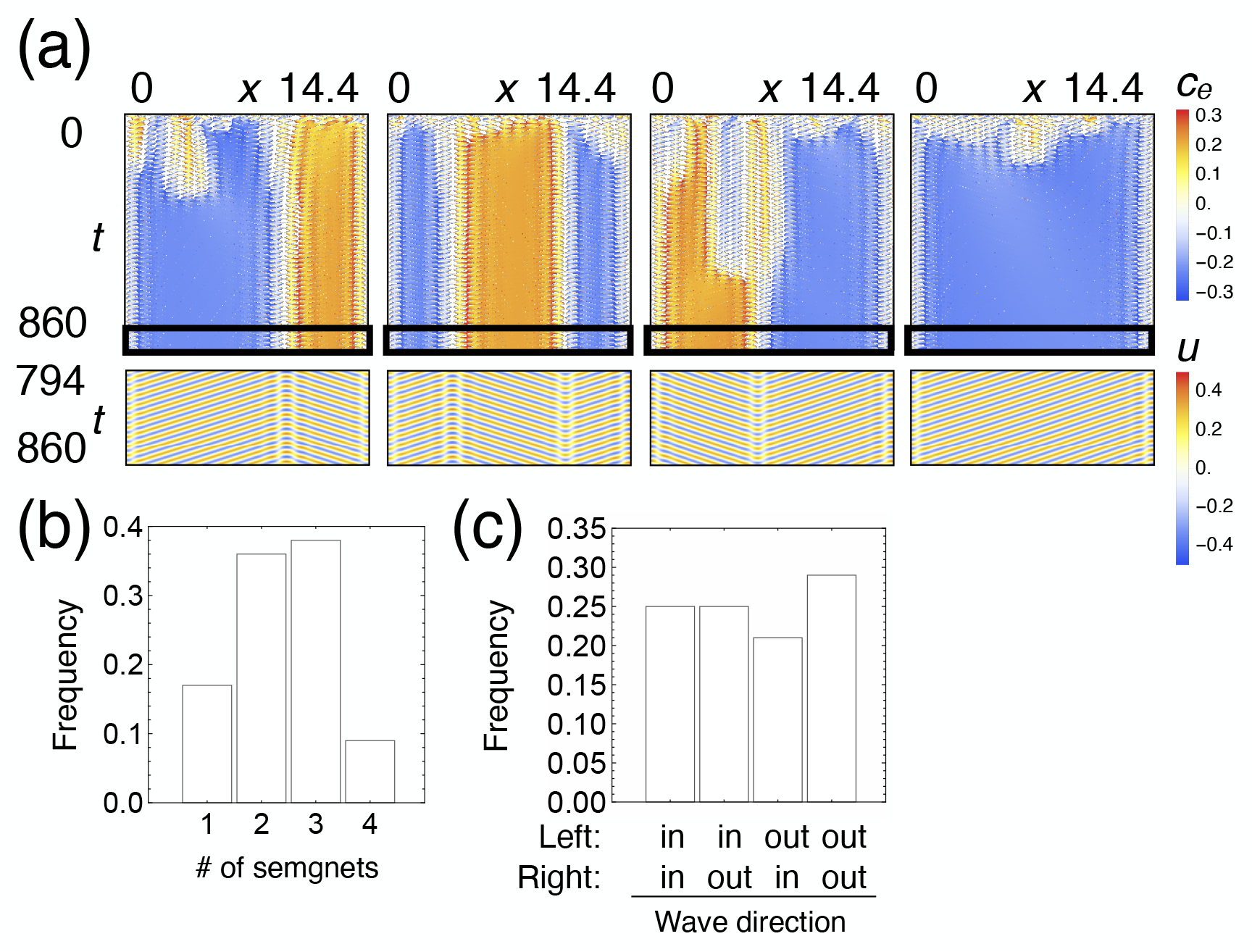
Frequency and directional bias of segmented patterns at the biologically relevant spatiotemporal scale. (a) Numerical simulation results shown as the estimated local velocity *c*_*e*_ (upper) and *x* − *t* plot with *u* shown in color (lower) for *L* = 14.4 with Neumann boundary conditions. (b) The distribution of segment number, corresponding to (a) (*n* = 100). (c) The frequency of wave direction patterns at both edges, corresponding to (a). No significant bias was observed (*p* = 0.73; Pearson’s *χ*^2^ test under the null hypothesis of equal probability; *n* = 100).

However, we did not observe any directional bias at both edges (Fig. 4c). The model system itself is spatially symmetric at the level of the governing equations. Therefore, under periodic boundary conditions or in an infinite domain with unbiased initial conditions, there should be no preference for wave propagation direction. This absence of bias was also true for zero-flux boundary conditions here. In contrast, the observation of mouse seminiferous tubules revealed that the wavetrain pattern progresses from proximal to distal at the openings (Fig. 1b, (Curtis, 1918; Nakata et al., 2017)). This discrepancy highlights the necessity for other mechanisms to bias the wave propagation direction near the rete testis *in vivo*.

#### 3.1.3. Effect of tubule elongation on wave directional bias

The numerical simulations suggest the presence of an additional mechanism that affects the wave direction. However, the biological basis for this directional control remains unknown. Therefore, we examined possible hypothetical mechanisms to enable this bias with various unfixed initial conditions: (1) the expansion of spermatogenic domains (Nakata et al., 2017), (2) the flow inside the seminiferous tubules (Maekawa et al., Middendorff et al., 2002), (3) tubule diameter changes near the rete testis (Nakata et al., 2015), (4) distinctive signaling activity in the rete testis (Nagasawa et al., 2018; Teletin et al., 2019; Imura-Kishi et al., 2021), and (5) tubule elongation during development (Nakata et al., 2017). Among these, the tubule elongation most plausibly explains the inward propagation bias near the rete testis *in vivo*. Here, we focus on this most plausible mechanism. The analyses of the other mechanisms that did not reproduce the bias consistent with the observation are discussed in Appendix A.

During development, seminiferous tubules drastically elongate from ∼ 6 mm at birth to around 14 cm in mice before postnatal day 90 (Nakata et al., 2017). We wondered if such domain growth could affect the wave direction near the boundary.

First, as the simplest hypothesis, we considered the uniform domain growth (Fig. 5a). We implemented the domain growth by gradually increasing the spatial grid size 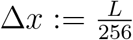 and stopping it midway, so that the domain size *L* satisfies:

**Figure 5:**
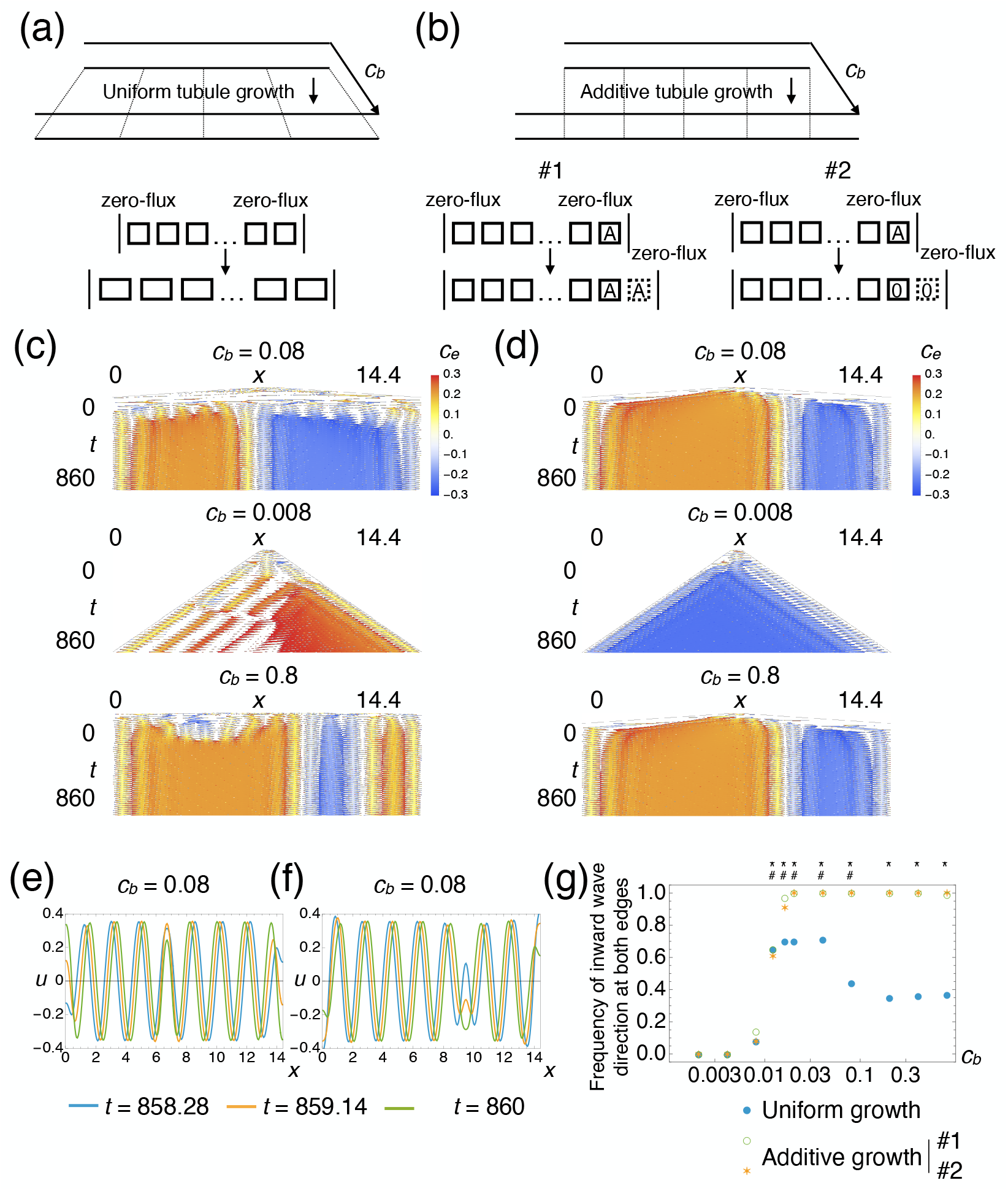
Elongation of seminiferous tubules may contribute to the inward wave direction near the rete testis. (a,b) Schematic representation of two modes of tubule elongation. (a) Uniform tubule growth. (b) Additive tubule growth at both sides of the boundaries. For (b), two numerical implementation methods were considered. (c,d) *x* − *t* plot of local wave velocity *c*_*e*_ for different tubule growth speeds *c*_*b*_ with uniform domain growth (c) or additive domain growth (d). (e,f) *x* − *u* plot for three different time points with uniform domain growth (e) or additive domain growth (f). (g) The effect of the tubule growth speed *c*_*b*_ on the wave direction bias with two modes of domain growth. #: *p <* 1 × 10^− 13^ for uniform growth, ∗: *p <* 1 × 10^− 13^ for additive growth (binomial test under the null hypothesis П = 0.25; one-sided; *n* = 100).

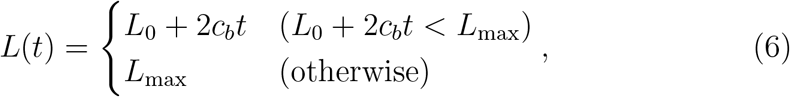

where the initial domain length is *L*_0_ = 0.56 and the maximum length is *L*_max_ = 14.4.

For intermediate values of *c*_*b*_ (0.01 ≤ *c*_*b*_ ≤ 0.1), a segmented pattern with inward wave propagation near both boundaries was observed significantly more often than random, reaching approximately 70% at the maximum (Fig. 5c upper, e, g, Supplementary Movie 4). However, for lower *c*_*b*_, unidirectional wavetrains were observed (Fig. 5c middle, g). For higher *c*_*b*_, the bias in wave direction disappeared (Fig. 5c lower, g). They were consistent with the biologically relevant *c*_*b*_, estimated as (14-0.6) cm/90 days/2 ∼ 0.07, based on the assumption that both border linearly grows at the same constant speed.

Then, we also considered the case of non-uniform elongation, as an extreme case where the elongation only occurs at both ends of the tubules. This hypothesis is inspired by the reports that Sertoli cells near the rete testis retain proliferative activity for a longer period after birth than those in the convoluted tubule in hamsters and rats (Aiyama et al., 2015; Figueiredo et al., 2016). To assess this effect, we performed numerical simulations in the peripherally growing domain. We numerically implemented domain growth by adding lattices at the periphery, so that both boundaries grow at a constant speed in two ways while keeping the spatial grid size fixed (Fig. 5b).

For intermediate and high *c*_*b*_ values (*c*_*b*_ ≥ 0.01), the segmented pattern with inward wave directions at both ends appeared more frequently than random (Fig. 5d upper, lower, f, g, Supplementary Movie 5). Moreover, for *c*_*b*_ ≥ 0.02, the bias was observed in all of 100 simulations. However, the wave direction bias disappeared for lower *c*_*b*_ (Fig. 5d middle, g). For both implementation methods, the degree of the bias depended on the tubule growth speed *c*_*b*_ (Fig. 5g).

Taken together, these results suggest that tubule elongation during development, particularly near the rete testis, may contribute to the inward propagation of spermatogenic waves.

### 3.2. Exploration of the principle of pattern formation through numerical simulations

The mathematical rigorous analysis of the segmented patterns in our model has not yet been obtained. Alternatively, we explored their dynamics through extensive numerical simulation to uncover the underlying principles of pattern formation.

#### 3.2.1. Numerical stability of the segmented patterns

First, to investigate whether the segmented patterns numerically show stability, we performed numerical simulations over a considerably long period (*t* ∼ 1.4 × 10^6^). The pattern was observed to persist during extended numerical simulations, suggesting that it reflects an actual solution of the system rather than a numerical artifact (Fig. 6a, Supplementary Movie 6). We estimated local wave velocity *c*_*e*_ as described in Section 2.2 and found that all segments have similar absolute velocity values but with different directions (Fig. 6b,c).

**Figure 6:**
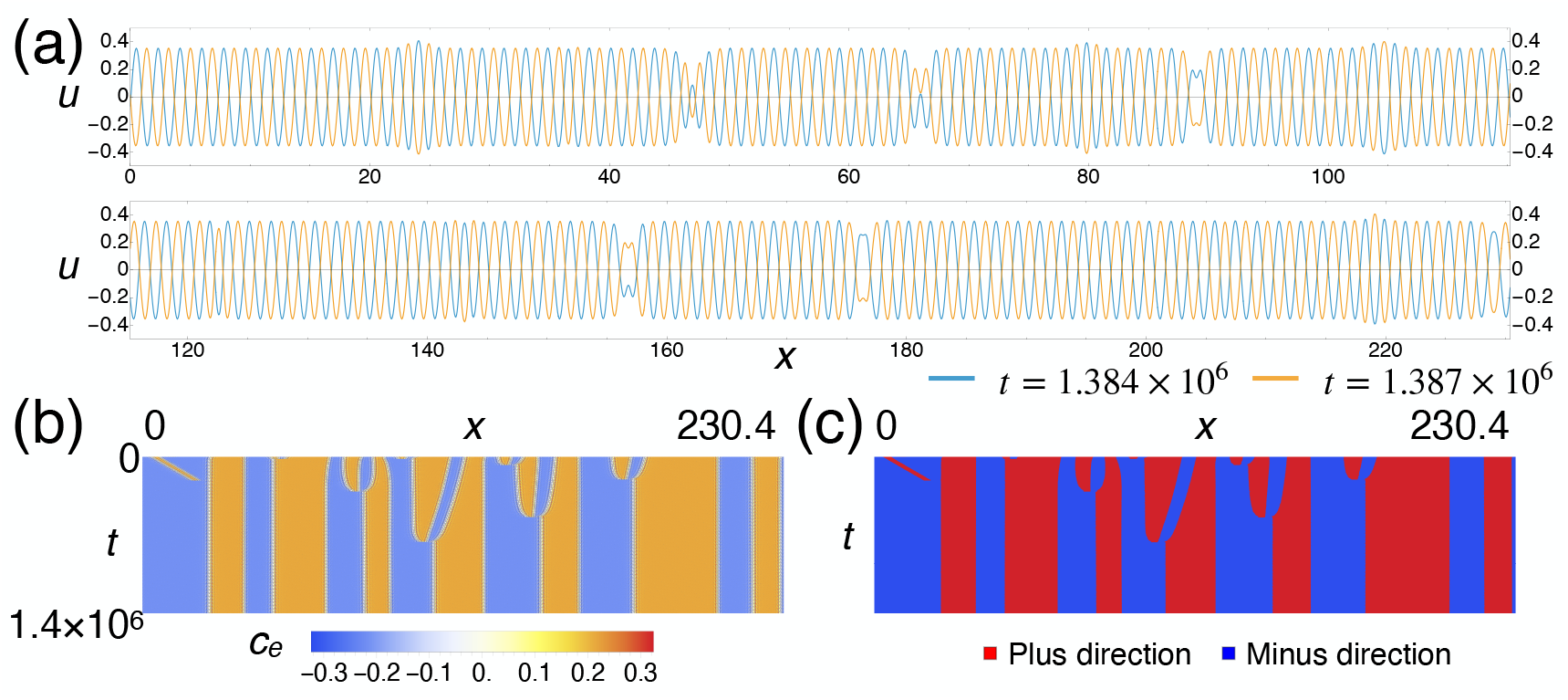
Long-term numerical stability of the segmented pattern. (a) Values of *u* were plotted at two time points, split into *x*-direction. The waves shift while the irregular part remains stationary, suggesting a segmented pattern. (b) The estimated local wave velocity *c*_*e*_, corresponding to (a). *x* − *t* plot with *c*_*e*_ shown in color. (c) Identified segments shown in *x* − *t* plot. The red and blue segments correspond to the plus and minus wave directions, respectively.

#### 3.2.2. Segmented pattern is frequently observed in longer domains

Next, we wondered how frequently those patterns emerge from white noise if segmented patterns exist as a stable solution of the model (Eq. (1)). We performed numerical simulations at the biologically relevant timescale for different domain lengths *L* with randomly selected white noise initial conditions (Fig. 7a). Here, we present numerical simulations using large domain sizes to elucidate the mathematical characteristics of the pattern. We did not observe any segmented pattern when *L* ≤ 3.6 (Fig. 7a, b blue). However, as the domain length increased, segmented patterns emerged more frequently, and we only observed segmented patterns when *L* ≤ 57.6 (Fig. 7a, b blue).

**Figure 7:**
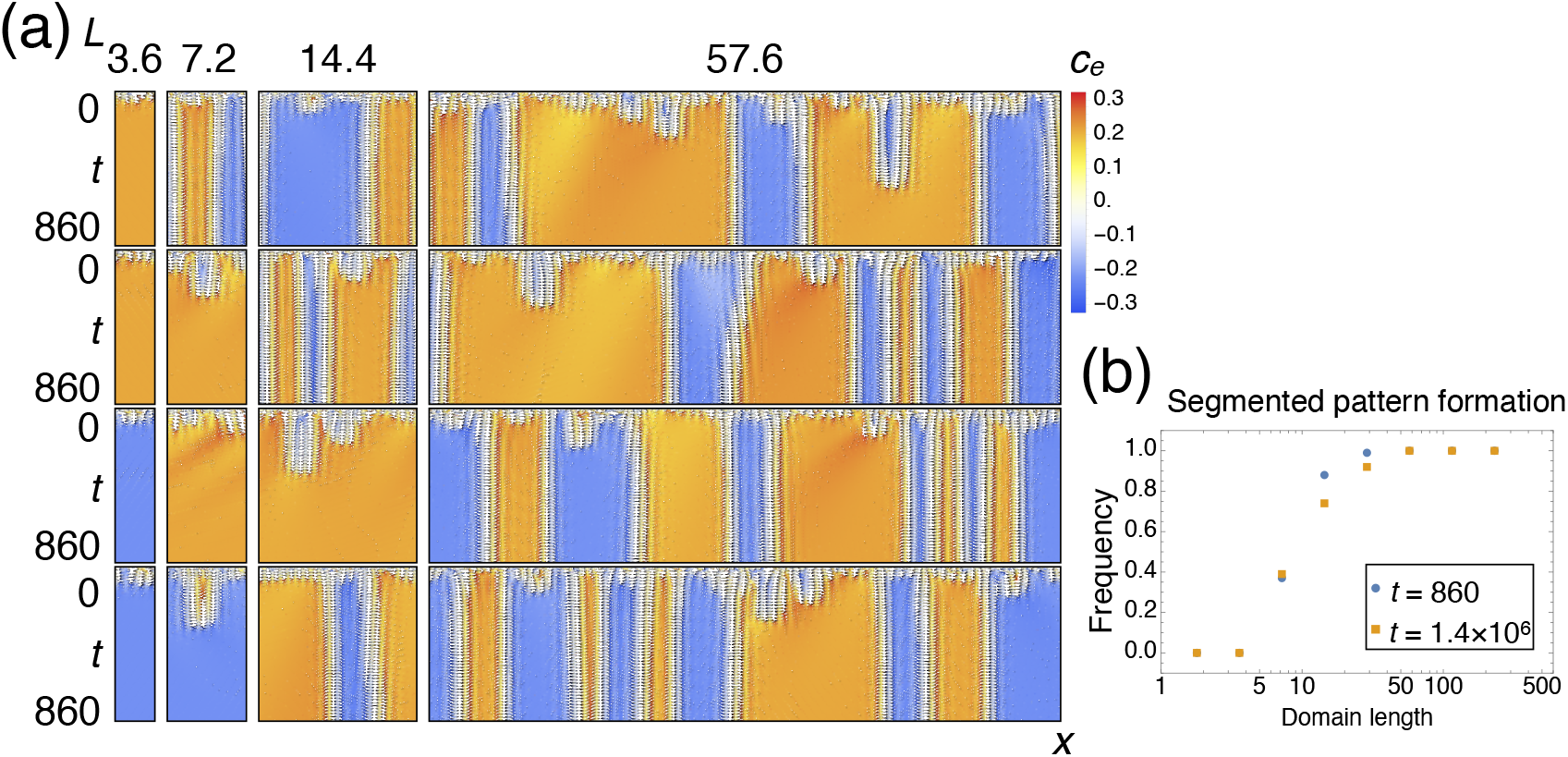
Domain length affects the frequency of segmented pattern formation. (a) Numerical simulation results shown as the estimated local velocity *c*_*e*_ for different domain lengths with periodic boundary conditions. (b) Frequency of segmented pattern emergence for various domain lengths *L* at *t* = 860 and *t* = 1.4 × 10^6^. 100 independent random white noises were given as the initial conditions.

We also assessed the pattern formation frequency on a much longer timescale (*t* = 1.4 × 10^6^). The frequency decreased with time, but it still depended on the domain size (Fig. 7d, blue vs. orange).

When we observed the time course of the long-term simulation results (Fig. 6b,c), we found that the initially generated shorter segments interact to form a smaller number of longer segments, which remain numerically stable for a prolonged period.

Furthermore, we also evaluated its dependency on the control parameter *µ* at a biologically relevant timescale (Supplementary Fig. 2b). The domain size dependency was observed throughout all *µ* we tested. These observations suggest that the frequency of segmented pattern formation is affected by the domain size.

#### 3.2.3. Segment size obeys an exponential distribution

Next, given that the initially emerged smaller segments appeared to interact with each other, we assessed the segment size distribution to understand how segmented patterns spontaneously emerge from fluctuation from a mathematical point of view. We performed the numerical simulations on a long one-dimensional domain, equivalent to 1024 times the most unstable wavelength, to minimize the influence of specific bounded intervals and boundary conditions when compared to the entire real space. The quantification method was described in Section 2.3.

We found that the size distribution largely obeys the exponential distribution at the biologically relevant timescale (Fig. 8a). We also systematically confirmed this tendency for different *µ* (Fig. 8b). We note that the half-decay length of the frequency tends to be longer when the system approaches the bifurcation (high *µ*).

**Figure 8:**
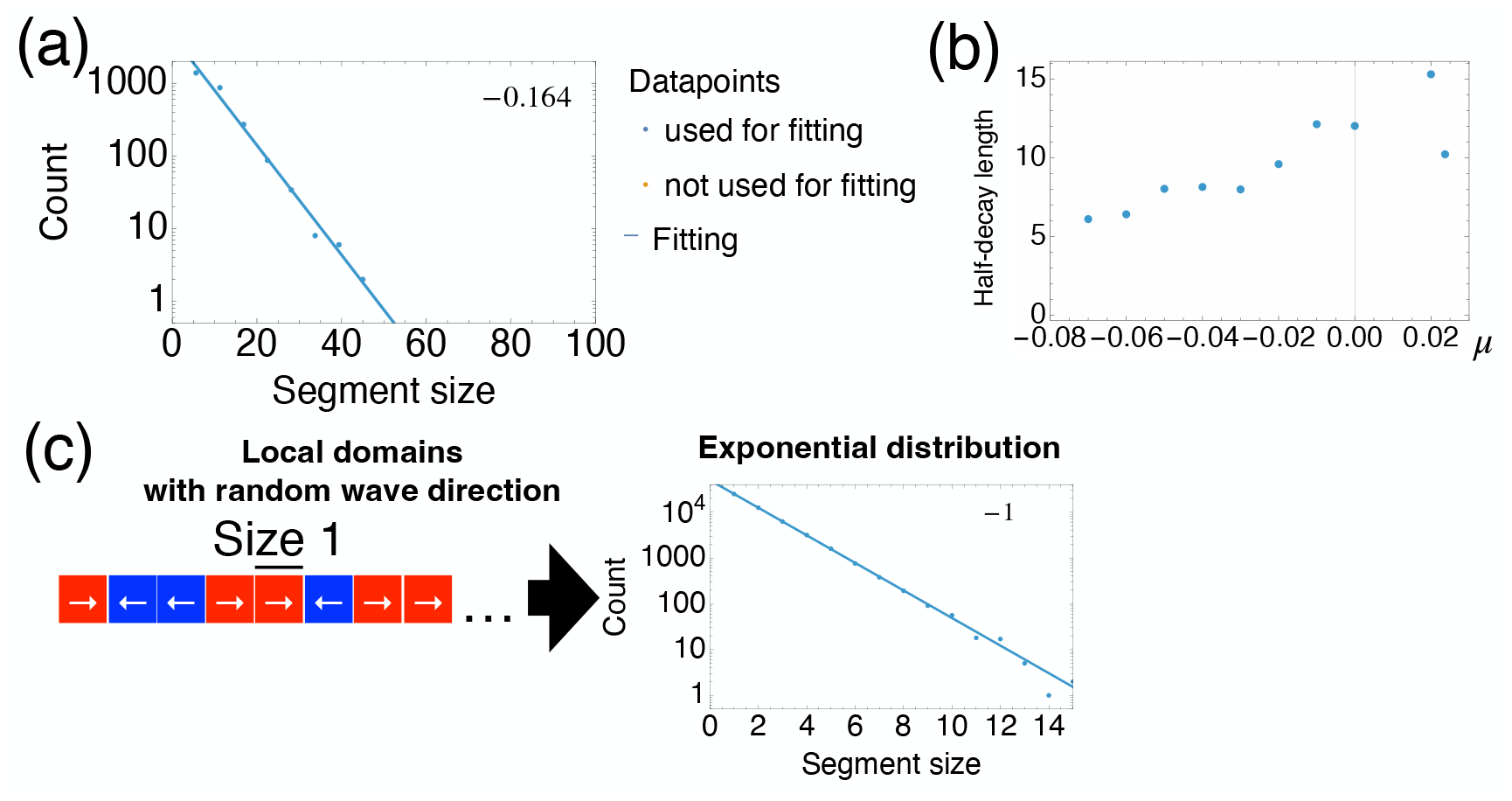
The exponential distribution of segment size implies the existence of minimum building blocks in the segmented pattern. (a) Segment size distribution at the biologically relevant timescale (*t* = 860) from white noise initial conditions. The numbers in the upper right corner represent the slope of the fitted line. (b) Estimated half-lengths for segment size frequency for different *µ*. The selection of data fitting points was based on the corrected Akaike information criterion (AICc). (c) Building blocks whose wave direction is randomly determined can explain the exponential segment size distribution.

We also examined the effect of timescale. At a very long timescale (*t* ∼ 1.4 × 10^6^), the system tended to exhibit fewer but larger segments (Supplementary Fig. 3a). This temporal difference was emphasized near the bifurcation as the segmented pattern disappeared for *µ* = 0.0237 (Supplementary Fig. 3a right). At a shorter timescale, it showed a large number of smaller segments (Supplementary Fig. 3b, c). The segment size appeared to follow the exponential distribution regardless of the timescale. The difference across timescales would again reflect the merging of initially emerged segments over the time course, as observed in Fig. 6.

The distribution implied the existence of a building block of a minimum segment length. Suppose the wave direction is determined locally, dependent on the initial white noise; that of each building block will be randomly determined, and thus, the segment size distribution will obey the exponential distribution (Fig. 8c).

#### 3.2.4. Interactions of segment boundaries

Motivated by the implied existence of locally emerged minimum building blocks of wavetrain segments, we next observed the interactions between segments with the specified segment length. To design the initial condition for the subsequent numerical simulations, we first estimated the amplitude and phase of the pure wavetrain pattern for the most unstable mode using weakly nonlinear analysis. Assume that *u, v*, and *w* have the following form,

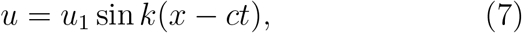

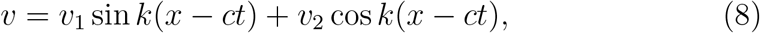

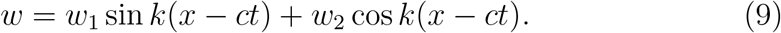

By calculating the difference of lhs and rhs in Eq. (1) and considering the corresponding wave components, we obtain,

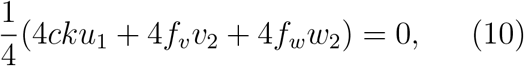

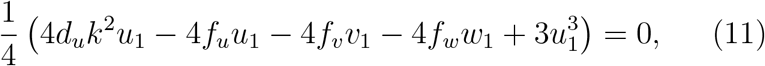

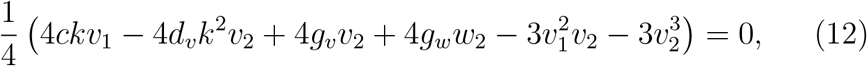

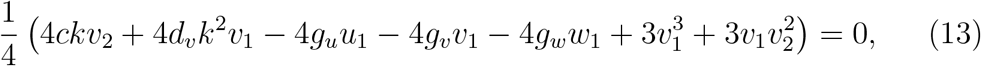

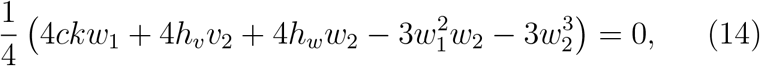

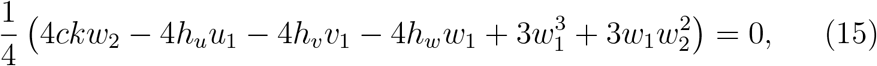

where

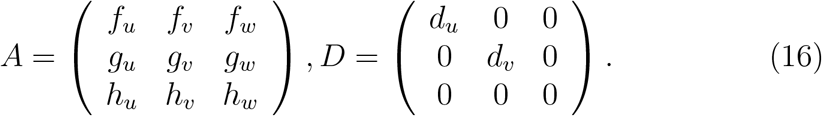

Then, we numerically solved Eqs. (10)-(15) for *c, u*_1_, *v*_1_, *v*_2_, *w*_1_, and *w*_2_, using *Mathematica* (Wolfram Research, Inc).

We performed numerical simulations for a certain period (1.4 × 10^3^) to ensure convergence, using the aforementioned solution as the initial condition. Then, we flipped the solution in *x* direction for an interval of *L*_1_ to obtain the initial condition for segment interaction observation (Fig. 9a).

**Figure 9:**
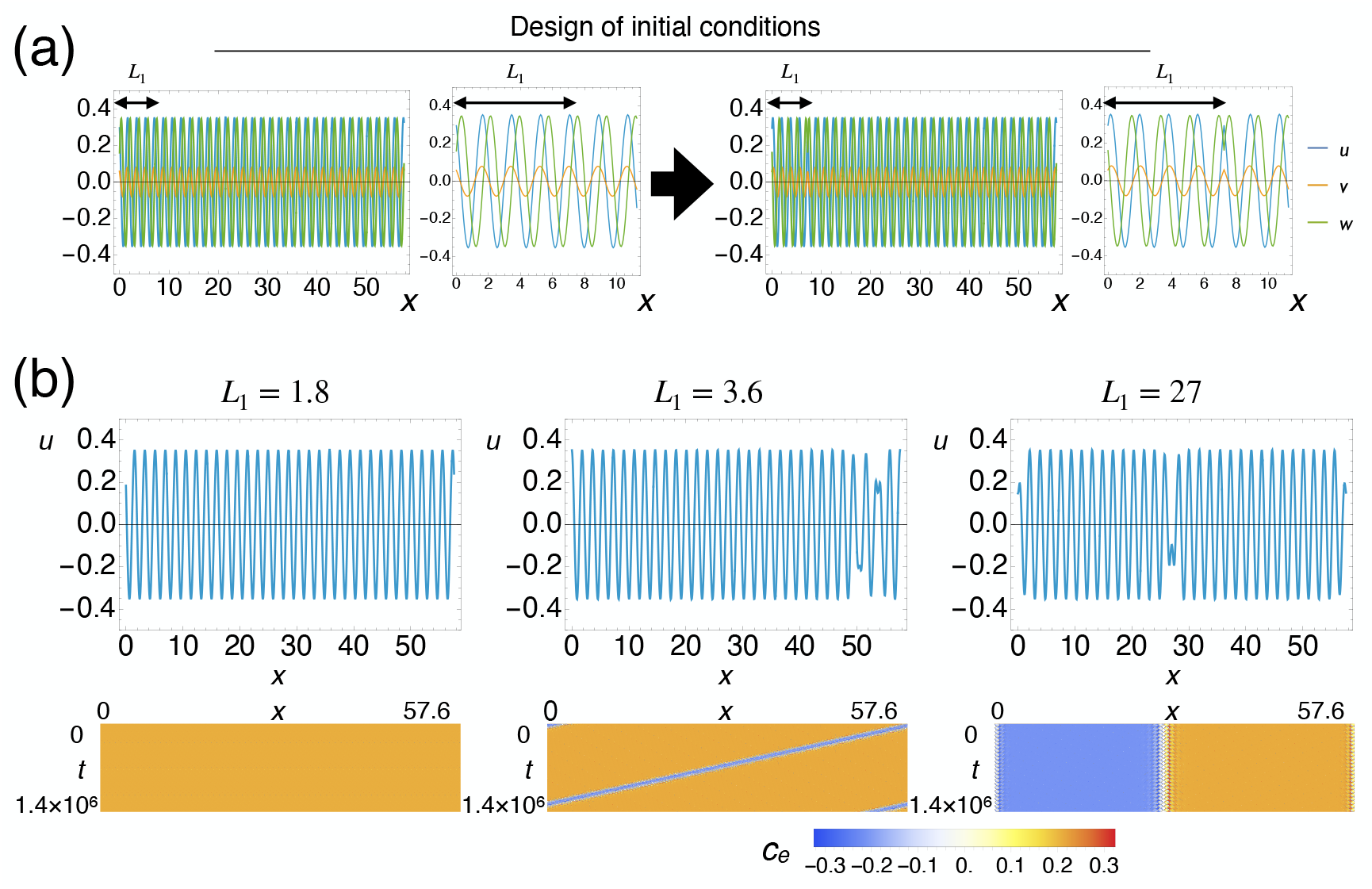
Segments shorter than a certain length cannot stably exist. (a) Schematic representation of initial condition construction. Interactions between segments were observed by numerical simulations with the constructed initial conditions. The values of *u, v*, and *w* for the interval of *L*_1_ were flipped in *x* direction. Left: the whole domain. Right: zoom-up. (b) Upper: plots of *u* at *t* ∼ 1.4 × 10^6^. Lower: color-coded *x* − *t* plots of the local wave velocity *c*_*e*_.

We changed *L*_1_ and performed numerical simulations for an extended period (1.4 × 10^6^) to observe whether segmented patterns were stably observed (Fig. 9b). When a flipped segment was relatively short, the segment was stably observed, but it moved at a constant speed (Fig. 9b center, Supplementary Movie 7). The flipped segment instantly vanished when the segment was too short (Fig. 9b left). The movement was slower when the flipped segment length was longer (Fig. 9b right, Supplementary Movie 8). We speculate that the moving segment boundary was observed because the system allows various wavenumber segments to exist in a periodic boundary domain, resulting in a very slow movement of the segments. We note that this movement is too slow to be detectable at a biological timescale.

We also explored the cases of different *µ*. In the previous model (Kawamura et al., 2021), which showed weaker instability, when a flipped segment was relatively short, it gradually shortened and eventually vanished (Supplementary Fig. 4a left). However, when a flipped segment was longer than a certain length (∼ 12.6), the segmented pattern was stably observed (Supplementary Fig. 4a center and right, Supplementary Movie 9). Then, with a system near bifurcation, the flipped segment did not stably exist, even if its length was 15-wavelength long (Supplementary Fig. 4b). We also expanded this analysis to a wider variety of *µ* and *L*_1_ and confirmed this trend (Fig. 4c).

Taken together, these observations indicated that short segments cannot stably exist within a finite-length domain, and their threshold depends on how far the system is from wave bifurcation. This is also consistent with the possible existence of building blocks and their implied size differences among the systems, as discussed in Section 3.2.3.

## 4 Discussion

### 4.1. Research summary

In this study, we revealed that the segmented pattern can frequently emerge at the biologically relevant spatiotemporal scale in the modified model. However, an additional mechanism was required for wave directional bias near the rete testis. Among the hypotheses tested, this bias is best explained by tubule elongation. Furthermore, large-scale numerical simulations have demonstrated that the frequency of segmented wavetrain formation depends on the domain size. The segment sizes follow an exponential distribution, and the stable segment size has a lower limit, implying that the interaction of locally emerged unidirectional wavetrain segments generates stable segmented patterns.

### 4.2. Elongation of the seminiferous tubules

In this study, we identified the tubule elongation as a possible contributor to inward-propagating spermatogenic waves observed in the mouse testis. However, the biological details of seminiferous tubule elongation remain unclear.

To the best of our knowledge, no reports have directly examined whether tubule elongation is spatially uniform or heterogeneous. A previous report indicates that the number of Sertoli cells ceases to increase around two weeks after birth, after which their nuclear size gradually increases (Auharek and de França, 2010), implying that an increase in cell size rather than cell number is a major contributor. In contrast, recent studies in hamsters and rats demonstrated that a subset of Sertoli cells retains proliferative capacity in the transition region after early postnatal stages, which connects seminiferous tubules and the rete testis, unlike in the convoluted seminiferous tubules (Aiyama et al., 2015; Figueiredo et al., 2016). Therefore, the tubules might elongate preferentially near the rete testis, which we incorporated as a hypothesis in our numerical exploration.

However, because this region contains a specialized valve-like structure called the Sertoli valve (Aiyama et al., 2015; Nagasawa et al., 2018), their proliferation may instead contribute to local structural remodeling rather than tubule elongation. The spatiotemporal observation of the size and proliferation of Sertoli cells, as well as their experimental manipulation, would provide valuable insights into their role in influencing the directional bias of wave propagation.

### 4.3. Mechanistic interpretation of inward wave generation by domain growth

It remains unclear how domain growth leads to the directional bias from a mathematical perspective, especially for uniform elongation. Although we have not yet obtained a rigorous mathematical characterization, we can provide additional numerical evidence for preferential domain growth.

For simplicity, we consider increasing the domain length instantaneously while keeping the newly added domains spatially uniform. The newly added domains serve as the source of perturbation. When the pattern propagates to the boundary, high-wavenumber components in the extended domain first approach the zero solution, and then the characteristic wavenumber components of the wavetrain grow due to wave instability (Supplementary Movie 10). The perturbation may first propagate to the boundary side and then reflect at the boundary, forming an inwardly moving wavetrain segment. On the other hand, when the pattern moves away from the boundary, the extended domain will be incorporated into the same segment (Supplementary Movie 11). Similar phenomena may underlie the direction bias observed in Section 3.1.3.

While this interpretation is based on numerical simulations, it is also consistent with known properties of boundary contact defects in reaction-diffusion systems with Neumann boundary conditions (Sandstede and Scheel, 2004). The reason why a moderate directional bias can be observed under uniform elongation remains unclear. However, since the spatial pattern size is continuously modulated by elongation, small perturbations similar to the cases of preferential elongation may also be effectively introduced near the boundaries.

### 4.4. Mathematical interpretation of segmented patterns

Our exploration implied that the segmented patterns can be understood as emerging from the interaction among locally emerged unidirectional wavetrain segments. In this study, we employed a numerical approach to investigate the segmented patterns and their biological relevance. Although the mathematically rigorous analysis of the pattern is not yet available, the observed boundaries between segments can be qualitatively interpreted as the interfaces between two stable wavetrain domains.

In the reaction-diffusion system, such interfaces are referred to as “defects” (Sandstede and Scheel, 2004). Specifically, the segmented patterns in our model can be viewed as sequences of source and sink defects, which separate wavetrains propagating in opposite directions. However, although the case for large but bounded domains is also discussed (see p.45 in Sandstede and Scheel (2004)), existing theories are primarily developed for a single interface in an infinite domain and cannot handle complex phenomena that involve multiple interacting defects in finite regions.

Importantly, we emphasize that our results are primarily based on numerical simulations inspired by biologically observed patterns and do not rely on prior theoretical results. The segmented patterns we investigate here emerge in a finite and sometimes small domain, especially when using biologically motivated boundary conditions, making them analytically challenging. In such settings, numerical simulation is an essential tool for revealing its structure and creating hypotheses for further theoretical investigation. This approach to complex nonlinear phenomena is widely used in the literature (Mimura, 2013; Datseris and Parlitz, 2022).

The concept of defects would serve as a helpful conceptual reference for understanding the mathematical structure of the patterns we have examined. Taken together, future theoretical developments will be necessary for a rigorous mathematical treatment of the numerically observed phenomena in this study.

### 4.5. Physiological implications

The physiological meanings of the cellular association pattern and the sites of reversal are unclear. If the primary function is to maintain the constant rate of sperm production, a random distribution would be sufficient, as previously discussed (Kawamura et al., 2021). However, when the sperm use some diffusive signals to regulate their differentiation cycle, local synchronization would occur. The wavetrain pattern may have evolved to avoid the periodic production and enable the constant production rate, while a diffusive signal is necessary for their differentiation.

Whether and how the site of reversal affects the sperm production function remains completely unknown. Our model predicts that the tubule elongation frequently produces at least one reversal site in a single seminiferous tubule. Further study is required to understand the physiological aspects of this pattern formation in seminiferous tubules.

### 4.6. Wavetrain patterns in broader biological contexts

Besides non-biological examples such as water waves or electrochemical oscillations (Fukushima et al., 2008), wavetrain patterns have been reported in a few biological systems. One example is the actin filament dynamics in the cell cortex (Michaud et al., 2022). In this study, *Xenopus* oocytes overexpressing Rho GEF Ect2 and Rho GAP RGA-3/4 showed traveling wavetrains. Another example is the accumulation patterns of myxobacteria, which exhibit two overlapping wavetrains with different directions (Welch and Kaiser, 2001; Igoshin et al., 2001). Cell tracking and genetic data revealed that their cells at early fruiting body stages formed wavetrain patterns based on the repulsion of neighboring cell populations driven by cell contact-mediated signaling. Our research contributes to the understanding of wavetrain patterns observed in living organisms.

### 4.7. Limitations and future perspectives

In our two studies (this and Kawamura et al. (2021)), we utilized threespecies reaction-diffusion equations as one of the simplest possible models. We adopted this framework because the complete picture of the molecular controls of sperm differentiation and their cellular association patterns remains largely unknown, except for diffusive retinoic acid signaling (Sugimoto et al., 2012). Therefore, this model is not intended to describe specific signaling networks.

Although we also explored the effect of different signaling activities near the rete testis in Appendix A.4, these simulations were intended to explore their possible effects abstractly. They should not be interpreted as direct predictions of the effects of the actual signaling pathways. Additionally, due to the limited quantitative data on actual wave dynamics, we were not able to perform a quantitative comparison. Quantitative large-scale data on actual wave dynamics would be a valuable future direction for understanding biological mechanisms and validating mathematical models. Further experimental research would enable us to develop more detailed mathematical models that can quantitatively connect the actual biochemical networks with the spermatogenic wave dynamics *in vivo*.

## Appendix A. Biological features that were found not to affect the directionality of spermatogenic wave progression

In this Appendix, we discussed the analyses of the possible mechanisms other than tubule elongation.

### Appendix A.1. Expansion of sperm-generating regions

Here, we considered the effect of expanding the spermatogenic region during development. At the initial stage of sexual maturation, spermatogenesis preferentially occurs at the region near the rete testis, but everywhere in the adult (Nakata et al., 2017). To simplify this situation, we performed numerical simulations with two very short domains (*L* = 0.225). We then expanded the domain by adding lattices of zero value on one boundary while maintaining zero-flux boundary conditions (Fig. A.1a). Finally, these two domains merged and became a single domain. We assumed that this expansion occurs before sexual maturation and thus merged at *t* = 42 (6 weeks old) within a biologically comparable domain size of *L* = 14.4. In this case, two segments appeared. Both sides of the tube robustly exhibited a wavetrain segment with an outward direction, regardless of the initial white noise (Fig. A.1b).

A different implementation method was also examined. Here, we defined the reaction term with spatiotemporal dependence using the control parameter *µ*,

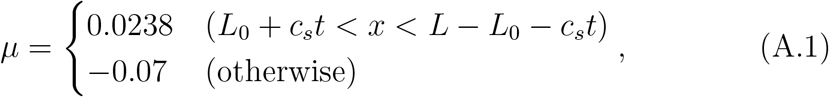

to represent a shrinking non-spermatogenic domain (Fig. A.1c). *L*_0_ and *c*_*s*_ are the initial spermatogenic domain length at each end and the expanding speed of spermatogenic regions, respectively. With zero-flux boundary conditions and biologically relevant expansion speed *c*_*s*_ ∼ 3.2 similar to the former case, wavetrains formed near both boundaries initially, and then propagated to the inner part (Fig. A.1d). The resulting pattern was two segments with outward directions, consistent with the observation in Fig. A.1b.

Taken together, the expansion of spermatogenic regions within a seminiferous tubule can determine the wave direction near the rete testis, but the direction would be the opposite to that observed *in vivo*.

**Figure A.1:**
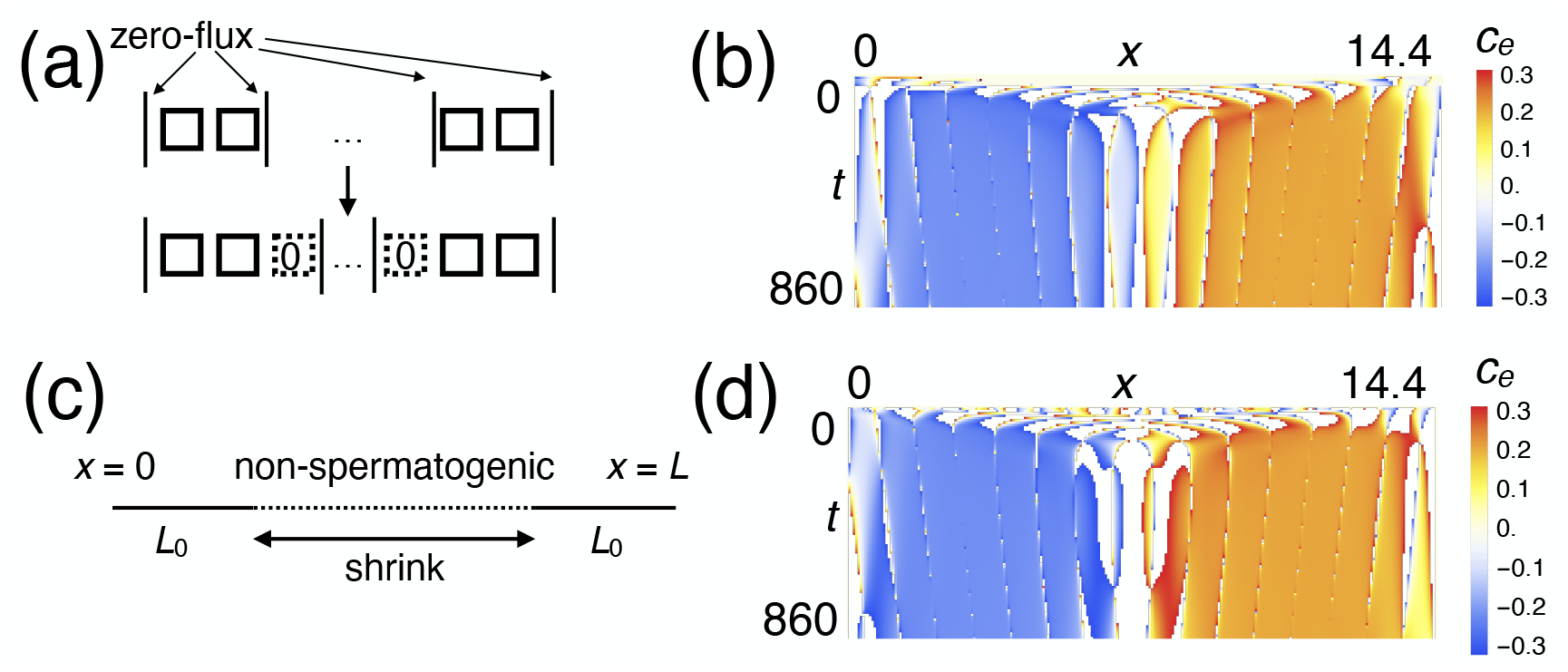
Effect of expanding spermatogenic region during development. (a) Schematic representation of the implementation of the additive domain growth. (b) *x* − *t* plot of local wave velocity *c*_*e*_. (c) Schematic representation of the implementation of the shrinking non-spermatogenic regions using different reaction terms. (d) *x* − *t* plot of local wave velocity *c*_*e*_.

### Appendix A.2. Flow inside the seminiferous tubules

Next, we examined the effect of flow inside the tubules. The luminal fluid moves along the seminiferous tubule to transport sperm from the testis into the epididymis. This motion is believed to involve peritubular myoid cells (Maekawa et al., 1996) or tunica albuginea (Middendorff et al., 2002). We assume the flow speed inside the seminiferous tubule is faster near the rete testis, as flow should reflect the sum of the excretion and pumping of the entire seminiferous tubule.

To incorporate the effect of luminal fluid flow, we added a space-dependent advection term for diffusive components *u* and *v* to the original model (Fig. A2a),

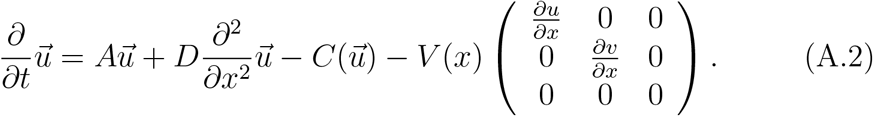

*V* (*x*) represents the flow speed at the position *x* and set as 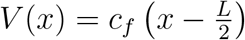 with *c*_*f*_ *>* 0, to express the flow towards the rete testis.

Segmented patterns were generated with amplitude deviation along *x* direction when *c*_*f*_ ∼ 0.0002 (Fig. A.2b). We performed multiple numerical simulations with different initial conditions of white noise. Around 80% of the simulations resulted in segmented patterns where two outermost segments have outward directions (Fig. A.2c). With a smaller *c*_*f*_, such a significant bias was not observed. At the same time, the frequency of inward direction at both ends, consistent with the actual wave direction, remains infrequent (Fig. A.2c). The flow can bias the segment wave direction at the edge. However, the bias reproduced in the model was inconsistent with the observation.

**Figure A.2:**
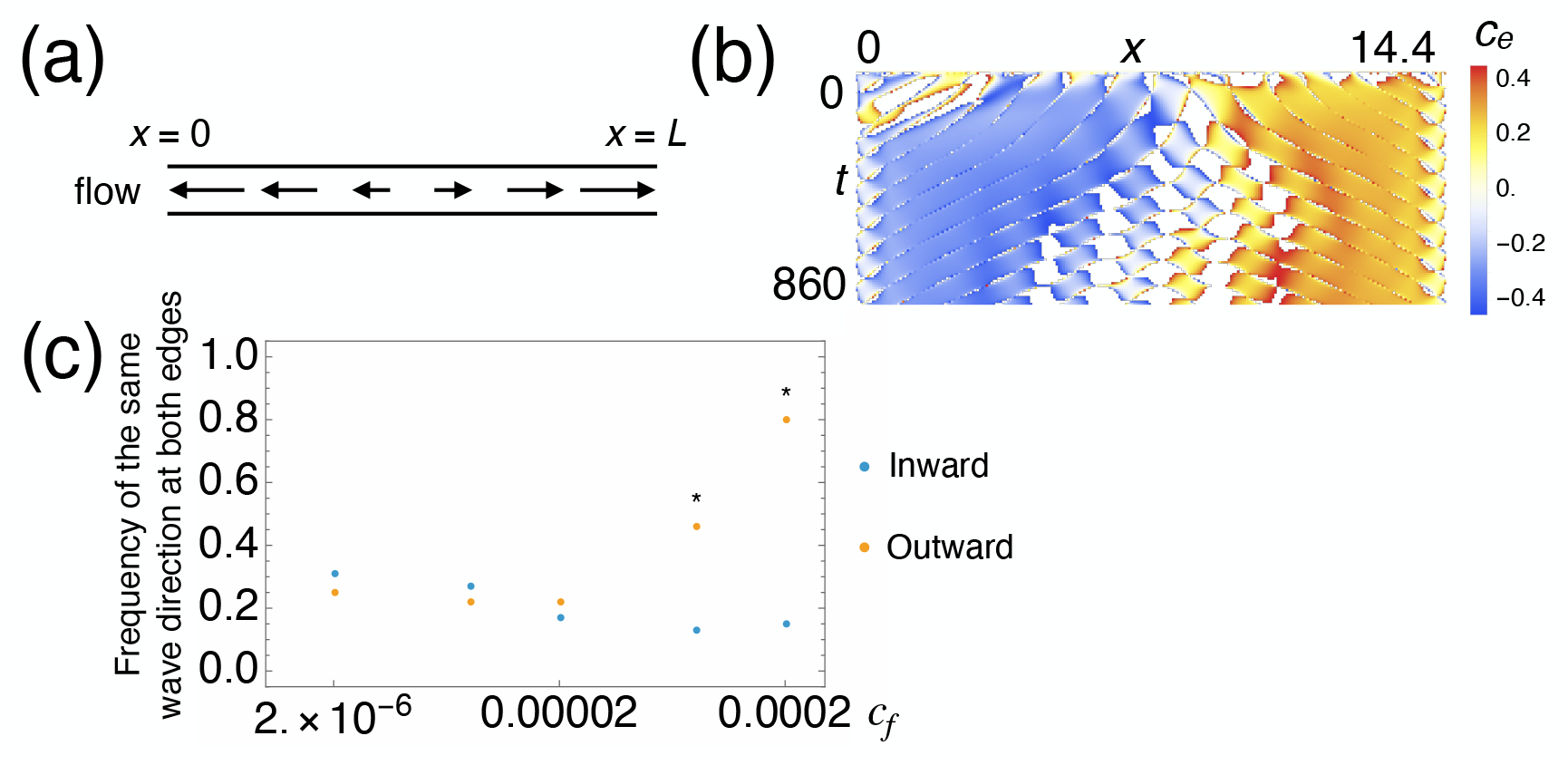
Effect of luminal flow within seminiferous tubules. (a) Schematic representation of the implementation of flow in the model. (b) *x* - *t* plot of local wave velocity *c*_*e*_ (*c*_*f*_ ∼ 0.0002). (c) The frequency of the same wave direction (inward or outward) at both edges for various flow gradients *c*_*f*_. ^*^: *p <* 1 × 10^-5^ (binomial test under the null hypothesis П = 0.25; two-sided; *n* = 100).

### Appendix A.3. Diameter change of seminiferous tubules around the rete testis

Intermediate regions and straight seminiferous tubules connect convoluted seminiferous tubules and the rete testis (Fig. A.3a, Nakata et al. (2015)). Intermediate regions are located just proximal to convoluted seminiferous tubules. They have Sertoli-like epithelial cells but no spermatogenic cells. Straight seminiferous tubules exist between the intermediate regions and the rete testis. They form an epithelial structure similar to the rete testis and are considerably thinner than convoluted ones. We wondered if such a diameter change could affect the wave direction.

Assume a tubular structure has a diameter *r*(*x*) at a position *x*. We can parameterize a tubular surface as (*x, r* cos *θ, r* sin *θ*). Diffusion can be expressed as Laplacian, Δ_*s*_ = ∇ · (∇ − **n**(**n** · ∇)), where **n** is a normal vector. Here, we consider a part of a cone surface whose slope is *c*_cone_ for simplicity. The surface can be parameterized as *r*(*x*) = *c*_cone_*x*+*r*_0_. The effect of diffusion in a longitudinal direction is given as

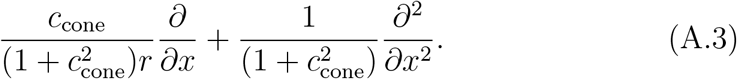

We simulated this effect using a one-dimensional domain by substituting the diffusion terms in Eq. (1) with Eq. (A.3). We considered a truncated cone with a ratio of the smallest and largest radii of 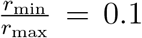. Because the typical diameter of convoluted seminiferous tubules is 200 *µ*m, we set the maximum radius as *r*_max_ = 0.1. The interface between segments moved to the minus direction, especially near the thinner boundary, and a segment of inward wave direction existed there with high frequency (Fig. A.3b). We performed multiple numerical simulations with different radius ratios. The probability of observing such a segment was significantly higher than random if the ratio is small enough (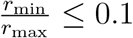, Supplementary Fig. A.3c).

The wave direction at the boundary is consistent with the observation in these numerical simulations. However, the diameter-changing intermediate regions are not necessarily long enough to affect the pattern dynamics *in vivo*. We performed a morphological measurement in a micrograph of a testis crosssection in a published report (Nakata, 2016). The diameters of convoluted and straight seminiferous tubules were approximately 200 *µ*m and 50 *µ*m, respectively. The intermediate region length was also around 200 *µ*m. The slope of a cone was higher than expected, but the diameter-changing regions were short.

Based on this information, we set the cone-cylinder-cone-like structure for numerical simulations. Cylinder part length was set as eight times the characteristic pattern size (*L* = 14.4), with two very short truncated cones at both ends (*L*_cone_ = 0.2025, *r*_min_ = 0.025, *r*_max_ = 0.1, *c*_cone_ ∼ 0.37). In this setting, we did not observe any bias in the wave direction at both edges for 100 simulations (Fig. A.3d,e). In the realistic setting, the diameter change may not affect the wave direction within the convoluted seminiferous tubules.

**Figure A.3:**
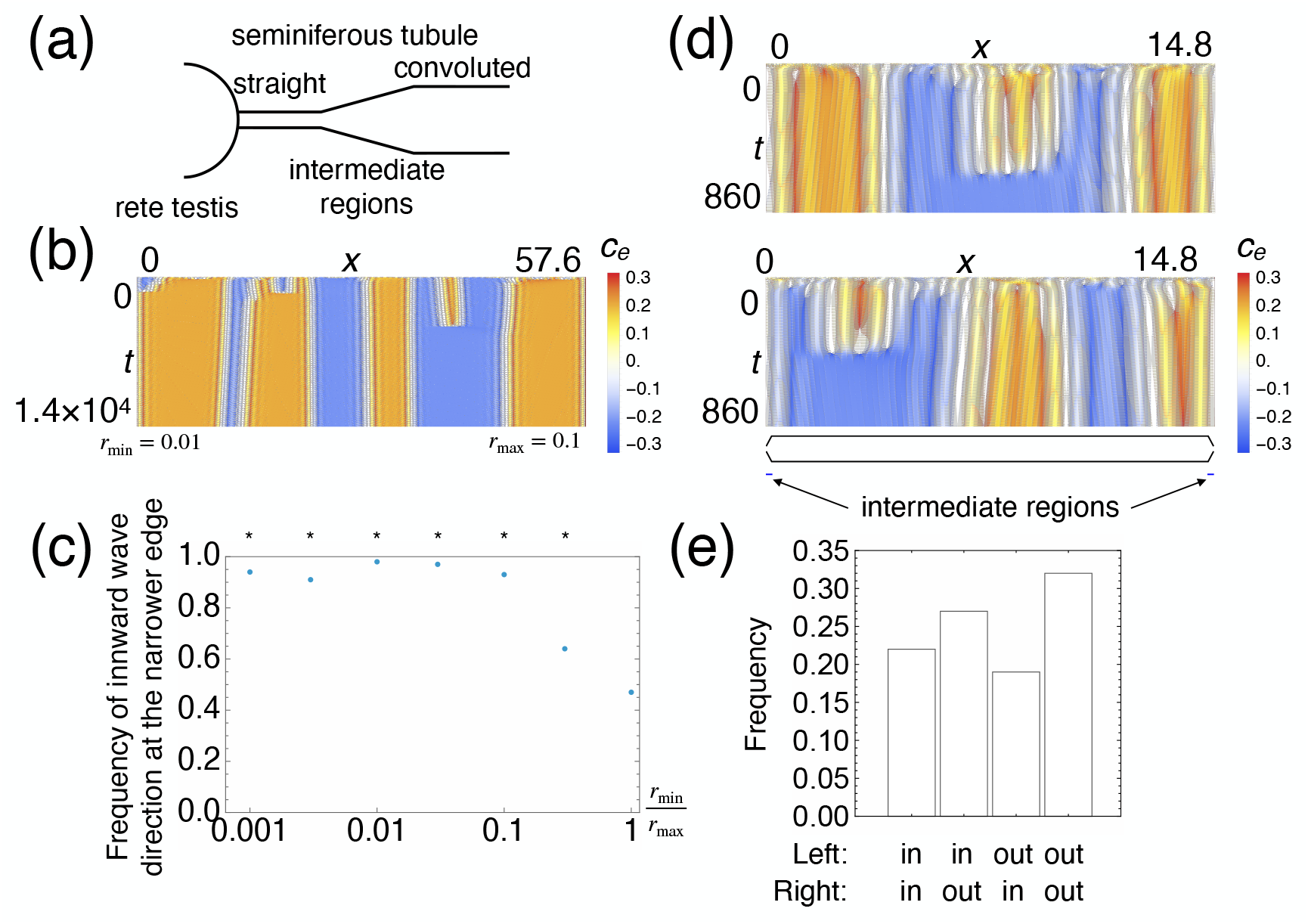
Effect of diameter change near the rete testis. (a) Schematic representation of diameter changes around the rete testis. (b) *x* − *t* plot of local wave velocity *c*_*e*_ on a one-dimensionally projected truncated cone surface. (c) The effect of the minimum diameter on the wave direction bias at the thinner edge. ^*^: *p <* 0.01 (binomial test under the null hypothesis П = 0.5; one-sided; *n* = 100). (d) Numerical simulation results shown as *c*_*e*_ in a realistic geometry. (e) The frequency of wave direction patterns at both edges in a realistic geometry. No significant bias was observed (*p* = 0.27; Pearson’s *χ*^2^ test under the null hypothesis of equal probability; *n* = 100).

### Appendix A.4. Distinct signaling activity in the rete testis

The rete testis and the transition region between the rete testis and seminiferous tubules are known to have distinct signaling activity from the seminiferous tubules. Recent studies show that the transitional zone between the seminiferous tubules and rete testis exhibits a specialized activity, including STAT3, retinoic acid, and FGF (Nagasawa et al., 2018; Imura-Kishi et al., 2021). Another study shows that the retinoic acid produced in the rete testis can re-activate the spermatogenesis in the mice with Sertoli cellspecific deletion of retinoic acid-synthesizing enzymes (Teletin et al., 2019). These findings lead us to consider whether any distinctive signaling state can dictate the wave propagation direction near the rete testis.

To examine this, we considered two scenarios. One is that the signaling activity is maintained distinctively in the rete testis or the transition region, and this affects the seminiferous tubules via a diffusive signaling molecule. This was implemented by introducing Dirichlet conditions for diffusive variables *u* and *v*, where *u*|_*x*=0_ = *u*|_*x*=*L*_ = *u*_boundary_ and *v*|_*x*=0_ = *v*|_*x*=*L*_ = *v*_boundary_ (Fig. A.4a). Another is that diffusive signaling molecules are constantly produced in the rete testis or the transition region, and this affects the seminiferous tubules. This was implemented by introducing constant source terms only in the boundary lattices:

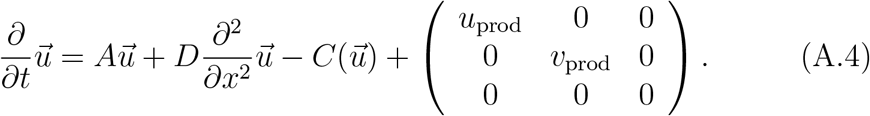

We performed the numerical simulations from random initial noise with various boundary values or production rates for *u* and *v*. In both implementation methods, we did not observe any significant dependency of inward propagating bias at both edges on them (Fig. A.4c,d). The frequency of such bias emergence was distributed 20-40% for both implementation methods (Fig. A.4e,f). We confirmed that these modifications indeed altered the behaviors near the boundary, as indicated by the large deviations observed (Fig. A.4g-j).

Because our model is extremely simplified and phenomenological, the correspondence with the actual molecular activity is not clear. Although other ways of implementing distinct signaling activities could contribute to the consistent bias in the wave direction, our exploration based on the simple assumptions did not identify any numerical evidence.

**Figure A.4:**
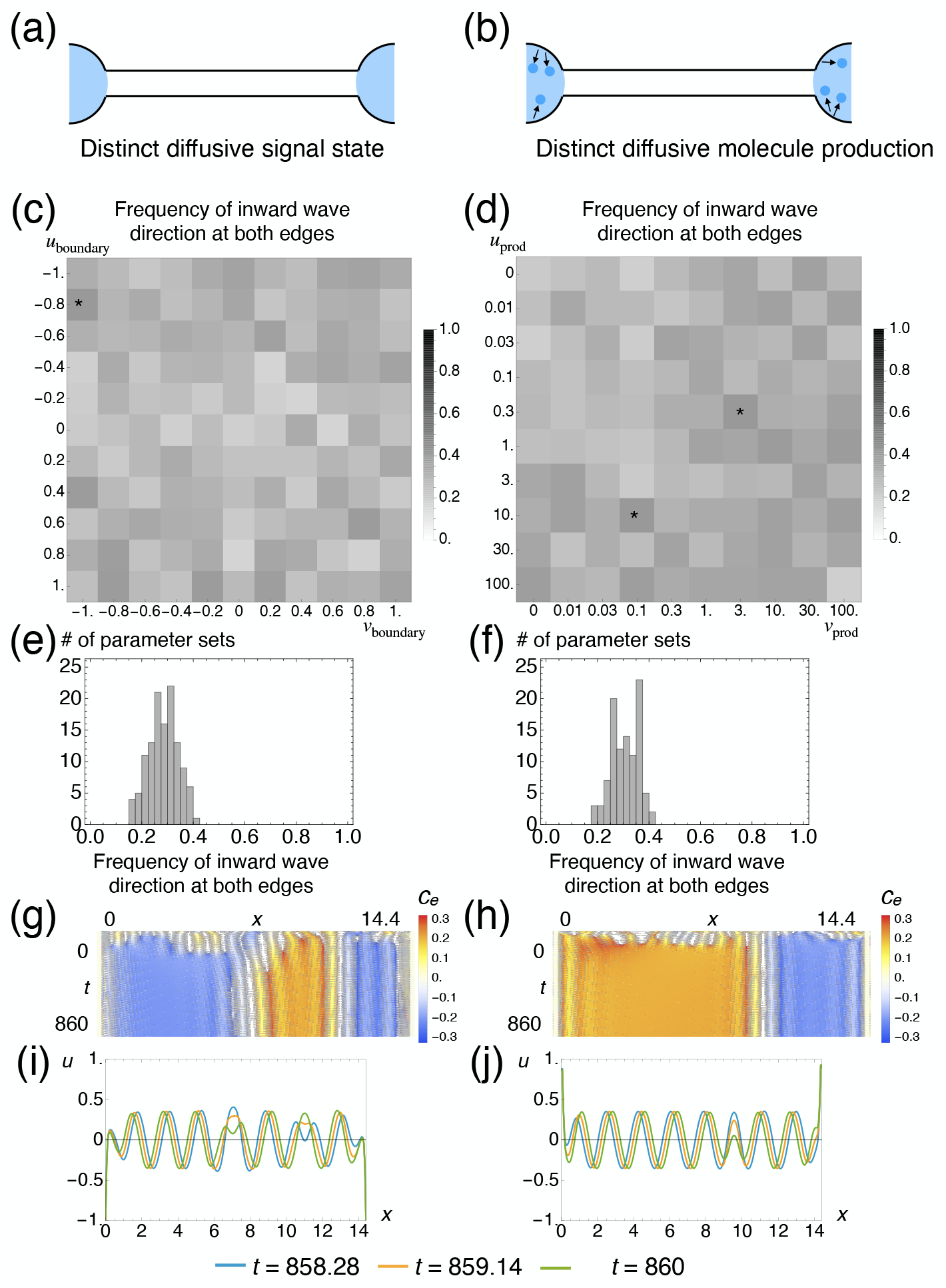
Effect of distinctive signaling activity near the rete testis. (a,b) Schematic representation of two implementation methods. The signaling state of the rete testis or the transition region is distinctively maintained (a,c,e,g,i). The diffusive signaling molecules are constantly produced at the rete testis or the transition region (b,d,f,h,j). (c,d) Frequencies of inward propagating wave emergence at both edges for various conditions at *t* = 860. 100 independent random white noises were given as the initial conditions. ^*^: Highest frequency (0.4). (e,f) The distributions of frequency of inward propagating wave emergence at both edges for various conditions. (g,h) *x* − *t* plots of local wave velocity *c*_*e*_ for a condition showing the highest frequency of inward propagating wave emergence at both edges. (i,j) *x* − *u* plots for three different time points, corresponding to (g,h).

## Supplementary material

- Supplementary Figures 1-4.
- Supplementary Movie 1. The animation of numerical simulation results corresponding to Fig. 3b,d.
- Supplementary Movie 2. The animation of numerical simulation results corresponding to Supplementary Fig. 1a,c.
- Supplementary Movie 3. The animation of numerical simulation results corresponding to Fig. 4.
- Supplementary Movie 4. The animation of numerical simulation results corresponding to Fig. 5c upper, e.
- Supplementary Movie 5. The animation of numerical simulation results corresponding to Fig. 5d upper, f.
- Supplementary Movie 6. The animation of numerical simulation results corresponding to Fig. 6.
- Supplementary Movie 7. The animation of numerical simulation results corresponding to Fig. 9b center.
- Supplementary Movie 8. The animation of numerical simulation results corresponding to Fig. 9b right.
- Supplementary Movie 9. The animation of numerical simulation results corresponding to Supplementary Fig. 4a right.
- Supplementary Movie 10. The animation of numerical simulation results that illustrates the emergence of a segmented pattern when the original wave propagates toward a boundary (outward) before the domain is extended on its side.
- Supplementary Movie 11. The animation of numerical simulation results that illustrates the absence of a segmented pattern when the original wave propagates away from a boundary (inward) before the domain is extended on its side.
- Supplementary File 1. Numerical simulation codes used in this study.

## Acknowledgments

The authors thank Dr Ryo Sugimoto for the original question about cellular association patterns and Professor Hiroki Nakata (Komatsu University), Professor Yoshiakira Kanai (University of Tokyo), and his lab members for helpful suggestions and comments.

## Declarations

### Data availability

The code used for the numerical simulations presented in this paper is available as supplementary material. Other data that support the findings of this study are available from the corresponding author upon reasonable request.

### Declaration of generative AI and AI-assisted technologies in the writing process

During the preparation of this work, the authors used Grammarly (Grammarly Inc.) and ChatGPT (OpenAI, Inc.) for language editing. After using these tools, the authors reviewed and edited the content as needed and take full responsibility for the content of the published article.

### Author contributions

Conceptualization: KS, TM. Data curation: KS, TM. Formal analysis: KS, AS. Funding acquisition: KS, AS. Investigation: KS, AS. Methodology: KS. Project administration: KS, TM. Resources: KS, TM. Software: KS. Supervision: TO, TM. Visualization: KS. Writing – original draft: KS, AS. Writing – review and editing: KS, AS, TO, TM.

