## Supplementary Figures for "Segmented wavetrains and sites of reversal in the mouse seminiferous tubules"

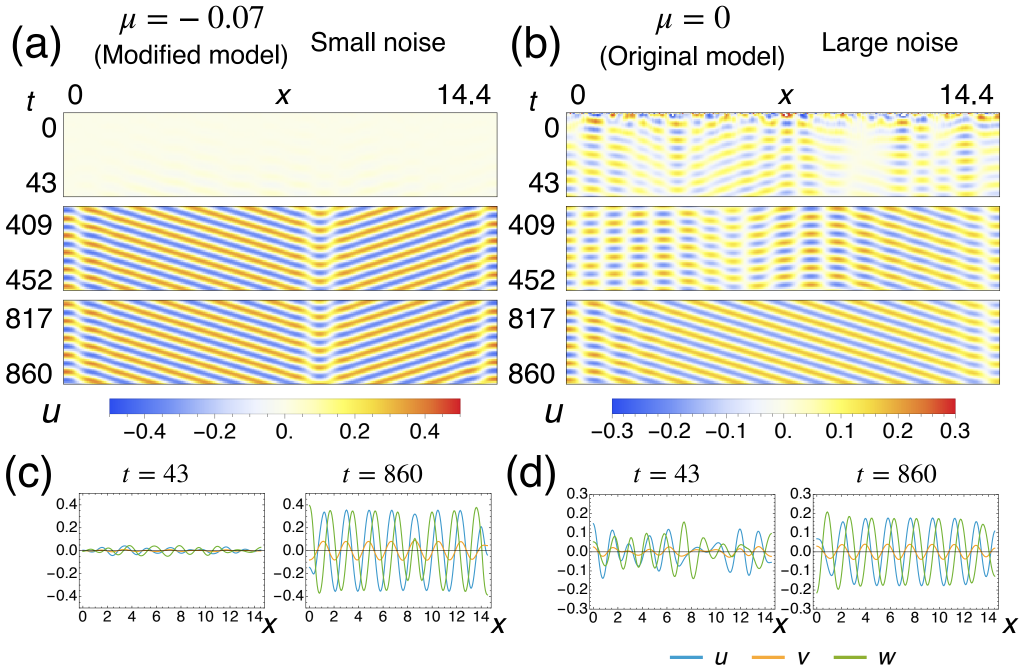


Supplementary Figure 1: Additional details for model modification for faster pattern formation that reflects the actual spatiotemporal scale *in vivo*. Representative numerical simulation results for $u$ in the modified model with the initial condition given by small-amplitude white noise (a,c) and the original model (Kawamura et al., 2021) with the initial condition given by larger-amplitude (50×) white noise (b,d). (a,b) $x-t$ plot with $u$ shown in color. Only initial, middle, and final time segments are visualized. (c,d) Representative snapshots for $u$, $v$, and $w$. Left: $t = 43$, corresponding to 7-week-old mice. Right: $t = 860$, corresponding to two-year-old mice.

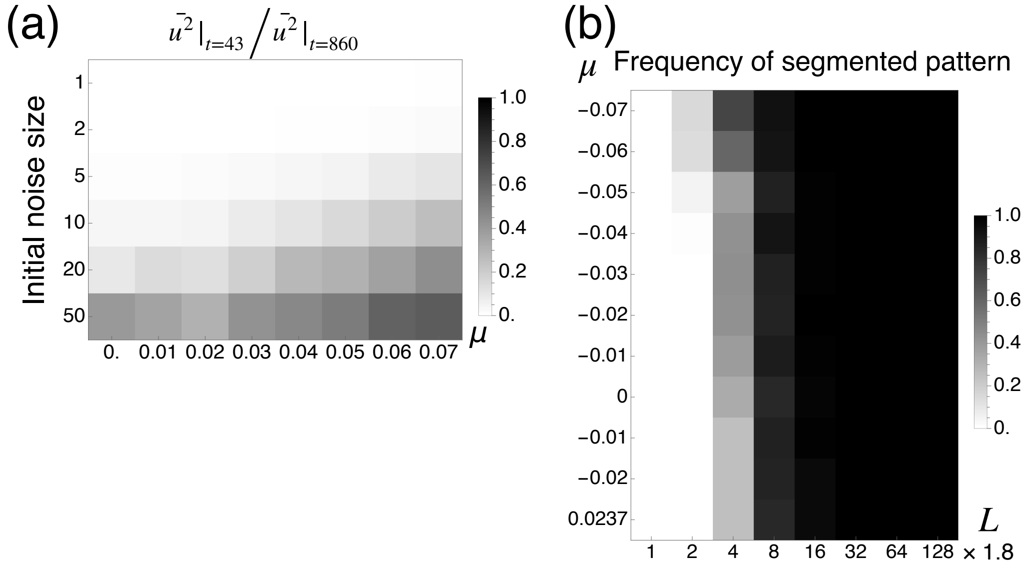


Supplementary Figure 2: Comparison for different $\mu$ values. (a) Normalized mean squared values of $u$ at $t = 43$ (vs. $t = 860$) for various $\mu$ and initial noise amplitude. (b) Frequency of segmented pattern emergence for various domain lengths $L$ and control parameters $\mu$ at $t = 860$. 100 independent random white noises were given as the initial conditions.

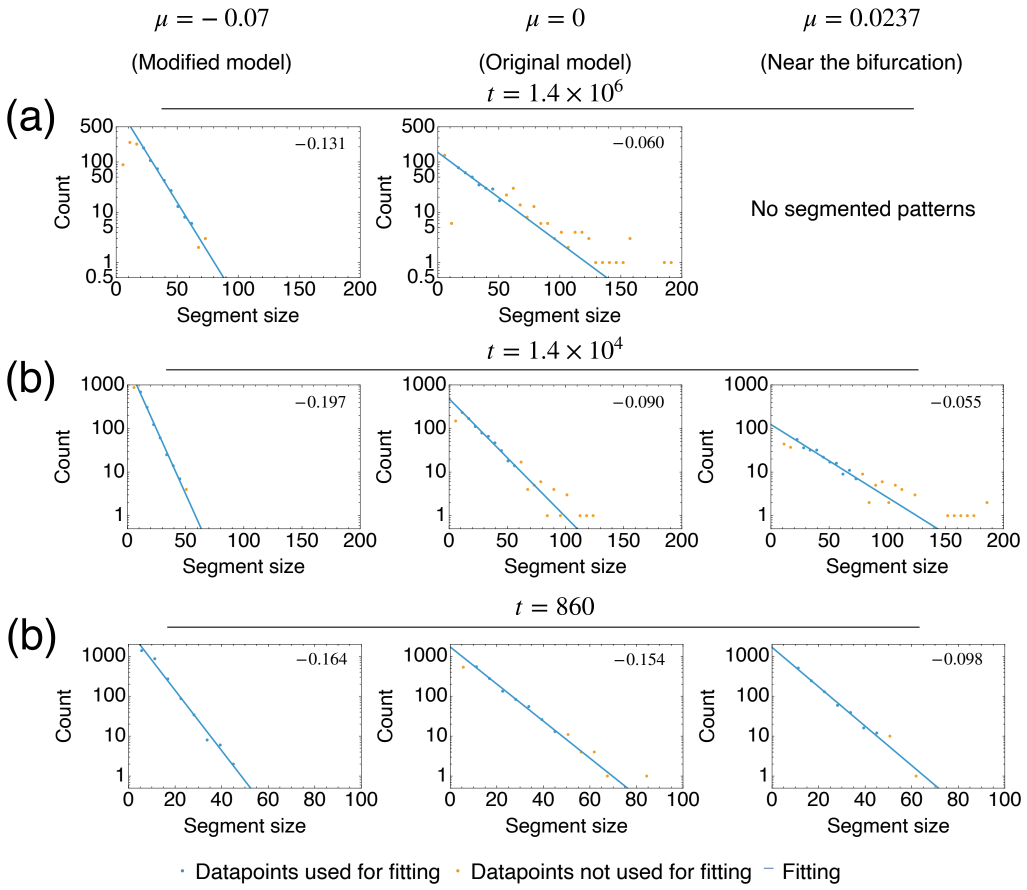


Supplementary Figure 3: Segment size distribution at different timescales. (a,b) Segment size distribution at $t \sim 1.4 \times 10^6$ (a), $t \sim 1.4 \times 10^4$ (b), and $t = 860$ (c), for long one-dimensional domains $L = 1843.2$ from white noise initial conditions. Left: Modified model in this paper ($\mu = -0.07$). Center: Original model ($\mu = 0$) (Kawamura et al., 2021). Right: A model with weaker instability ($\mu = 0.0237$). The histograms are shown for the combined data with ten different initial conditions as log-linear plots. The numbers in the upper right corner represent the slope of the fitted line.

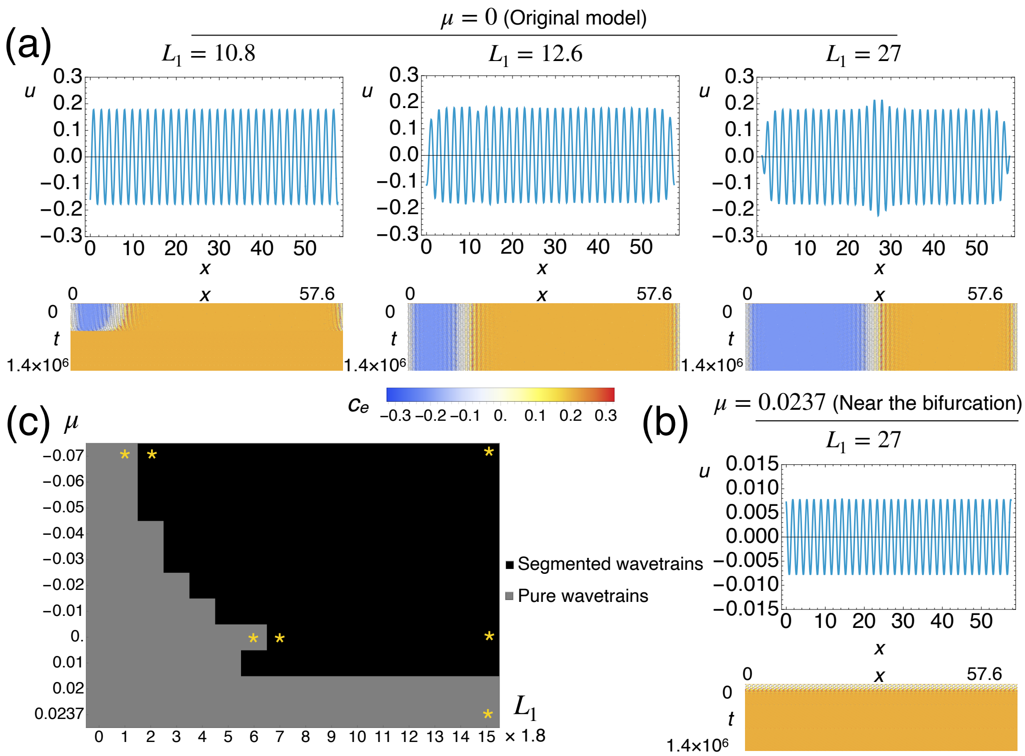


Supplementary Figure 4: Exploration of the numerical stability of short segments for different control parameter values. (a,b) Upper: plots of $u$ at $t \sim 1.4 \times 10^6$. Lower: color-coded $x - t$ plots of the local wave velocity $c_e$. (a) $\mu = 0$ (the original model in Kawamura et al. (2021)), (b) $\mu = 0.0237$ (near the bifurcation). (c) The pattern classification at $t \sim 1.4 \times 10^6$ for various $\mu$ and $L_1$. *: Parameter combinations shown in (a), (b), and Fig. 9.
