## Supplementary material for "Segmented wavetrains and sites of reversal in the mouse seminiferous tubules": SupplementaryFile.c

```
//Numerical Simulation Codes Used in This Study (C Language)
//The following C source codes were used for numerical simulations in this study.
//Attention: This single file contains multiple code segments arranged in order.
//Please extract the necessary parts before executing the code.
//The file MT.h (Mersenne Twister) is required and available at:　https://omitakahiro.github.io/random/code/MT.h
//***
//file1.c 	Numerical simulation using periodic boundary conditions.
//file2.c 	Numerical simulation using Neumann boundary conditions.
//file3.c 	Numerical simulation with uniformly growing domains. Corresponds to Fig. 5a,c,e
//file4.c 	Numerical simulation with preferentially growing domains. Corresponds to Fig. 5b,d,f.
//file5.c 	Numerical simulation using Neumann boundary conditions and predefined initial conditions. Corresponds to Fig. 9 and Fig. S3
//file6.c 	Numerical simulation using Neumann boundary conditions, where two domains expand inward and merge. Corresponds to Fig. A.1a,b.
//file7.c 	Numerical simulation using Neumann boundary conditions with multiple domains (expanding or shrinking) under different control parameters. Corresponds to Fig. A.1c,d.
//file8.c 	Numerical simulation including advection terms. Corresponds to Fig. A.2.
//file9.c 	Numerical simulation with varying diameter in a simplified setting. Corresponds to Fig. A.3b,c.
//file10.c 	Numerical simulation with varying diameter in a biologically realistic setting. Corresponds to Fig. A.3d,e.
//file11.c 	Numerical simulation with Dirichlet boundary conditions for u and v. Corresponds to Fig. A.4a,c,e,g,i.
//file12.c 	Numerical simulation with source terms at the boundaries. Corresponds to Fig. A.4b,d,f,h,j.
//***

//file1.c
//Numerical simiulation code with periodic boundary conditions.
//Note that in the codes, we used the different scales than in the manuscript.
//L = 3.2 and T = 6.2 in the codes corresponds to L = 1.8 and T = 8.6, respectively.

#include <stdio.h>
#include <stdlib.h>
#include <time.h>
#include "MT.h"// can be obtained from https://omitakahiro.github.io/random/code/MT.h

//Parallelization
//#ifdef _OPENMP
//#include <omp.h>
//#endif

#define _USE_MATH_DEFINES

#include <math.h>

#define stepsperout 1000000 //time steps per output
#define outsteps 1000 //number of outputs

double f(double u, double v, double w, double p){
  return 0.5 * u - v + w - u*u*u - p * u;
}

double g(double u, double v, double w, double p){
  return 1.5 * u - 2.1 * v - v*v*v - p * v;
}

double h(double u, double v, double w, double p){
  return - u - 0.1 * w - w*w*w - p * w;
}

int main(void) {
  //#pragma omp parallel
  //{
    int rep;
    //#pragma omp for
    for(rep=0;rep<1;rep=rep+1) {
      double dt = 0.001;//discrete time step
      int ln = 4096;//lattice number
      double dx = 0.1;//spatial lattice size
      double dx2inv = 1.0/dx/dx;
      double s = 1.1867777442770433;//spatial scaling paramter throughout the study
      double Du = 0.03 * s;//diffusion coefficient for u
      double Dv = 1.0 * s;//diffusion coefficient for v
      double p = -0.07;//control parameter \mu
      double r = 50.0;//white noise amplitude
      int n1,m,y;

      double *u, *v, *w, *uold, *pv;
      double *unew, *vnew, *wnew;
      u = (double*) malloc(sizeof(double) *  ln);
      v = (double*) malloc(sizeof(double) *  ln);
      w = (double*) malloc(sizeof(double) *  ln);
      unew = (double*) malloc(sizeof(double) *  ln);
      vnew = (double*) malloc(sizeof(double) *  ln);
      wnew = (double*) malloc(sizeof(double) *  ln);
      uold = (double*) malloc(sizeof(double) *  ln);
      pv = (double*) malloc(sizeof(double) *  ln);

      srand((unsigned)time(NULL)+rep*100);
      for (n1 = 0; n1 < ln; n1++){
        u[n1] = (0.02 * rand()/RAND_MAX -0.01)*r;
        v[n1] = (0.02 * rand()/RAND_MAX -0.01)*r;
        w[n1] = (0.02 * rand()/RAND_MAX -0.01)*r;
        pv[n1] = 0.0;
      }

      double k1u, k1v, k1w, k2u, k2v, k2w, k3u, k3v, k3w, k4u, k4v, k4w;
      n1 = 0;
      FILE *outputfile; 
      char filename[50];
      sprintf(filename, "turing_feedback-%03d.txt",rep);
      outputfile = fopen(filename, "w");
      while(n1 < outsteps) {
        for (y=0; y<ln; y++){
          fprintf(outputfile, "%g\t", (u[y])); 
        }
        for (y=0; y<ln; y++){
          fprintf(outputfile, "%g\t", (pv[y]));
        }
        for (y=0; y<ln; y++){
          fprintf(outputfile, "%g\t", (v[y])); 
        }
        for (y=0; y<ln; y++){
          fprintf(outputfile, "%g\t", (w[y])); 
        }
        fprintf(outputfile, "\n");
        
        n1++;
        for (m = 0; m < stepsperout; m++){
          for (y=0; y< ln; y++){
            k1u = f(u[y], v[y], w[y],p);
            k1v = g(u[y], v[y], w[y],p);
            k1w = h(u[y], v[y], w[y],p);
            k2u = f(u[y] + k1u * dt / 2, v[y] + k1v * dt / 2, w[y] + k1w * dt / 2,p);
            k2v = g(u[y] + k1u * dt / 2, v[y] + k1v * dt / 2, w[y] + k1w * dt / 2,p);
            k2w = h(u[y] + k1u * dt / 2, v[y] + k1v * dt / 2, w[y] + k1w * dt / 2,p);
            k3u = f(u[y] + k2u * dt / 2, v[y] + k2v * dt / 2, w[y] + k2w * dt / 2,p);
            k3v = g(u[y] + k2u * dt / 2, v[y] + k2v * dt / 2, w[y] + k2w * dt / 2,p);
            k3w = h(u[y] + k2u * dt / 2, v[y] + k2v * dt / 2, w[y] + k2w * dt / 2,p);
            k4u = f(u[y] + k3u * dt, v[y] + k3v * dt, w[y] + k3w * dt,p);
            k4v = g(u[y] + k3u * dt, v[y] + k3v * dt, w[y] + k3w * dt,p);
            k4w = h(u[y] + k3u * dt, v[y] + k3v * dt, w[y] + k3w * dt,p);
            unew[y] = u[y] + (k1u + 2 * k2u + 2 * k3u + k4u) / 6 * dt + Du * (u[(y+1)%ln]+u[(y-1+ln)%ln]-2*u[y])*dx2inv*dt;
            vnew[y] = v[y] + (k1v + 2 * k2v + 2 * k3v + k4v) / 6 * dt + Dv * (v[(y+1)%ln]+v[(y-1+ln)%ln]-2*v[y])*dx2inv*dt;
            wnew[y] = w[y] + (k1w + 2 * k2w + 2 * k3w + k4w) / 6 * dt;
          }
          for (y=0; y< ln; y++){
            uold[y] = u[y];
            u[y] = unew[y];
            v[y] = vnew[y];
            w[y] = wnew[y];          
          }
          for (y=0; y< ln; y++){
            pv[y] = -((u[y]-uold[y])/dt)/((u[(y+1)%ln]-u[(y-1+ln)%ln])/2.0/dx);
          }
        }
        printf("%d %d\n",rep,n1);
      }
      fclose(outputfile);
    }
  //}
  return 0;
}

//file2.c
//Numerical simiulation code with Neumann boundary conditions.
//Note that in the codes, we used the different scales than in the manuscript.
//L = 3.2 and T = 6.2 in the codes corresponds to L = 1.8 and T = 8.6, respectively.

#include <stdio.h>
#include <stdlib.h>
#include <time.h>
#include "MT.h" // can be obtained from https://omitakahiro.github.io/random/code/MT.h

//Parallelization
//#ifdef _OPENMP
//#include <omp.h>
//#endif

#define _USE_MATH_DEFINES

#include <math.h>

#define stepsperout 620 //time steps per output
#define outsteps 1000 //number of outputs

double f(double u, double v, double w, double p){
  return 0.5 * u - v + w - u*u*u - p * u;
}

double g(double u, double v, double w, double p){
  return 1.5 * u - 2.1 * v - v*v*v - p * v;
}

double h(double u, double v, double w, double p){
  return - u - 0.1 * w - w*w*w - p * w;
}

int main(void) {
  //#pragma omp parallel
  //{
    int rep;
    //#pragma omp for
    for(rep=0;rep<1;rep=rep+1) {
      double dt = 0.001;//discrete time step
      int ln = 32*8+2;//lattice number
      double dx = 0.1;//spatial lattice size
      double dx2inv = 1.0/dx/dx;
      double s = 1.1867777442770433;//spatial scaling paramter throughout the study
      double Du = 0.03 * s;//diffusion coefficient for u
      double Dv = 1.0 * s;//diffusion coefficient for v
      double p = -0.07;//control parameter \mu
      double r = 50.0;//white noise amplitude
      int n1,m,y;

      double *u, *v, *w, *uold, *pv;
      double *unew, *vnew, *wnew;
      u = (double*) malloc(sizeof(double) *  ln);
      v = (double*) malloc(sizeof(double) *  ln);
      w = (double*) malloc(sizeof(double) *  ln);
      unew = (double*) malloc(sizeof(double) *  ln);
      vnew = (double*) malloc(sizeof(double) *  ln);
      wnew = (double*) malloc(sizeof(double) *  ln);
      uold = (double*) malloc(sizeof(double) *  ln);
      pv = (double*) malloc(sizeof(double) *  ln);

      srand((unsigned)time(NULL)+rep*100);
      for (n1 = 0; n1 < ln; n1++){
        u[n1] = (0.02 * rand()/RAND_MAX -0.01)*r;
        v[n1] = (0.02 * rand()/RAND_MAX -0.01)*r;
        w[n1] = (0.02 * rand()/RAND_MAX -0.01)*r;
        pv[n1] = 0.0;
      }

      double k1u, k1v, k1w, k2u, k2v, k2w, k3u, k3v, k3w, k4u, k4v, k4w;
      n1 = 0;
      FILE *outputfile;
      char filename[50];
      sprintf(filename, "turing_feedback-%03d.txt",rep);
      outputfile = fopen(filename, "w");
      while(n1 < outsteps) {
        for (y=0; y<ln; y++){
          fprintf(outputfile, "%g\t", (u[y]));
        }
        for (y=0; y<ln; y++){
          fprintf(outputfile, "%g\t", (pv[y]));
        }
        for (y=0; y<ln; y++){
          fprintf(outputfile, "%g\t", (v[y]));
        }
        for (y=0; y<ln; y++){
          fprintf(outputfile, "%g\t", (w[y]));
        }
        fprintf(outputfile, "\n");

        n1++;
        for (m = 0; m < stepsperout; m++){
          for (y=0; y< ln; y++){
            k1u = f(u[y], v[y], w[y],p);
            k1v = g(u[y], v[y], w[y],p);
            k1w = h(u[y], v[y], w[y],p);
            k2u = f(u[y] + k1u * dt / 2, v[y] + k1v * dt / 2, w[y] + k1w * dt / 2,p);
            k2v = g(u[y] + k1u * dt / 2, v[y] + k1v * dt / 2, w[y] + k1w * dt / 2,p);
            k2w = h(u[y] + k1u * dt / 2, v[y] + k1v * dt / 2, w[y] + k1w * dt / 2,p);
            k3u = f(u[y] + k2u * dt / 2, v[y] + k2v * dt / 2, w[y] + k2w * dt / 2,p);
            k3v = g(u[y] + k2u * dt / 2, v[y] + k2v * dt / 2, w[y] + k2w * dt / 2,p);
            k3w = h(u[y] + k2u * dt / 2, v[y] + k2v * dt / 2, w[y] + k2w * dt / 2,p);
            k4u = f(u[y] + k3u * dt, v[y] + k3v * dt, w[y] + k3w * dt,p);
            k4v = g(u[y] + k3u * dt, v[y] + k3v * dt, w[y] + k3w * dt,p);
            k4w = h(u[y] + k3u * dt, v[y] + k3v * dt, w[y] + k3w * dt,p);
            unew[y] = u[y] + (k1u + 2 * k2u + 2 * k3u + k4u) / 6 * dt + Du * (u[(y+1)%ln]+u[(y-1+ln)%ln]-2*u[y])*dx2inv*dt;
            vnew[y] = v[y] + (k1v + 2 * k2v + 2 * k3v + k4v) / 6 * dt + Dv * (v[(y+1)%ln]+v[(y-1+ln)%ln]-2*v[y])*dx2inv*dt;
            wnew[y] = w[y] + (k1w + 2 * k2w + 2 * k3w + k4w) / 6 * dt;
          }
          for (y=0; y< ln; y++){
            uold[y] = u[y];
            u[y] = unew[y];
            v[y] = vnew[y];
            w[y] = wnew[y];          
          }
          u[0] = u[1];
          v[0] = v[1];
          w[0] = w[1];
          u[ln-1] = u[ln-2];
          v[ln-1] = v[ln-2];
          w[ln-1] = w[ln-2];        
          for (y=0; y< ln; y++){
            pv[y] = -((u[y]-uold[y])/dt)/((u[(y+1)%ln]-u[(y-1+ln)%ln])/2.0/dx);
          }
        }

        printf("%d %d\n",rep,n1);
      }
      fclose(outputfile);
    }
  //}
  return 0;
}

//file3.c
//Numerical simulation code with uniformly growing domains.

#include <stdio.h>
#include <stdlib.h>
#include <time.h>
#include "MT.h"// can be obtained from https://omitakahiro.github.io/random/code/MT.h

//Parallelization
//#ifdef _OPENMP
//#include <omp.h>
//#endif

#define _USE_MATH_DEFINES

#include <math.h>

#define stepsperout 620//time steps per output
#define outsteps 1000 //number of outputs

double f(double u, double v, double w, double c){
  return 0.57 *u - v + w - u*u*u + c*u*u;
}

double g(double u, double v, double w, double c){
  return 1.5 * u - 2.03 * v - v*v*v + c*v*v;
}

double h(double u, double v, double w, double c){
  return - u - 0.03 * w - w*w*w + c*w*w;
}

int main(void) {
  #pragma omp parallel
  {
    int rep;
    #pragma omp for
    for(rep=0;rep<1;rep=rep+1) {
      double dt = 0.001;//discrete time step
      int ln = 32*8+2;//lattice number
      int halfln = ln/2;
      int inithalfln = 5; // init L = 1.0
      int temphalfln = inithalfln;
      //double scaling[] = {0.8, 1.0, 1.2, 1.4, 1.6, 1.8, 2.0};
      double dx = 0.1;//spatial lattice size
      double dx2inv = 1.0/dx/dx;
      double dxinv = 1.0/dx;
      double s = 1.1867777442770433;//spatial scaling paramter throughout the study
      double Du = 0.03 * s;//diffusion coefficient for u
      double Dv = 1.0 * s;//diffusion coefficient for v
      int n1,m,y,n,x,j;
      double c = 0.05;//domain growth speed

      double *u, *v, *w, *uold, *pv;
      double *unew, *vnew, *wnew;
      u = (double*) malloc(sizeof(double) *  ln);
      v = (double*) malloc(sizeof(double) *  ln);
      w = (double*) malloc(sizeof(double) *  ln);
      unew = (double*) malloc(sizeof(double) *  ln);
      vnew = (double*) malloc(sizeof(double) *  ln);
      wnew = (double*) malloc(sizeof(double) *  ln);
      uold = (double*) malloc(sizeof(double) *  ln);
      pv = (double*) malloc(sizeof(double) *  ln);

      srand((unsigned)time(NULL)+rep*100);
      for (n1 = 0; n1 < ln; n1++){
        u[n1] = 0.0;
        v[n1] = 0.0;
        w[n1] = 0.0;
        pv[n1] = 0.0;
        if(halfln-temphalfln<=n1) {
          if(halfln+temphalfln>n1) {
            u[n1] = 50.0*(0.02 * rand()/RAND_MAX -0.01);
            v[n1] = 50.0*(0.02 * rand()/RAND_MAX -0.01);
            w[n1] = 50.0*(0.02 * rand()/RAND_MAX -0.01);
          }
        }
      }
      u[halfln-temphalfln-1] = u[halfln-temphalfln+1];
      v[halfln-temphalfln-1] = v[halfln-temphalfln+1];
      w[halfln-temphalfln-1] = w[halfln-temphalfln+1];
      u[halfln+temphalfln] = u[halfln+temphalfln-2];
      v[halfln+temphalfln] = v[halfln+temphalfln-2];
      w[halfln+temphalfln] = w[halfln+temphalfln-2];

      double t, k1u, k1v, k1w, k2u, k2v, k2w, k3u, k3v, k3w, k4u, k4v, k4w;
      n1 = 0;
      int currenthalfln;
      FILE *outputfile;
      char filename[50];
      sprintf(filename, "turing_feedback-%03d.txt",rep);
      outputfile = fopen(filename, "w");
      while(n1 < outsteps) {
        for (y=1; y<ln-1; y++){
          fprintf(outputfile, "%g\t", (u[y]));
        }
        for (y=1; y<ln-1; y++){
          fprintf(outputfile, "%g\t", (pv[y]));
        }
        for (y=1; y<ln-1; y++){
          fprintf(outputfile, "%g\t", (v[y]));
        }
        for (y=1; y<ln-1; y++){
          fprintf(outputfile, "%g\t", (w[y]));
        }
        fprintf(outputfile, "\n");
        
        n1++;
        for (m = 0; m < stepsperout; m++){
          t = dt * (m+stepsperout*n1);
          currenthalfln = inithalfln + t * c;
          if (currenthalfln > halfln-1) {
            currenthalfln = halfln-1;
          }
          if (currenthalfln > temphalfln) {
            temphalfln = currenthalfln;
            if(1>0) {//Implementation #1
              u[halfln-temphalfln-1] = u[halfln-temphalfln+1];
              v[halfln-temphalfln-1] = v[halfln-temphalfln+1];
              w[halfln-temphalfln-1] = w[halfln-temphalfln+1];
              u[halfln-temphalfln] = u[halfln-temphalfln+1];
              v[halfln-temphalfln] = v[halfln-temphalfln+1];
              w[halfln-temphalfln] = w[halfln-temphalfln+1];
              u[halfln+temphalfln] = u[halfln+temphalfln-2];
              v[halfln+temphalfln] = v[halfln+temphalfln-2];
              w[halfln+temphalfln] = w[halfln+temphalfln-2];
              u[halfln+temphalfln-1] = u[halfln+temphalfln-2];
              v[halfln+temphalfln-1] = v[halfln+temphalfln-2];
              w[halfln+temphalfln-1] = w[halfln+temphalfln-2];
            } else {//Implementation #2
              u[halfln-temphalfln-1] = 0.0;
              v[halfln-temphalfln-1] = 0.0;
              w[halfln-temphalfln-1] = 0.0;
              u[halfln-temphalfln] = 0.0;
              v[halfln-temphalfln] = 0.0;
              w[halfln-temphalfln] = 0.0;
              u[halfln+temphalfln] = 0.0;
              v[halfln+temphalfln] = 0.0;
              w[halfln+temphalfln] = 0.0;
              u[halfln+temphalfln-1] = 0.0;
              v[halfln+temphalfln-1] = 0.0;
              w[halfln+temphalfln-1] = 0.0;
            }
          }
          for (y=halfln-temphalfln; y<halfln+temphalfln; y++){
            k1u = f(u[y], v[y], w[y], 0.0);
            k1v = g(u[y], v[y], w[y], 0.0);
            k1w = h(u[y], v[y], w[y], 0.0);
            unew[y] = u[y] + k1u * dt + Du * dt * dx2inv * (u[(y+1)%ln]+u[(y-1+ln)%ln]-2*u[y]);
            vnew[y] = v[y] + k1v * dt + Dv * dt * dx2inv * (v[(y+1)%ln]+v[(y-1+ln)%ln]-2*v[y]);
            wnew[y] = w[y] + k1w * dt;
          }
          for (y=halfln-temphalfln; y<halfln+temphalfln; y++){
            uold[y] = u[y];
            u[y] = unew[y];
            v[y] = vnew[y];
            w[y] = wnew[y];
          }
          u[halfln-temphalfln-1] = u[halfln-temphalfln+1];
          v[halfln-temphalfln-1] = v[halfln-temphalfln+1];
          w[halfln-temphalfln-1] = w[halfln-temphalfln+1];
          u[halfln+temphalfln] = u[halfln+temphalfln-2];
          v[halfln+temphalfln] = v[halfln+temphalfln-2];
          w[halfln+temphalfln] = w[halfln+temphalfln-2];
          for (y=halfln-temphalfln; y<halfln+temphalfln; y++){
            pv[y] = -((u[y]-uold[y])/dt)/((u[(y+1)%ln]-u[(y-1+ln)%ln])/2.0/dx);
          }
        }

        printf("%d %d\n",rep,n1);
      }
      fclose(outputfile);
    }
  }
  return 0;
}

//file4.c
//Numerical simulation code with preferentially growing domains.

#include <stdio.h>
#include <stdlib.h>
#include <time.h>
#include "MT.h"// can be obtained from https://omitakahiro.github.io/random/code/MT.h

//Parallelization
//#ifdef _OPENMP
//#include <omp.h>
//#endif

#define _USE_MATH_DEFINES
#include <math.h>

#define outsteps 1000 //number of outputs

double f(double u, double v, double w){
  return 0.57 * u - v + w - u*u*u;
}

double g(double u, double v, double w){
  return 1.5 * u - 2.03 * v - v*v*v;
}

double h(double u, double v, double w){
  return - u - 0.03 * w - w*w*w;
}

int main(void) {
  #pragma omp parallel
  {
    int rep;
    #pragma omp for
    for(rep=0;rep<1;rep=rep+1) {
      int stepsperout = 620*200;//time steps per output
      double dt = 0.001/200;//timestep
      int ln = 32*8+2;//lattice number
      double l0 = 1.0;//init L = 1.0
      double dx = l0/(ln-2);//spatial lattice size
      double dx2inv = 1.0/dx/dx;
      double dxinv = 1.0/dx;
      double s = 1.1867777442770433;//spatial scaling paramter throughout the study
      double Du = 0.03 * s;//diffusion coefficient for u
      double Dv = 1.0 * s;//diffusion coefficient for v
      int n1,m,y,n,x,j;
      double clist[] = {0.02,0.05,0.1,0.2,0.3,0.4,0.5,1.0,2.0,5.0,10.0,20.0};
      double c = 0.05;//domain growth speed
      
      double t;
      double *u, *v, *w, *uold, *pv;
      double *unew, *vnew, *wnew;
      u = (double*) malloc(sizeof(double) *  ln);
      v = (double*) malloc(sizeof(double) *  ln);
      w = (double*) malloc(sizeof(double) *  ln);
      unew = (double*) malloc(sizeof(double) *  ln);
      vnew = (double*) malloc(sizeof(double) *  ln);
      wnew = (double*) malloc(sizeof(double) *  ln);
      uold = (double*) malloc(sizeof(double) *  ln);
      pv = (double*) malloc(sizeof(double) *  ln);

      // フーリエ変換前の数列値を設定
      srand((unsigned)time(NULL)+rep*100);
      for (n1 = 0; n1 < ln; n1++){
        u[n1] = 50.0 * (0.02 * rand()/RAND_MAX -0.01);
        v[n1] = 50.0 * (0.02 * rand()/RAND_MAX -0.01);
        w[n1] = 50.0 * (0.02 * rand()/RAND_MAX -0.01);
        pv[n1] = 0.0;
      }

      double k1u, k1v, k1w, k2u, k2v, k2w, k3u, k3v, k3w, k4u, k4v, k4w, u_reaction, v_reaction, w_reaction, u_diffusion, v_diffusion, w_diffusion, l2normdif, l2normold, l2norm;
      double l2normcutoff = 1.0e-10;
      l2normold = 1000.0;
      l2normdif = 1000.0;
      n1 = 0;
      FILE *outputfile;
      char filename[50];
      sprintf(filename, "turing_feedback-%03d.txt",rep);
      outputfile = fopen(filename, "w");
      while(n1 < outsteps) {
        for (y=0; y<ln; y++){
          fprintf(outputfile, "%g\t", (u[y]));
        }
        for (y=0; y<ln; y++){
          fprintf(outputfile, "%g\t", (pv[y]));
        }
        for (y=0; y<ln; y++){
          fprintf(outputfile, "%g\t", (v[y]));
        }
        for (y=0; y<ln; y++){
          fprintf(outputfile, "%g\t", (w[y]));
        }
        fprintf(outputfile, "\n");

        n1++;
        for (m = 0; m < stepsperout; m++){
          t = dt * (m+stepsperout*n1);
          dx = (l0 + 2.0 * c * t * 0.1)/(ln-2);
          if(dx > 0.1) {
            dx = 0.1;
          }
          dx2inv = 1.0/dx/dx;
          dxinv = 1.0/dx;
          for (y=0; y< ln; y++){
            k1u = f(u[y], v[y], w[y]);
            k1v = g(u[y], v[y], w[y]);
            k1w = h(u[y], v[y], w[y]);
            k2u = f(u[y] + k1u * dt / 2, v[y] + k1v * dt / 2, w[y] + k1w * dt / 2);
            k2v = g(u[y] + k1u * dt / 2, v[y] + k1v * dt / 2, w[y] + k1w * dt / 2);
            k2w = h(u[y] + k1u * dt / 2, v[y] + k1v * dt / 2, w[y] + k1w * dt / 2);
            k3u = f(u[y] + k2u * dt / 2, v[y] + k2v * dt / 2, w[y] + k2w * dt / 2);
            k3v = g(u[y] + k2u * dt / 2, v[y] + k2v * dt / 2, w[y] + k2w * dt / 2);
            k3w = h(u[y] + k2u * dt / 2, v[y] + k2v * dt / 2, w[y] + k2w * dt / 2);
            k4u = f(u[y] + k3u * dt, v[y] + k3v * dt, w[y] + k3w * dt);
            k4v = g(u[y] + k3u * dt, v[y] + k3v * dt, w[y] + k3w * dt);
            k4w = h(u[y] + k3u * dt, v[y] + k3v * dt, w[y] + k3w * dt);
            unew[y] = u[y] + (k1u + 2 * k2u + 2 * k3u + k4u) / 6 * dt + Du * (u[(y+1)%ln]+u[(y-1+ln)%ln]-2*u[y])*dx2inv*dt;
            vnew[y] = v[y] + (k1v + 2 * k2v + 2 * k3v + k4v) / 6 * dt + Dv * (v[(y+1)%ln]+v[(y-1+ln)%ln]-2*v[y])*dx2inv*dt;
            wnew[y] = w[y] + (k1w + 2 * k2w + 2 * k3w + k4w) / 6 * dt;
          }
          for (y=0; y< ln; y++){
            uold[y] = u[y];
            u[y] = unew[y];
            v[y] = vnew[y];
            w[y] = wnew[y];          
          }
          
          u[0] = u[1];
          v[0] = v[1];
          w[0] = w[1];
          u[ln-1] = u[ln-2];
          v[ln-1] = v[ln-2];
          w[ln-1] = w[ln-2];
        
          for (y=0; y< ln; y++){
            pv[y] = -((u[y]-uold[y])/dt)/((u[(y+1)%ln]-u[(y-1+ln)%ln])/2.0/dx);
          }
        }
        printf("%d %d\n",rep,n1);
      }
      fclose(outputfile);
    }
  }
  return 0;
}

//file5.c
//Numerical simiulation code using predetermined initial conditions with Neumann boundary conditions.
//Note that in the codes, we used the different scales than in the manuscript.
//L = 3.2 and T = 6.2 in the codes corresponds to L = 1.8 and T = 8.6, respectively.

#include <stdio.h>
#include <stdlib.h>
#include <time.h>
#include "MT.h" // can be obtained from https://omitakahiro.github.io/random/code/MT.h

//Parallelization
#ifdef _OPENMP
#include <omp.h>
#endif

#define _USE_MATH_DEFINES

#include <math.h>

#define outsteps 1000 //number of outputs

double f(double u, double v, double w, double p){
  return (0.5-p) *u - v + w - u*u*u;
}

double g(double u, double v, double w, double p){
  return 1.5 * u - (2.1+p) * v - v*v*v;
}

double h(double u, double v, double w, double p){
  return - u - (0.1+p) * w - w*w*w;
}

int main(void) {
  #pragma omp parallel
  {
    int rep;
    #pragma omp for
    for(rep=0;rep<11*16;rep=rep+1) {
      double dt = 0.001;
      long long int stepsperout = 1000000;//time steps per output
      int ln = 32*32;
      double dx = 0.1;//spatial lattice size
      double dx2inv = 1.0/dx/dx;
      double s = 1.1867777442770433;//spatial scaling paramter throughout the study
      double Du = 0.03 * s;//diffusion coefficient for u
      double Dv = 1.0 * s;//diffusion coefficient for v
      double r = 50.0;//white noise amplitude
      double plist[] = {-0.07,-0.06,-0.05,-0.04,-0.03,-0.02,-0.01,0.0,0.01,0.02,0.0237};
      double p = plist[rep/16];//control parameter \mu
      int thy = 32*(rep%16);//lattice number of L1 (flipped region)
      double initlist[11][5] = {{0.354217, 0.0785467, 0.0120015, -0.0414038, -0.345755}, \
      {0.334826, 0.0741455, 0.0113118, -0.0392455, -0.32683},\
       {0.314241, 0.0694924, 0.0105858, -0.036934, -0.30674},\
        {0.292212, 0.0645327, 0.00981539, -0.0344386, -0.285239}, {0.268381, 0.0591894, \
        0.00898904, -0.0317158, -0.261979}, {0.242219, 0.0533469, \
        0.0080895, -0.0287012, -0.236443}, {0.212867, 0.0468188, \
        0.00708884, -0.0252907, -0.207793}, {0.178761, 0.039264, \
        0.00593601, -0.021295, -0.174501}, {0.136379, 0.0299145, \
        0.00451571, -0.0162892, -0.13313}, {0.0724136, 0.0158623, \
        0.00239087, -0.0086718, -0.0706893}, {0.0173891, 0.00380721, \
        0.000573529, -0.00208443, -0.0169751}};
      int n1,m,y;

      FILE *outputfile;
      char filename[50];
      sprintf(filename, "turing_feedback-%03d.txt", rep);
      outputfile = fopen(filename, "w"); 
      if (outputfile == NULL) {          
        printf("cannot open\n");        
        exit(1);                        
      }

      double *u, *v, *w, *uold, *pv;
      double *unew, *vnew, *wnew;
      u = (double*) malloc(sizeof(double) *  ln);
      uold = (double*) malloc(sizeof(double) *  ln);
      pv = (double*) malloc(sizeof(double) *  ln);
      v = (double*) malloc(sizeof(double) *  ln);
      w = (double*) malloc(sizeof(double) *  ln);
      unew = (double*) malloc(sizeof(double) *  ln);
      vnew = (double*) malloc(sizeof(double) *  ln);
      wnew = (double*) malloc(sizeof(double) *  ln);

      srand((unsigned)time(NULL)+rep*100);
      for (n1 = 0; n1 < ln; n1++){
        y = n1;
        double k,u1,v1,v2,w1,w2;
        k = 2.0*M_PI/(32*dx);
        u1 = initlist[rep/16][0];
        v1 = initlist[rep/16][1];
        v2 = initlist[rep/16][2];
        w1 = initlist[rep/16][3];
        w2 = initlist[rep/16][4];
        u[n1] = u1 * sin(k*y*dx);
        v[n1] = v1 * sin(k*y*dx) + v2 * cos(k*y*dx);
        w[n1] = w1 * sin(k*y*dx) + w2 * cos(k*y*dx);
      }

      double k1u, k1v, k1w, k2u, k2v, k2w, k3u, k3v, k3w, k4u, k4v, k4w;
      
      //convergence of pure patterns
      for (m = 0; m < stepsperout; m++){
        for (y=0; y< ln; y++){
          k1u = f(u[y], v[y], w[y],p);
          k1v = g(u[y], v[y], w[y],p);
          k1w = h(u[y], v[y], w[y],p);
          k2u = f(u[y] + k1u * dt / 2, v[y] + k1v * dt / 2, w[y] + k1w * dt / 2,p);
          k2v = g(u[y] + k1u * dt / 2, v[y] + k1v * dt / 2, w[y] + k1w * dt / 2,p);
          k2w = h(u[y] + k1u * dt / 2, v[y] + k1v * dt / 2, w[y] + k1w * dt / 2,p);
          k3u = f(u[y] + k2u * dt / 2, v[y] + k2v * dt / 2, w[y] + k2w * dt / 2,p);
          k3v = g(u[y] + k2u * dt / 2, v[y] + k2v * dt / 2, w[y] + k2w * dt / 2,p);
          k3w = h(u[y] + k2u * dt / 2, v[y] + k2v * dt / 2, w[y] + k2w * dt / 2,p);
          k4u = f(u[y] + k3u * dt, v[y] + k3v * dt, w[y] + k3w * dt,p);
          k4v = g(u[y] + k3u * dt, v[y] + k3v * dt, w[y] + k3w * dt,p);
          k4w = h(u[y] + k3u * dt, v[y] + k3v * dt, w[y] + k3w * dt,p);
          unew[y] = u[y] + (k1u + 2 * k2u + 2 * k3u + k4u) / 6 * dt + Du * (u[(y+1)%ln]+u[(y-1+ln)%ln]-2*u[y])*dx2inv*dt;
          vnew[y] = v[y] + (k1v + 2 * k2v + 2 * k3v + k4v) / 6 * dt + Dv * (v[(y+1)%ln]+v[(y-1+ln)%ln]-2*v[y])*dx2inv*dt;
          wnew[y] = w[y] + (k1w + 2 * k2w + 2 * k3w + k4w) / 6 * dt;
        }
        for (y=0; y< ln; y++){
          uold[y] = u[y];
          u[y] = unew[y];
          v[y] = vnew[y];
          w[y] = wnew[y];
        }
        for (y=0; y< ln; y++){
          pv[y] = -((u[y]-uold[y])/dt)/((u[(y+1)%ln]-u[(y-1+ln)%ln])/2.0/dx);
        }
      }
      //flip L1 in x directoin
      for (y=0; y< thy; y++){
        u[y] = unew[thy-y];
        v[y] = vnew[thy-y];
        w[y] = wnew[thy-y];
      }
      for (y=0; y<ln; y++){
        fprintf(outputfile, "%g\t", (u[y]));
      }
      for (y=0; y<ln; y++){
        fprintf(outputfile, "%g\t", (pv[y]));
      }
      fprintf(outputfile, "\n");
      //use the partially flipped pattern as an initial condition
      n1 = 0;
      while(n1 < outsteps) {
        n1++;
        for (m = 0; m < stepsperout; m++){
          for (y=0; y< ln; y++){
            k1u = f(u[y], v[y], w[y],p);
            k1v = g(u[y], v[y], w[y],p);
            k1w = h(u[y], v[y], w[y],p);
            k2u = f(u[y] + k1u * dt / 2, v[y] + k1v * dt / 2, w[y] + k1w * dt / 2,p);
            k2v = g(u[y] + k1u * dt / 2, v[y] + k1v * dt / 2, w[y] + k1w * dt / 2,p);
            k2w = h(u[y] + k1u * dt / 2, v[y] + k1v * dt / 2, w[y] + k1w * dt / 2,p);
            k3u = f(u[y] + k2u * dt / 2, v[y] + k2v * dt / 2, w[y] + k2w * dt / 2,p);
            k3v = g(u[y] + k2u * dt / 2, v[y] + k2v * dt / 2, w[y] + k2w * dt / 2,p);
            k3w = h(u[y] + k2u * dt / 2, v[y] + k2v * dt / 2, w[y] + k2w * dt / 2,p);
            k4u = f(u[y] + k3u * dt, v[y] + k3v * dt, w[y] + k3w * dt,p);
            k4v = g(u[y] + k3u * dt, v[y] + k3v * dt, w[y] + k3w * dt,p);
            k4w = h(u[y] + k3u * dt, v[y] + k3v * dt, w[y] + k3w * dt,p);
            unew[y] = u[y] + (k1u + 2 * k2u + 2 * k3u + k4u) / 6 * dt + Du * (u[(y+1)%ln]+u[(y-1+ln)%ln]-2*u[y])*dx2inv*dt;
            vnew[y] = v[y] + (k1v + 2 * k2v + 2 * k3v + k4v) / 6 * dt + Dv * (v[(y+1)%ln]+v[(y-1+ln)%ln]-2*v[y])*dx2inv*dt;
            wnew[y] = w[y] + (k1w + 2 * k2w + 2 * k3w + k4w) / 6 * dt;
          }
          for (y=0; y< ln; y++){
            uold[y] = u[y];
            u[y] = unew[y];
            v[y] = vnew[y];
            w[y] = wnew[y];
          }
          for (y=0; y< ln; y++){
            pv[y] = -((u[y]-uold[y])/dt)/((u[(y+1)%ln]-u[(y-1+ln)%ln])/2.0/dx);
          }
        }
        printf("%d %d %g %g\n",rep,n1);
        for (y=0; y<ln; y++){
          fprintf(outputfile, "%g\t", (u[y])); // ファイルに書く
        }
        for (y=0; y<ln; y++){
          fprintf(outputfile, "%g\t", (pv[y])); // ファイルに書く
        }
        fprintf(outputfile, "\n");
      }
      fclose(outputfile);
    }
  }
  return 0;
}

//file6.c
//Numerical simulation code using Neumann boundary conditions where two domains inwardly expands and merge, corresponding to Fig. 7a,b
//Note that in the codes, we used the different scales than in the manuscript.
//L = 3.2 and T = 6.2 in the codes corresponds to L = 1.8 and T = 8.6, respectively.

#include <stdio.h>
#include <stdlib.h>
#include <time.h>
#include "MT.h" // can be obtained from https://omitakahiro.github.io/random/code/MT.h

//Parallelization
//#ifdef _OPENMP
//#include <omp.h>
//#endif

#define _USE_MATH_DEFINES

#include <math.h>

#define stepsperout 620 //time steps per output
#define outsteps 1000 //number of outputs

double f(double u, double v, double w){
  return 0.57 *u - v + w - u*u*u;
}

double g(double u, double v, double w){
  return 1.5 * u - 2.03 * v - v*v*v;
}

double h(double u, double v, double w){
  return - u - 0.03 * w - w*w*w; 
}

int main(void) {
  //#pragma omp parallel
  //{
    int rep;
    //#pragma omp for
    for(rep=0;rep<1;rep=rep+1) {
      double dt = 0.001;//discrete time step
      int lninit = 8;// initial lattice number
      int lnfin = 32*8;// final lattice number
      int ln;
      double dx = 0.1;//spatial lattice size
      double dx2inv = 1.0/dx/dx;
      double s = 1.1867777442770433;//spatial scaling paramter throughout the study
      double Du = 0.03 * s;//diffusion coefficient for u
      double Dv = 1.0 * s;//diffusion coefficient for v
      double p = -0.07;//control parameter \mu
      double r = 50.0;//white noise amplitude
      int n1,m,y;
      double t;

      double *u, *v, *w, *uold, *pv;
      double *unew, *vnew, *wnew;
      u = (double*) malloc(sizeof(double) *  lnfin);
      v = (double*) malloc(sizeof(double) *  lnfin);
      w = (double*) malloc(sizeof(double) *  lnfin);
      unew = (double*) malloc(sizeof(double) *  lnfin);
      vnew = (double*) malloc(sizeof(double) *  lnfin);
      wnew = (double*) malloc(sizeof(double) *  lnfin);
      uold = (double*) malloc(sizeof(double) *  lnfin);
      pv = (double*) malloc(sizeof(double) *  lnfin);

      // フーリエ変換前の数列値を設定
      srand((unsigned)time(NULL)+rep*100);
      for (n1 = 0; n1 < lnfin; n1++){
        if(n1<lninit/2) {
          u[n1] = (0.02 * rand()/RAND_MAX -0.01)*50;
          v[n1] = (0.02 * rand()/RAND_MAX -0.01)*50;
          w[n1] = (0.02 * rand()/RAND_MAX -0.01)*50;
          unew[n1] = 0.0;
          vnew[n1] = 0.0;
          wnew[n1] = 0.0;
        } else {
          if(n1>=lnfin-lninit/2){
            u[n1] = (0.02 * rand()/RAND_MAX -0.01)*50;
            v[n1] = (0.02 * rand()/RAND_MAX -0.01)*50;
            w[n1] = (0.02 * rand()/RAND_MAX -0.01)*50;
            unew[n1] = 0.0;
            vnew[n1] = 0.0;
            wnew[n1] = 0.0;
          } else {
            u[n1] = 0.0;
            v[n1] = 0.0;
            w[n1] = 0.0;
            unew[n1] = 0.0;
            vnew[n1] = 0.0;
            wnew[n1] = 0.0;
          }
        }
        pv[n1] = 0.0;
      }

      FILE *outputfile;
      char filename[50];
      sprintf(filename, "turing_feedback-%03d.txt",rep);
      outputfile = fopen(filename, "w");

      for (y=0; y<lnfin; y++){
        fprintf(outputfile, "%g\t", (u[y]));
      }
      for (y=0; y<lnfin; y++){
        fprintf(outputfile, "%g\t", (pv[y]));
      }
      for (y=0; y<lnfin; y++){
        fprintf(outputfile, "%g\t", (v[y]));
      }
      for (y=0; y<lnfin; y++){
        fprintf(outputfile, "%g\t", (w[y]));
      }
      for (y=0; y<lnfin; y++){
        fprintf(outputfile, "%g\t", (uold[y]));
      }
      fprintf(outputfile, "\n");

      double ku,kv,kw,k1u, k1v, k1w, k2u, k2v, k2w, k3u, k3v, k3w, k4u, k4v, k4w;
      
      n1 = 0;
      while(n1 < outsteps) {
        n1++;
        for (m = 0; m < stepsperout; m++){
          t = dt * (n1 * stepsperout + m);
          ln = (int)(lninit + (lnfin-lninit)*t/30.28);
          ln = (ln/2)*2;
          if(ln>lnfin) {
            ln = lnfin;
            for (y= 1; y< ln-1; y++){
              k1u = f(u[y], v[y], w[y]);
              k1v = g(u[y], v[y], w[y]);
              k1w = h(u[y], v[y], w[y]);
              k2u = f(u[y] + k1u * dt / 2, v[y] + k1v * dt / 2, w[y] + k1w * dt / 2);
              k2v = g(u[y] + k1u * dt / 2, v[y] + k1v * dt / 2, w[y] + k1w * dt / 2);
              k2w = h(u[y] + k1u * dt / 2, v[y] + k1v * dt / 2, w[y] + k1w * dt / 2);
              k3u = f(u[y] + k2u * dt / 2, v[y] + k2v * dt / 2, w[y] + k2w * dt / 2);
              k3v = g(u[y] + k2u * dt / 2, v[y] + k2v * dt / 2, w[y] + k2w * dt / 2);
              k3w = h(u[y] + k2u * dt / 2, v[y] + k2v * dt / 2, w[y] + k2w * dt / 2);
              k4u = f(u[y] + k3u * dt, v[y] + k3v * dt, w[y] + k3w * dt);
              k4v = g(u[y] + k3u * dt, v[y] + k3v * dt, w[y] + k3w * dt);
              k4w = h(u[y] + k3u * dt, v[y] + k3v * dt, w[y] + k3w * dt);
              ku = (k1u + 2 * k2u + 2 * k3u + k4u) / 6;
              kv = (k1v + 2 * k2v + 2 * k3v + k4v) / 6;
              kw = (k1w + 2 * k2w + 2 * k3w + k4w) / 6;
              unew[y] = u[y] + ku * dt + Du * dt * dx2inv * (u[y+1]+u[y-1]-2*u[y]);
              vnew[y] = v[y] + kv * dt + Dv * dt * dx2inv * (v[y+1]+v[y-1]-2*v[y]);
              wnew[y] = w[y] + kw * dt;
            }
            y = 0;
            k1u = f(u[y], v[y], w[y]);
            k1v = g(u[y], v[y], w[y]);
            k1w = h(u[y], v[y], w[y]);
            k2u = f(u[y] + k1u * dt / 2, v[y] + k1v * dt / 2, w[y] + k1w * dt / 2);
            k2v = g(u[y] + k1u * dt / 2, v[y] + k1v * dt / 2, w[y] + k1w * dt / 2);
            k2w = h(u[y] + k1u * dt / 2, v[y] + k1v * dt / 2, w[y] + k1w * dt / 2);
            k3u = f(u[y] + k2u * dt / 2, v[y] + k2v * dt / 2, w[y] + k2w * dt / 2);
            k3v = g(u[y] + k2u * dt / 2, v[y] + k2v * dt / 2, w[y] + k2w * dt / 2);
            k3w = h(u[y] + k2u * dt / 2, v[y] + k2v * dt / 2, w[y] + k2w * dt / 2);
            k4u = f(u[y] + k3u * dt, v[y] + k3v * dt, w[y] + k3w * dt);
            k4v = g(u[y] + k3u * dt, v[y] + k3v * dt, w[y] + k3w * dt);
            k4w = h(u[y] + k3u * dt, v[y] + k3v * dt, w[y] + k3w * dt);
            ku = (k1u + 2 * k2u + 2 * k3u + k4u) / 6;
            kv = (k1v + 2 * k2v + 2 * k3v + k4v) / 6;
            kw = (k1w + 2 * k2w + 2 * k3w + k4w) / 6;
            unew[y] = u[y] + ku * dt + Du * dt * dx2inv * (u[y+1]+u[y+1]-2*u[y]);
            vnew[y] = v[y] + kv * dt + Dv * dt * dx2inv * (v[y+1]+v[y+1]-2*v[y]);
            wnew[y] = w[y] + kw * dt;
            y = ln-1;
            k1u = f(u[y], v[y], w[y]);
            k1v = g(u[y], v[y], w[y]);
            k1w = h(u[y], v[y], w[y]);k2u = f(u[y] + k1u * dt / 2, v[y] + k1v * dt / 2, w[y] + k1w * dt / 2);
            k2v = g(u[y] + k1u * dt / 2, v[y] + k1v * dt / 2, w[y] + k1w * dt / 2);
            k2w = h(u[y] + k1u * dt / 2, v[y] + k1v * dt / 2, w[y] + k1w * dt / 2);
            k3u = f(u[y] + k2u * dt / 2, v[y] + k2v * dt / 2, w[y] + k2w * dt / 2);
            k3v = g(u[y] + k2u * dt / 2, v[y] + k2v * dt / 2, w[y] + k2w * dt / 2);
            k3w = h(u[y] + k2u * dt / 2, v[y] + k2v * dt / 2, w[y] + k2w * dt / 2);
            k4u = f(u[y] + k3u * dt, v[y] + k3v * dt, w[y] + k3w * dt);
            k4v = g(u[y] + k3u * dt, v[y] + k3v * dt, w[y] + k3w * dt);
            k4w = h(u[y] + k3u * dt, v[y] + k3v * dt, w[y] + k3w * dt);
            ku = (k1u + 2 * k2u + 2 * k3u + k4u) / 6;
            kv = (k1v + 2 * k2v + 2 * k3v + k4v) / 6;
            kw = (k1w + 2 * k2w + 2 * k3w + k4w) / 6;
            unew[y] = u[y] + ku * dt + Du * dt * dx2inv * (u[y-1]+u[y-1]-2*u[y]);
            vnew[y] = v[y] + kv * dt + Dv * dt * dx2inv * (v[y-1]+v[y-1]-2*v[y]);
            wnew[y] = w[y] + kw * dt;
            for (y=0; y< lnfin; y++){
              uold[y] = u[y];
              u[y] = unew[y];
              v[y] = vnew[y];
              w[y] = wnew[y];
            }
            y = 0;
            pv[y] = 0.0;
            y = ln-1;
            pv[y] = 0.0;
            for (y=1; y< ln-1; y++){
              pv[y] = -((u[y]-uold[y])/dt)/((u[(y+1)%lnfin]-u[(y-1+ln)%lnfin])/2.0/dx);
            }
          } else {
            for (y= 1; y< ln/2-1; y++){
              k1u = f(u[y], v[y], w[y]);
              k1v = g(u[y], v[y], w[y]);
              k1w = h(u[y], v[y], w[y]);
              k2u = f(u[y] + k1u * dt / 2, v[y] + k1v * dt / 2, w[y] + k1w * dt / 2);
              k2v = g(u[y] + k1u * dt / 2, v[y] + k1v * dt / 2, w[y] + k1w * dt / 2);
              k2w = h(u[y] + k1u * dt / 2, v[y] + k1v * dt / 2, w[y] + k1w * dt / 2);
              k3u = f(u[y] + k2u * dt / 2, v[y] + k2v * dt / 2, w[y] + k2w * dt / 2);
              k3v = g(u[y] + k2u * dt / 2, v[y] + k2v * dt / 2, w[y] + k2w * dt / 2);
              k3w = h(u[y] + k2u * dt / 2, v[y] + k2v * dt / 2, w[y] + k2w * dt / 2);
              k4u = f(u[y] + k3u * dt, v[y] + k3v * dt, w[y] + k3w * dt);
              k4v = g(u[y] + k3u * dt, v[y] + k3v * dt, w[y] + k3w * dt);
              k4w = h(u[y] + k3u * dt, v[y] + k3v * dt, w[y] + k3w * dt);
              ku = (k1u + 2 * k2u + 2 * k3u + k4u) / 6;
              kv = (k1v + 2 * k2v + 2 * k3v + k4v) / 6;
              kw = (k1w + 2 * k2w + 2 * k3w + k4w) / 6;
              unew[y] = u[y] + ku * dt + Du * dt * dx2inv * (u[y+1]+u[y-1]-2*u[y]);
              vnew[y] = v[y] + kv * dt + Dv * dt * dx2inv * (v[y+1]+v[y-1]-2*v[y]);
              wnew[y] = w[y] + kw * dt;
            }
            y = 0;
            k1u = f(u[y], v[y], w[y]);
            k1v = g(u[y], v[y], w[y]);
            k1w = h(u[y], v[y], w[y]);
            k2u = f(u[y] + k1u * dt / 2, v[y] + k1v * dt / 2, w[y] + k1w * dt / 2);
            k2v = g(u[y] + k1u * dt / 2, v[y] + k1v * dt / 2, w[y] + k1w * dt / 2);
            k2w = h(u[y] + k1u * dt / 2, v[y] + k1v * dt / 2, w[y] + k1w * dt / 2);
            k3u = f(u[y] + k2u * dt / 2, v[y] + k2v * dt / 2, w[y] + k2w * dt / 2);
            k3v = g(u[y] + k2u * dt / 2, v[y] + k2v * dt / 2, w[y] + k2w * dt / 2);
            k3w = h(u[y] + k2u * dt / 2, v[y] + k2v * dt / 2, w[y] + k2w * dt / 2);
            k4u = f(u[y] + k3u * dt, v[y] + k3v * dt, w[y] + k3w * dt);
            k4v = g(u[y] + k3u * dt, v[y] + k3v * dt, w[y] + k3w * dt);
            k4w = h(u[y] + k3u * dt, v[y] + k3v * dt, w[y] + k3w * dt);
            ku = (k1u + 2 * k2u + 2 * k3u + k4u) / 6;
            kv = (k1v + 2 * k2v + 2 * k3v + k4v) / 6;
            kw = (k1w + 2 * k2w + 2 * k3w + k4w) / 6;
            unew[y] = u[y] + ku * dt + Du * dt * dx2inv * (u[y+1]+u[y+1]-2*u[y]);
            vnew[y] = v[y] + kv * dt + Dv * dt * dx2inv * (v[y+1]+v[y+1]-2*v[y]);
            wnew[y] = w[y] + kw * dt;
            y = ln-1;
            k1u = f(u[y], v[y], w[y]);
            k1v = g(u[y], v[y], w[y]);
            k1w = h(u[y], v[y], w[y]);
            k2u = f(u[y] + k1u * dt / 2, v[y] + k1v * dt / 2, w[y] + k1w * dt / 2);
            k2v = g(u[y] + k1u * dt / 2, v[y] + k1v * dt / 2, w[y] + k1w * dt / 2);
            k2w = h(u[y] + k1u * dt / 2, v[y] + k1v * dt / 2, w[y] + k1w * dt / 2);
            k3u = f(u[y] + k2u * dt / 2, v[y] + k2v * dt / 2, w[y] + k2w * dt / 2);
            k3v = g(u[y] + k2u * dt / 2, v[y] + k2v * dt / 2, w[y] + k2w * dt / 2);
            k3w = h(u[y] + k2u * dt / 2, v[y] + k2v * dt / 2, w[y] + k2w * dt / 2);
            k4u = f(u[y] + k3u * dt, v[y] + k3v * dt, w[y] + k3w * dt);
            k4v = g(u[y] + k3u * dt, v[y] + k3v * dt, w[y] + k3w * dt);
            k4w = h(u[y] + k3u * dt, v[y] + k3v * dt, w[y] + k3w * dt);
            ku = (k1u + 2 * k2u + 2 * k3u + k4u) / 6;
            kv = (k1v + 2 * k2v + 2 * k3v + k4v) / 6;
            kw = (k1w + 2 * k2w + 2 * k3w + k4w) / 6;
            unew[y] = u[y] + ku * dt + Du * dt * dx2inv * (u[y-1]+u[y-1]-2*u[y]);
            vnew[y] = v[y] + kv * dt + Dv * dt * dx2inv * (v[y-1]+v[y-1]-2*v[y]);
            wnew[y] = w[y] + kw * dt;
            for (y= lnfin-ln/2+1; y< lnfin-1; y++){
              k1u = f(u[y], v[y], w[y]);
              k1v = g(u[y], v[y], w[y]);
              k1w = h(u[y], v[y], w[y]);
              k2u = f(u[y] + k1u * dt / 2, v[y] + k1v * dt / 2, w[y] + k1w * dt / 2);
              k2v = g(u[y] + k1u * dt / 2, v[y] + k1v * dt / 2, w[y] + k1w * dt / 2);
              k2w = h(u[y] + k1u * dt / 2, v[y] + k1v * dt / 2, w[y] + k1w * dt / 2);
              k3u = f(u[y] + k2u * dt / 2, v[y] + k2v * dt / 2, w[y] + k2w * dt / 2);
              k3v = g(u[y] + k2u * dt / 2, v[y] + k2v * dt / 2, w[y] + k2w * dt / 2);
              k3w = h(u[y] + k2u * dt / 2, v[y] + k2v * dt / 2, w[y] + k2w * dt / 2);
              k4u = f(u[y] + k3u * dt, v[y] + k3v * dt, w[y] + k3w * dt);
              k4v = g(u[y] + k3u * dt, v[y] + k3v * dt, w[y] + k3w * dt);
              k4w = h(u[y] + k3u * dt, v[y] + k3v * dt, w[y] + k3w * dt);
              ku = (k1u + 2 * k2u + 2 * k3u + k4u) / 6;
              kv = (k1v + 2 * k2v + 2 * k3v + k4v) / 6;
              kw = (k1w + 2 * k2w + 2 * k3w + k4w) / 6;
              unew[y] = u[y] + ku * dt + Du * dt * dx2inv * (u[y+1]+u[y-1]-2*u[y]);
              vnew[y] = v[y] + kv * dt + Dv * dt * dx2inv * (v[y+1]+v[y-1]-2*v[y]);
              wnew[y] = w[y] + kw * dt;
            }
            y = lnfin-ln/2;
            k1u = f(u[y], v[y], w[y]);
            k1v = g(u[y], v[y], w[y]);
            k1w = h(u[y], v[y], w[y]);
            k2u = f(u[y] + k1u * dt / 2, v[y] + k1v * dt / 2, w[y] + k1w * dt / 2);
            k2v = g(u[y] + k1u * dt / 2, v[y] + k1v * dt / 2, w[y] + k1w * dt / 2);
            k2w = h(u[y] + k1u * dt / 2, v[y] + k1v * dt / 2, w[y] + k1w * dt / 2);
            k3u = f(u[y] + k2u * dt / 2, v[y] + k2v * dt / 2, w[y] + k2w * dt / 2);
            k3v = g(u[y] + k2u * dt / 2, v[y] + k2v * dt / 2, w[y] + k2w * dt / 2);
            k3w = h(u[y] + k2u * dt / 2, v[y] + k2v * dt / 2, w[y] + k2w * dt / 2);
            k4u = f(u[y] + k3u * dt, v[y] + k3v * dt, w[y] + k3w * dt);
            k4v = g(u[y] + k3u * dt, v[y] + k3v * dt, w[y] + k3w * dt);
            k4w = h(u[y] + k3u * dt, v[y] + k3v * dt, w[y] + k3w * dt);
            ku = (k1u + 2 * k2u + 2 * k3u + k4u) / 6;
            kv = (k1v + 2 * k2v + 2 * k3v + k4v) / 6;
            kw = (k1w + 2 * k2w + 2 * k3w + k4w) / 6;
            unew[y] = u[y] + ku * dt + Du * dt * dx2inv * (u[y+1]+u[y+1]-2*u[y]);
            vnew[y] = v[y] + kv * dt + Dv * dt * dx2inv * (v[y+1]+v[y+1]-2*v[y]);
            wnew[y] = w[y] + kw * dt;
            y = lnfin-1;
            k1u = f(u[y], v[y], w[y]);
            k1v = g(u[y], v[y], w[y]);
            k1w = h(u[y], v[y], w[y]);k2u = f(u[y] + k1u * dt / 2, v[y] + k1v * dt / 2, w[y] + k1w * dt / 2);
            k2v = g(u[y] + k1u * dt / 2, v[y] + k1v * dt / 2, w[y] + k1w * dt / 2);
            k2w = h(u[y] + k1u * dt / 2, v[y] + k1v * dt / 2, w[y] + k1w * dt / 2);
            k3u = f(u[y] + k2u * dt / 2, v[y] + k2v * dt / 2, w[y] + k2w * dt / 2);
            k3v = g(u[y] + k2u * dt / 2, v[y] + k2v * dt / 2, w[y] + k2w * dt / 2);
            k3w = h(u[y] + k2u * dt / 2, v[y] + k2v * dt / 2, w[y] + k2w * dt / 2);
            k4u = f(u[y] + k3u * dt, v[y] + k3v * dt, w[y] + k3w * dt);
            k4v = g(u[y] + k3u * dt, v[y] + k3v * dt, w[y] + k3w * dt);
            k4w = h(u[y] + k3u * dt, v[y] + k3v * dt, w[y] + k3w * dt);
            ku = (k1u + 2 * k2u + 2 * k3u + k4u) / 6;
            kv = (k1v + 2 * k2v + 2 * k3v + k4v) / 6;
            kw = (k1w + 2 * k2w + 2 * k3w + k4w) / 6;
            unew[y] = u[y] + ku * dt + Du * dt * dx2inv * (u[y-1]+u[y-1]-2*u[y]);
            vnew[y] = v[y] + kv * dt + Dv * dt * dx2inv * (v[y-1]+v[y-1]-2*v[y]);
            wnew[y] = w[y] + kw * dt;
            for (y=0; y< lnfin; y++){
              uold[y] = u[y];
              u[y] = unew[y];
              v[y] = vnew[y];
              w[y] = wnew[y];
            }
            y = 0;
            pv[y] = 0.0;
            y = ln/2-1;
            pv[y] = 0.0;
            for (y=1; y< ln/2-1; y++){
              pv[y] = -((u[y]-uold[y])/dt)/((u[(y+1)%lnfin]-u[(y-1+ln)%lnfin])/2.0/dx);
            }
            y = lnfin-ln/2;
            pv[y] = 0.0;
            y = lnfin-1;
            pv[y] = 0.0;
            for (y= lnfin-ln/2+1; y< lnfin-1; y++){
              pv[y] = -((u[y]-uold[y])/dt)/((u[(y+1)%lnfin]-u[(y-1+ln)%lnfin])/2.0/dx);
            }
          }          
        }
        for (y=0; y<lnfin; y++){
          fprintf(outputfile, "%g\t", (u[y]));
        }
        for (y=0; y<lnfin; y++){
          fprintf(outputfile, "%g\t", (pv[y]));
        }
        for (y=0; y<lnfin; y++){
          fprintf(outputfile, "%g\t", (v[y]));
        }
        for (y=0; y<lnfin; y++){
          fprintf(outputfile, "%g\t", (w[y]));
        }
        for (y=0; y<lnfin; y++){
          fprintf(outputfile, "%g\t", (uold[y]));
        }
        fprintf(outputfile, "\n");
        printf("%d %d %g %g %d\n",rep,n1,u[1],dx*ln,ln);
      }
      fclose(outputfile);
    }
  //}
  return 0;
}

//file7.c
//Numerical simulation code using Neumann boundary conditions with multiple domains (expanding or shrinking) with different control paramters
//Note that in the codes, we used the different scales than in the manuscript.
//L = 3.2 and T = 6.2 in the codes corresponds to L = 1.8 and T = 8.6, respectively.

#include <stdio.h>
#include <stdlib.h>
#include <time.h>
#include "MT.h"// can be obtained from https://omitakahiro.github.io/random/code/MT.h

//Parallelization
//#ifdef _OPENMP
//#include <omp.h>
//#endif

#define _USE_MATH_DEFINES

#include <math.h>

#define stepsperout 620 //time steps per output
#define outsteps 1000 //number of outputs

double f(double u, double v, double w){
  return 0.57 *u - v + w - u*u*u;
}

double g(double u, double v, double w){
  return 1.5 * u - 2.03 * v - v*v*v;
}

double h(double u, double v, double w){
  return - u - 0.03 * w - w*w*w; 
}

double f1(double u, double v, double w){
  return 0.57 *u - v + w - u*u*u - (0.7+0.0238)*u;
}

double g1(double u, double v, double w){
  return 1.5 * u - 2.03 * v - v*v*v - (0.7+0.0238)*v;
}

double h1(double u, double v, double w){
  return - u - 0.03 * w - w*w*w - (0.7+0.0238)*w;  
}

int main(void) {
  //#pragma omp parallel
  //{
    int rep;
    //#pragma omp for
    for(rep=0;rep<1;rep=rep+1) {
      double dt = 0.001;//discrete time step
      int lninit = 4;//initial lattice number
      int lnfin = 32*8;//final lattice number
      int lnhalf;
      double s = 1.1867777442770433;//spatial scaling paramter throughout the study
      double Du = 0.03 * s;//diffusion coefficient for u
      double Dv = 1.0 * s;//diffusion coefficient for v
      int n1,m,y;
      double dx = 0.1;//spatial lattice size
      double dx2inv = 1.0/dx/dx;
      double t;

      double *u, *v, *w, *uold, *pv;
      double *unew, *vnew, *wnew;
      double *checker;
      u = (double*) malloc(sizeof(double) *  lnfin);
      v = (double*) malloc(sizeof(double) *  lnfin);
      w = (double*) malloc(sizeof(double) *  lnfin);
      unew = (double*) malloc(sizeof(double) *  lnfin);
      vnew = (double*) malloc(sizeof(double) *  lnfin);
      wnew = (double*) malloc(sizeof(double) *  lnfin);
      uold = (double*) malloc(sizeof(double) *  lnfin);
      pv = (double*) malloc(sizeof(double) *  lnfin);
      checker = (double*) malloc(sizeof(double) *  lnfin);

      srand((unsigned)time(NULL)+rep*100);
      for (n1 = 0; n1 < lnfin; n1++){
          u[n1] = (0.02 * rand()/RAND_MAX -0.01)*50;
          v[n1] = (0.02 * rand()/RAND_MAX -0.01)*50;
          w[n1] = (0.02 * rand()/RAND_MAX -0.01)*50;
          unew[n1] = 0.0;
          vnew[n1] = 0.0;
          wnew[n1] = 0.0;
          pv[n1] = 0.0;
          checker[n1] = 0.0;
      }

      FILE *outputfile;
      char filename[50];
      sprintf(filename, "turing_feedback-%03d.txt",rep);
      outputfile = fopen(filename, "w");

      for (y=0; y<lnfin; y++){
        fprintf(outputfile, "%g\t", (u[y]));
      }
      for (y=0; y<lnfin; y++){
        fprintf(outputfile, "%g\t", (pv[y]));
      }
      for (y=0; y<lnfin; y++){
        fprintf(outputfile, "%g\t", (v[y]));
      }
      for (y=0; y<lnfin; y++){
        fprintf(outputfile, "%g\t", (w[y]));
      }
      for (y=0; y<lnfin; y++){
        fprintf(outputfile, "%g\t", (checker[y]));
      }
      fprintf(outputfile, "\n");

      double ku,kv,kw, k1u, k1v, k1w, k2u, k2v, k2w, k3u, k3v, k3w, k4u, k4v, k4w;
      n1 = 0;
      while(n1 < outsteps) {
        n1++;
        for (m = 0; m < stepsperout; m++){
          t = dt * (n1 * stepsperout + m);
          lnhalf = (int)(lninit + (lnfin/2-lninit)*t/30.28);
          if(lnhalf > lnfin/2) {
            lnhalf = lnfin/2;
          }
          for (y= 1; y< lnhalf; y++){
            k1u = f(u[y], v[y], w[y]);
            k1v = g(u[y], v[y], w[y]);
            k1w = h(u[y], v[y], w[y]);
            k2u = f(u[y] + k1u * dt / 2, v[y] + k1v * dt / 2, w[y] + k1w * dt / 2);
            k2v = g(u[y] + k1u * dt / 2, v[y] + k1v * dt / 2, w[y] + k1w * dt / 2);
            k2w = h(u[y] + k1u * dt / 2, v[y] + k1v * dt / 2, w[y] + k1w * dt / 2);
            k3u = f(u[y] + k2u * dt / 2, v[y] + k2v * dt / 2, w[y] + k2w * dt / 2);
            k3v = g(u[y] + k2u * dt / 2, v[y] + k2v * dt / 2, w[y] + k2w * dt / 2);
            k3w = h(u[y] + k2u * dt / 2, v[y] + k2v * dt / 2, w[y] + k2w * dt / 2);
            k4u = f(u[y] + k3u * dt, v[y] + k3v * dt, w[y] + k3w * dt);
            k4v = g(u[y] + k3u * dt, v[y] + k3v * dt, w[y] + k3w * dt);
            k4w = h(u[y] + k3u * dt, v[y] + k3v * dt, w[y] + k3w * dt);
            ku = (k1u + 2 * k2u + 2 * k3u + k4u) / 6;
            kv = (k1v + 2 * k2v + 2 * k3v + k4v) / 6;
            kw = (k1w + 2 * k2w + 2 * k3w + k4w) / 6;
            unew[y] = u[y] + ku * dt + Du * dt * dx2inv * (u[y+1]+u[y-1]-2*u[y]);
            vnew[y] = v[y] + kv * dt + Dv * dt * dx2inv * (v[y+1]+v[y-1]-2*v[y]);
            wnew[y] = w[y] + kw * dt;
            checker[y] = 1.0;
          }
          for (y= lnhalf; y< lnfin-lnhalf; y++){
            k1u = f1(u[y], v[y], w[y]);
            k1v = g1(u[y], v[y], w[y]);
            k1w = h1(u[y], v[y], w[y]);
            k2u = f1(u[y] + k1u * dt / 2, v[y] + k1v * dt / 2, w[y] + k1w * dt / 2);
            k2v = g1(u[y] + k1u * dt / 2, v[y] + k1v * dt / 2, w[y] + k1w * dt / 2);
            k2w = h1(u[y] + k1u * dt / 2, v[y] + k1v * dt / 2, w[y] + k1w * dt / 2);
            k3u = f1(u[y] + k2u * dt / 2, v[y] + k2v * dt / 2, w[y] + k2w * dt / 2);
            k3v = g1(u[y] + k2u * dt / 2, v[y] + k2v * dt / 2, w[y] + k2w * dt / 2);
            k3w = h1(u[y] + k2u * dt / 2, v[y] + k2v * dt / 2, w[y] + k2w * dt / 2);
            k4u = f1(u[y] + k3u * dt, v[y] + k3v * dt, w[y] + k3w * dt);
            k4v = g1(u[y] + k3u * dt, v[y] + k3v * dt, w[y] + k3w * dt);
            k4w = h1(u[y] + k3u * dt, v[y] + k3v * dt, w[y] + k3w * dt);
            ku = (k1u + 2 * k2u + 2 * k3u + k4u) / 6;
            kv = (k1v + 2 * k2v + 2 * k3v + k4v) / 6;
            kw = (k1w + 2 * k2w + 2 * k3w + k4w) / 6;
            unew[y] = u[y] + ku * dt + Du * dt * dx2inv * (u[y+1]+u[y-1]-2*u[y]);
            vnew[y] = v[y] + kv * dt + Dv * dt * dx2inv * (v[y+1]+v[y-1]-2*v[y]);
            wnew[y] = w[y] + kw * dt;
            checker[y] = 0.0;
          }
          for (y= lnfin-lnhalf; y<lnfin-1;y++){
            k1u = f(u[y], v[y], w[y]);
            k1v = g(u[y], v[y], w[y]);
            k1w = h(u[y], v[y], w[y]);
            k2u = f(u[y] + k1u * dt / 2, v[y] + k1v * dt / 2, w[y] + k1w * dt / 2);
            k2v = g(u[y] + k1u * dt / 2, v[y] + k1v * dt / 2, w[y] + k1w * dt / 2);
            k2w = h(u[y] + k1u * dt / 2, v[y] + k1v * dt / 2, w[y] + k1w * dt / 2);
            k3u = f(u[y] + k2u * dt / 2, v[y] + k2v * dt / 2, w[y] + k2w * dt / 2);
            k3v = g(u[y] + k2u * dt / 2, v[y] + k2v * dt / 2, w[y] + k2w * dt / 2);
            k3w = h(u[y] + k2u * dt / 2, v[y] + k2v * dt / 2, w[y] + k2w * dt / 2);
            k4u = f(u[y] + k3u * dt, v[y] + k3v * dt, w[y] + k3w * dt);
            k4v = g(u[y] + k3u * dt, v[y] + k3v * dt, w[y] + k3w * dt);
            k4w = h(u[y] + k3u * dt, v[y] + k3v * dt, w[y] + k3w * dt);
            ku = (k1u + 2 * k2u + 2 * k3u + k4u) / 6;
            kv = (k1v + 2 * k2v + 2 * k3v + k4v) / 6;
            kw = (k1w + 2 * k2w + 2 * k3w + k4w) / 6;
            unew[y] = u[y] + ku * dt + Du * dt * dx2inv * (u[y+1]+u[y-1]-2*u[y]);
            vnew[y] = v[y] + kv * dt + Dv * dt * dx2inv * (v[y+1]+v[y-1]-2*v[y]);
            wnew[y] = w[y] + kw * dt;
            checker[y] = 1.0;
          }
          for (y=1; y< lnfin-1; y++){
            uold[y] = u[y];
            u[y] = unew[y];
            v[y] = vnew[y];
            w[y] = wnew[y];
          }
          uold[0] = uold[1];
          u[0] = u[1];
          v[0] = v[1];
          w[0] = w[1];
          uold[lnfin-1] = uold[lnfin-2];
          u[lnfin-1] = u[lnfin-2];
          v[lnfin-1] = v[lnfin-2];
          w[lnfin-1] = w[lnfin-2];
          y = 0;
          pv[y] = 0.0;
          y = lnfin-1;
          pv[y] = 0.0;
          for (y= 1; y< lnfin-1; y++){
            pv[y] = -((u[y]-uold[y])/dt)/((u[(y+1)%lnfin]-u[(y-1+lnfin)%lnfin])/2.0/dx);
          }
        }
        for (y=0; y<lnfin; y++){
          fprintf(outputfile, "%g\t", (u[y]));
        }
        for (y=0; y<lnfin; y++){
          fprintf(outputfile, "%g\t", (pv[y]));
        }
        for (y=0; y<lnfin; y++){
          fprintf(outputfile, "%g\t", (v[y]));
        }
        for (y=0; y<lnfin; y++){
          fprintf(outputfile, "%g\t", (w[y]));
        }
        for (y=0; y<lnfin; y++){
          fprintf(outputfile, "%g\t", (checker[y]));
        }
        fprintf(outputfile, "\n");
        printf("%d %d\n",rep,n1);
      }
      fclose(outputfile);
    }
  //}
  return 0;
}

//file8.c
//Numerical simulation code with advection term.
//Note that in the codes, we used the different scales than in the manuscript.
//L = 3.2 and T = 6.2 in the codes corresponds to L = 1.8 and T = 8.6, respectively.

#include <stdio.h>
#include <stdlib.h>
#include <time.h>
#include "MT.h"// can be obtained from https://omitakahiro.github.io/random/code/MT.h

//Parallelization
//#ifdef _OPENMP
//#include <omp.h>
//#endif

#define _USE_MATH_DEFINES

#include <math.h>

#define stepsperout 620 //time steps per output
#define outsteps 1000 //number of outputs

double f(double u, double v, double w){
  return 0.57 *u - v + w - u*u*u;
}

double g(double u, double v, double w){
  return 1.5 * u - 2.03 * v - v*v*v;
}

double h(double u, double v, double w){
  return - u - 0.03 * w - w*w*w;
}

int main(void) {
  //#pragma omp parallel
  //{
    int rep;
    //#pragma omp for
    for(rep=0;rep<1;rep=rep+1) {
      double dt = 0.001;//discrete time step
      int ln = 32*8;//lattice number
      double dx = 0.1;//spatial lattice size
      double dx2inv = 1.0/dx/dx;
      double dxinv = 1.0/dx;
      double s = 1.1867777442770433;//spatial scaling paramter throughout the study
      double Du = 0.03 * s;//diffusion coefficient for u
      double Dv = 1.0 * s;//diffusion coefficient for v
      double cv;
      double cvr = 0.005;//advection term gradient
      int n1,m,y;

      double *u, *v, *w, *uold, *pv;
      double *unew, *vnew, *wnew;
      u = (double*) malloc(sizeof(double) *  ln);
      v = (double*) malloc(sizeof(double) *  ln);
      w = (double*) malloc(sizeof(double) *  ln);
      unew = (double*) malloc(sizeof(double) *  ln);
      vnew = (double*) malloc(sizeof(double) *  ln);
      wnew = (double*) malloc(sizeof(double) *  ln);
      uold = (double*) malloc(sizeof(double) *  ln);
      pv = (double*) malloc(sizeof(double) *  ln);

      srand((unsigned)time(NULL)+rep*100);
      for (n1 = 0; n1 < ln; n1++){
        u[n1] = (0.02 * rand()/RAND_MAX -0.01)*50.0;
        v[n1] = (0.02 * rand()/RAND_MAX -0.01)*50.0;
        w[n1] = (0.02 * rand()/RAND_MAX -0.01)*50.0;
        pv[n1] = 0.0;
      }
      u[0] = u[1];
      v[0] = v[1];
      w[0] = w[1];
      u[ln-1] = u[ln-2];
      v[ln-1] = v[ln-2];
      w[ln-1] = w[ln-2];

      FILE *outputfile;
      char filename[50];
      sprintf(filename, "turing_feedback-%03d.txt",rep);
      outputfile = fopen(filename, "w");
      for (y=1; y<ln-1; y++){
        fprintf(outputfile, "%g\t", (u[y]));
      }
      for (y=1; y<ln-1; y++){
        fprintf(outputfile, "%g\t", (pv[y]));
      }
      for (y=1; y<ln-1; y++){
        fprintf(outputfile, "%g\t", (v[y]));
      }
      for (y=1; y<ln-1; y++){
        fprintf(outputfile, "%g\t", (w[y]));
      }
      fprintf(outputfile, "\n");

      double ku,kv,kw,k1u, k1v, k1w, k2u, k2v, k2w, k3u, k3v, k3w, k4u, k4v, k4w;

      n1 = 0;
      while(n1 < outsteps) {
        n1++;
        for (m = 0; m < stepsperout; m++){
          //計算をそのまま回す
          for (y=1; y< ln-1; y++){
            k1u = f(u[y], v[y], w[y]);
            k1v = g(u[y], v[y], w[y]);
            k1w = h(u[y], v[y], w[y]);
            k2u = f(u[y] + k1u * dt / 2, v[y] + k1v * dt / 2, w[y] + k1w * dt / 2);
            k2v = g(u[y] + k1u * dt / 2, v[y] + k1v * dt / 2, w[y] + k1w * dt / 2);
            k2w = h(u[y] + k1u * dt / 2, v[y] + k1v * dt / 2, w[y] + k1w * dt / 2);
            k3u = f(u[y] + k2u * dt / 2, v[y] + k2v * dt / 2, w[y] + k2w * dt / 2);
            k3v = g(u[y] + k2u * dt / 2, v[y] + k2v * dt / 2, w[y] + k2w * dt / 2);
            k3w = h(u[y] + k2u * dt / 2, v[y] + k2v * dt / 2, w[y] + k2w * dt / 2);
            k4u = f(u[y] + k3u * dt, v[y] + k3v * dt, w[y] + k3w * dt);
            k4v = g(u[y] + k3u * dt, v[y] + k3v * dt, w[y] + k3w * dt);
            k4w = h(u[y] + k3u * dt, v[y] + k3v * dt, w[y] + k3w * dt);
            ku = (k1u + 2 * k2u + 2 * k3u + k4u) / 6;
            kv = (k1v + 2 * k2v + 2 * k3v + k4v) / 6;
            kw = (k1w + 2 * k2w + 2 * k3w + k4w) / 6;
            cv = cvr * dx * (y - 0.5*(ln-1));
            unew[y] = u[y] + ku * dt + Du * dt * dx2inv * (u[(y+1)%ln]+u[(y-1+ln)%ln]-2*u[y]) - cv * dt * dxinv * 0.5 * (u[(y+1)%ln]-u[(y-1+ln)%ln]);
            vnew[y] = v[y] + kv * dt + Dv * dt * dx2inv * (v[(y+1)%ln]+v[(y-1+ln)%ln]-2*v[y]) - cv * dt * dxinv * 0.5 * (v[(y+1)%ln]-v[(y-1+ln)%ln]);
            wnew[y] = w[y] + kw * dt;
          }
          for (y=1; y< ln-1; y++){
            uold[y] = u[y];
            u[y] = unew[y];
            v[y] = vnew[y];
            w[y] = wnew[y];
          }
          u[0] = u[1];
          v[0] = v[1];
          w[0] = w[1];
          u[ln-1] = u[ln-2];
          v[ln-1] = v[ln-2];
          w[ln-1] = w[ln-2];
          for (y=1; y< ln-1; y++){
            pv[y] = -((u[y]-uold[y])/dt)/((u[(y+1)%ln]-u[(y-1+ln)%ln])/2.0/dx);
          }
        }
        for (y=1; y<ln-1; y++){
          fprintf(outputfile, "%g\t", (u[y]));
        }
        for (y=1; y<ln-1; y++){
          fprintf(outputfile, "%g\t", (pv[y]));
        }
        for (y=1; y<ln-1; y++){
          fprintf(outputfile, "%g\t", (v[y]));
        }
        for (y=1; y<ln-1; y++){
          fprintf(outputfile, "%g\t", (w[y]));
        }
        fprintf(outputfile, "\n");
        printf("%d %d\n",rep,n1);
      }
      fclose(outputfile);
    }
  //}
  return 0;
}

//file9.c
//Numerical simulation code with changing diameter in a simple setting.
//Note that in the codes, we used the different scales than in the manuscript.
//L = 3.2 and T = 6.2 in the codes corresponds to L = 1.8 and T = 8.6, respectively.

#include <stdio.h>
#include <stdlib.h>
#include <time.h>
#include "MT.h"// can be obtained from https://omitakahiro.github.io/random/code/MT.h

//Parallelization
//#ifdef _OPENMP
//#include <omp.h>
//#endif

#define _USE_MATH_DEFINES

#include <math.h>

#define outsteps 1000//number of outputs

double f(double u, double v, double w){
  return 0.57 *u - v + w - u*u*u;
}

double g(double u, double v, double w){
  return 1.5 * u - 2.03 * v - v*v*v;
}

double h(double u, double v, double w){
  return - u - 0.03 * w - w*w*w; 
}

int main(void) {
  //#pragma omp parallel
  //{
    int rep;
    //#pragma omp for
    for(rep=0;rep<2*2;rep=rep+1) {
      double dt = 0.001;//discrete time step
      int ln = 32*32+2;//lattice number
      double dx = 0.1;//spatial lattice size
      double dx2inv = 1.0/dx/dx;
      double dxinv = 1.0/dx;
      double s = 1.1867777442770433;//spatial scaling paramter throughout the study
      double Du = 0.03 * s;//diffusion coefficient for u
      double Dv = 1.0 * s;//diffusion coefficient for v
      int stepsperout = 620;//time steps per output
      double rmin = 0.02*0.5*3.2/1.8;//mininum radius
      double rmax = 0.2*0.5*3.2/1.8;//maximam radius
      double c = (rmax-rmin)/dx/ln;//gradient of radius
      int n1,m,y;
      double r;

      double *u, *v, *w, *uold, *pv;
      double *unew, *vnew, *wnew;
      u = (double*) malloc(sizeof(double) *  ln);
      v = (double*) malloc(sizeof(double) *  ln);
      w = (double*) malloc(sizeof(double) *  ln);
      unew = (double*) malloc(sizeof(double) *  ln);
      vnew = (double*) malloc(sizeof(double) *  ln);
      wnew = (double*) malloc(sizeof(double) *  ln);
      uold = (double*) malloc(sizeof(double) *  ln);
      pv = (double*) malloc(sizeof(double) *  ln);

      srand((unsigned)time(NULL)+rep*100);
      for (n1 = 0; n1 < ln; n1++){
        u[n1] = 50.0*(0.02 * rand()/RAND_MAX -0.01);
        v[n1] = 50.0*(0.02 * rand()/RAND_MAX -0.01);
        w[n1] = 50.0*(0.02 * rand()/RAND_MAX -0.01);
        pv[n1] = 0.0;
      }
      u[0] = u[2];
      v[0] = v[2];
      w[0] = w[2];
      u[ln-1] = u[ln-3];
      v[ln-1] = v[ln-3];
      w[ln-1] = w[ln-3];

      double ku,kv,kw, k1u, k1v, k1w, k2u, k2v, k2w, k3u, k3v, k3w, k4u, k4v, k4w;
      
      n1 = 0;

      FILE *outputfile;
      char filename[50];¥
      sprintf(filename, "turing_feedback-%03d.txt",rep);
      outputfile = fopen(filename, "w");
      for (y=1; y<ln-1; y++){
        fprintf(outputfile, "%g\t", (u[y]));
      }
      for (y=1; y<ln-1; y++){
        fprintf(outputfile, "%g\t", (pv[y]));
      }
      for (y=1; y<ln-1; y++){
        fprintf(outputfile, "%g\t", (v[y]));
      }
      for (y=1; y<ln-1; y++){
        fprintf(outputfile, "%g\t", (w[y]));
      }
      fprintf(outputfile, "\n");

      while(n1 < outsteps) {
        n1++;
        for (m = 0; m < stepsperout; m++){
          for (y=1; y< ln-1; y++){
            k1u = f(u[y], v[y], w[y]);
            k1v = g(u[y], v[y], w[y]);
            k1w = h(u[y], v[y], w[y]);
            k2u = f(u[y] + k1u * dt / 2, v[y] + k1v * dt / 2, w[y] + k1w * dt / 2);
            k2v = g(u[y] + k1u * dt / 2, v[y] + k1v * dt / 2, w[y] + k1w * dt / 2);
            k2w = h(u[y] + k1u * dt / 2, v[y] + k1v * dt / 2, w[y] + k1w * dt / 2);
            k3u = f(u[y] + k2u * dt / 2, v[y] + k2v * dt / 2, w[y] + k2w * dt / 2);
            k3v = g(u[y] + k2u * dt / 2, v[y] + k2v * dt / 2, w[y] + k2w * dt / 2);
            k3w = h(u[y] + k2u * dt / 2, v[y] + k2v * dt / 2, w[y] + k2w * dt / 2);
            k4u = f(u[y] + k3u * dt, v[y] + k3v * dt, w[y] + k3w * dt);
            k4v = g(u[y] + k3u * dt, v[y] + k3v * dt, w[y] + k3w * dt);
            k4w = h(u[y] + k3u * dt, v[y] + k3v * dt, w[y] + k3w * dt);
            ku = (k1u + 2 * k2u + 2 * k3u + k4u) / 6;
            kv = (k1v + 2 * k2v + 2 * k3v + k4v) / 6;
            kw = (k1w + 2 * k2w + 2 * k3w + k4w) / 6;
            r = (rmax-rmin)*y/ln + rmin;
            unew[y] = u[y] + ku * dt + Du / (1.0+c*c) * dt * dx2inv * (u[(y+1)%ln]+u[(y-1+ln)%ln]-2*u[y]) + Du * 1.0*c/r/(1.0+c*c) * dt * dxinv * 0.5 * (u[(y+1)%ln]-u[(y-1+ln)%ln]) ;
            vnew[y] = v[y] + kv * dt + Dv / (1.0+c*c) * dt * dx2inv * (v[(y+1)%ln]+v[(y-1+ln)%ln]-2*v[y]) + Dv * 1.0*c/r/(1.0+c*c) * dt * dxinv * 0.5 * (v[(y+1)%ln]-v[(y-1+ln)%ln]) ;
            wnew[y] = w[y] + kw * dt;
          }
          for (y=1; y< ln-1; y++){
            uold[y] = u[y];
            u[y] = unew[y];
            v[y] = vnew[y];
            w[y] = wnew[y];
          }
          u[0] = u[2];
          v[0] = v[2];
          w[0] = w[2];
          u[ln-1] = u[ln-3];
          v[ln-1] = v[ln-3];
          w[ln-1] = w[ln-3];
          for (y=1; y< ln-1; y++){
            pv[y] = -((u[y]-uold[y])/dt)/((u[(y+1)%ln]-u[(y-1+ln)%ln])/2.0/dx);
          }
        }
        printf("%d %d\n",rep,n1);          
        for (y=1; y<ln-1; y++){
          fprintf(outputfile, "%g\t", (u[y]));
        }
        for (y=1; y<ln-1; y++){
          fprintf(outputfile, "%g\t", (pv[y]));
        }
        for (y=1; y<ln-1; y++){
          fprintf(outputfile, "%g\t", (v[y]));
        }
        for (y=1; y<ln-1; y++){
          fprintf(outputfile, "%g\t", (w[y]));
        }
        fprintf(outputfile, "\n");
        }
      fclose(outputfile);
    }
  //}
  return 0;
}

//file10.c
//Numerical simulation code with changing diameter in a realistic setting.

#include <stdio.h>
#include <stdlib.h>
#include <time.h>
#include "MT.h"// can be obtained from https://omitakahiro.github.io/random/code/MT.h

//Parallelization
//#ifdef _OPENMP
//#include <omp.h>
//#endif

#define _USE_MATH_DEFINES

#include <math.h>

#define stepsperout 62000//time steps per output
#define outsteps 1000//number of outputs

double f(double u, double v, double w){
  return 0.57 *u - v + w - u*u*u;
}

double g(double u, double v, double w){
  return 1.5 * u - 2.03 * v - v*v*v;
}

double h(double u, double v, double w){
  return - u - 0.03 * w - w*w*w;
}

int main(void) {
  //#pragma omp parallel
  //{
    int rep;
    //#pragma omp for
    for(rep=0;rep<1;rep=rep+1) {
      double dt = 0.00001;//discrete time step
      int ln = 36+2560+36+2;// 0 1-36 31-2596 2597-2632 2633//lattice number
      // 0.2025 14 (~1.8x8) 0.2025 cm
      // 0.36 25.6 0.36
      double dx = 0.01;//spatial lattice size
      double dx2inv = 1.0/dx/dx;
      double dxinv = 1.0/dx;
      double s = 1.1867777442770433;//spatial scaling paramter throughout the study
      double Du = 0.03 * s;//diffusion coefficient for u
      double Dv = 1.0 * s;//diffusion coefficient for v
      double rmax = 0.2*0.5*3.2/1.8;
      double rmin = 0.05*0.5*3.2/1.8;
      double c = (0.1-0.025)/0.2025;
      int n1,m,y;
      double r;

      double *u, *v, *w, *uold, *pv;
      double *unew, *vnew, *wnew;
      u = (double*) malloc(sizeof(double) *  ln);
      v = (double*) malloc(sizeof(double) *  ln);
      w = (double*) malloc(sizeof(double) *  ln);
      unew = (double*) malloc(sizeof(double) *  ln);
      vnew = (double*) malloc(sizeof(double) *  ln);
      wnew = (double*) malloc(sizeof(double) *  ln);
      uold = (double*) malloc(sizeof(double) *  ln);
      pv = (double*) malloc(sizeof(double) *  ln);

      srand((unsigned)time(NULL)+rep*100);
      for (n1 = 0; n1 < ln; n1++){
        u[n1] = 50*(0.02 * rand()/RAND_MAX -0.01);
        v[n1] = 50*(0.02 * rand()/RAND_MAX -0.01);
        w[n1] = 50*(0.02 * rand()/RAND_MAX -0.01);
        pv[n1] = 0.0;
      }
      u[0] = u[2];
      v[0] = v[2];
      w[0] = w[2];
      u[ln-1] = u[ln-3];
      v[ln-1] = v[ln-3];
      w[ln-1] = w[ln-3];

      double ku,kv,kw,k1u, k1v, k1w, k2u, k2v, k2w, k3u, k3v, k3w, k4u, k4v, k4w;
      double l2normcutoff = 1.0e-10;
      l2normold = 1000.0;
      l2normdif = 1000.0;
      n1 = 0;
      while(n1 < outsteps) {
        FILE *outputfile;  
        char filename[50]; 
        sprintf(filename, "turing_feedback-%03d-%04d.txt",rep,n1);
        outputfile = fopen(filename, "w");
        for (y=1; y<ln-1; y++){
          fprintf(outputfile, "%g\t", (u[y]));
        }
        for (y=1; y<ln-1; y++){
          fprintf(outputfile, "%g\t", (pv[y]));
        }
        for (y=1; y<ln-1; y++){
          fprintf(outputfile, "%g\t", (v[y]));
        }
        for (y=1; y<ln-1; y++){
          fprintf(outputfile, "%g\t", (w[y]));
        }
        fprintf(outputfile, "\n");
        fclose(outputfile);
        
        n1++;
        for (m = 0; m < stepsperout; m++){
          for (y=1; y<=ln-1; y++){
            k1u = f(u[y], v[y], w[y]);
            k1v = g(u[y], v[y], w[y]);
            k1w = h(u[y], v[y], w[y]);
            k2u = f(u[y] + k1u * dt / 2, v[y] + k1v * dt / 2, w[y] + k1w * dt / 2);
            k2v = g(u[y] + k1u * dt / 2, v[y] + k1v * dt / 2, w[y] + k1w * dt / 2);
            k2w = h(u[y] + k1u * dt / 2, v[y] + k1v * dt / 2, w[y] + k1w * dt / 2);
            k3u = f(u[y] + k2u * dt / 2, v[y] + k2v * dt / 2, w[y] + k2w * dt / 2);
            k3v = g(u[y] + k2u * dt / 2, v[y] + k2v * dt / 2, w[y] + k2w * dt / 2);
            k3w = h(u[y] + k2u * dt / 2, v[y] + k2v * dt / 2, w[y] + k2w * dt / 2);
            k4u = f(u[y] + k3u * dt, v[y] + k3v * dt, w[y] + k3w * dt);
            k4v = g(u[y] + k3u * dt, v[y] + k3v * dt, w[y] + k3w * dt);
            k4w = h(u[y] + k3u * dt, v[y] + k3v * dt, w[y] + k3w * dt);
            ku = (k1u + 2 * k2u + 2 * k3u + k4u) / 6;
            kv = (k1v + 2 * k2v + 2 * k3v + k4v) / 6;
            kw = (k1w + 2 * k2w + 2 * k3w + k4w) / 6;
            unew[y] = u[y] + ku * dt + Du * dt * dx2inv * (u[(y+1)%ln]+u[(y-1+ln)%ln]-2*u[y]);
            vnew[y] = v[y] + kv * dt + Dv * dt * dx2inv * (v[(y+1)%ln]+v[(y-1+ln)%ln]-2*v[y]);
            wnew[y] = w[y] + kw * dt;
            //0 1-36 31-2596 2597-2632 2633
            //0 1-30 31-2590 2591-2620 2621
            if(y<=36){
              r = (rmax-rmin)*(y-0.5)/36 + rmin;
              c = 0.75;
              unew[y] = u[y] + ku * dt + Du / (1.0+c*c) * dt * dx2inv * (u[(y+1)%ln]+u[(y-1+ln)%ln]-2*u[y]) + Du * 1.0*c/r/(1.0+c*c) * dt * dxinv * 0.5 * (u[(y+1)%ln]-u[(y-1+ln)%ln]) ;
              vnew[y] = v[y] + kv * dt + Dv / (1.0+c*c) * dt * dx2inv * (v[(y+1)%ln]+v[(y-1+ln)%ln]-2*v[y]) + Dv * 1.0*c/r/(1.0+c*c) * dt * dxinv * 0.5 * (v[(y+1)%ln]-v[(y-1+ln)%ln]) ;
              wnew[y] = w[y] + kw * dt;
            }
            if(y>=2597){
              r = (rmax-rmin)*(2632.5-y)/36 + rmin;
              c = -0.75;
              unew[y] = u[y] + ku * dt + Du / (1.0+c*c) * dt * dx2inv * (u[(y+1)%ln]+u[(y-1+ln)%ln]-2*u[y]) + Du * 1.0*c/r/(1.0+c*c) * dt * dxinv * 0.5 * (u[(y+1)%ln]-u[(y-1+ln)%ln]) ;
              vnew[y] = v[y] + kv * dt + Dv / (1.0+c*c) * dt * dx2inv * (v[(y+1)%ln]+v[(y-1+ln)%ln]-2*v[y]) + Dv * 1.0*c/r/(1.0+c*c) * dt * dxinv * 0.5 * (v[(y+1)%ln]-v[(y-1+ln)%ln]) ;
              wnew[y] = w[y] + kw * dt;
            }
          }
          for (y=1; y< ln-1; y++){
            uold[y] = u[y];
            u[y] = unew[y];
            v[y] = vnew[y];
            w[y] = wnew[y];
          }
          u[0] = u[2];
          v[0] = v[2];
          w[0] = w[2];
          u[ln-1] = u[ln-3];
          v[ln-1] = v[ln-3];
          w[ln-1] = w[ln-3];
          for (y=1; y< ln-1; y++){
            pv[y] = -((u[y]-uold[y])/dt)/((u[(y+1)%ln]-u[(y-1+ln)%ln])/2.0/dx);
          }
        }
        printf("%d %d %g\n",rep,n1,u[ln/2]);
      }
    }
  //}
  return 0;
}

//file11.c
//Numerical simulation code with Dirichlet boundary conditions for u and v.

#include <stdio.h>
#include <stdlib.h>
#include <time.h>
#include "MT.h"// can be obtained from https://omitakahiro.github.io/random/code/MT.h

//Parallelization
//#ifdef _OPENMP
//#include <omp.h>
//#endif

#define _USE_MATH_DEFINES
#include <math.h>

#define outsteps 1000 //number of outputs

double f(double u, double v, double w){
    return 0.57 * u - v + w - u*u*u;
}

double g(double u, double v, double w){
    return 1.5 * u - 2.03 * v - v*v*v;
}

double h(double u, double v, double w){
    return - u - 0.03 * w - w*w*w;
}

int main(void) {
  #pragma omp parallel
  {
    int rep;
    #pragma omp for
    for(rep=0;rep<1;rep=rep+1) {
      int stepsperout = 620;//time steps per output
      double dt = 0.001;//timestep
      int ln = 32*8+2;//lattice number
      double dx = 0.1;//spatial lattice size
      double dx2inv = 1.0/dx/dx;
      double dxinv = 1.0/dx;
      double s = 1.1867777442770433;//spatial scaling paramter throughout the study
      double Du = 0.03 * s;//diffusion coefficient for u
      double Dv = 1.0 * s;//diffusion coefficient for v
      int n1,m,y,n,x,j;

      double ub = 0.0;//U boundary value
      double vb = 0.0;//V boundary value

      double *u, *v, *w, *uold, *pv;
      double *unew, *vnew, *wnew;
      u = (double*) malloc(sizeof(double) *  ln);
      v = (double*) malloc(sizeof(double) *  ln);
      w = (double*) malloc(sizeof(double) *  ln);
      unew = (double*) malloc(sizeof(double) *  ln);
      vnew = (double*) malloc(sizeof(double) *  ln);
      wnew = (double*) malloc(sizeof(double) *  ln);
      uold = (double*) malloc(sizeof(double) *  ln);
      pv = (double*) malloc(sizeof(double) *  ln);

      srand((unsigned)time(NULL)+rep*100);
      for (n1 = 0; n1 < ln; n1++){
        u[n1] = 50.0*(0.02 * rand()/RAND_MAX -0.01);
        v[n1] = 50.0*(0.02 * rand()/RAND_MAX -0.01);
        w[n1] = 50.0*(0.02 * rand()/RAND_MAX -0.01);
        pv[n1] = 0.0;
      }

      double k1u, k1v, k1w, k2u, k2v, k2w, k3u, k3v, k3w, k4u, k4v, k4w, u_reaction, v_reaction, w_reaction, u_diffusion, v_diffusion, w_diffusion, l2normdif, l2normold, l2norm;
      double l2normcutoff = 1.0e-10;
      l2normold = 1000.0;
      l2normdif = 1000.0;
      n1 = 0;
      FILE *outputfile;
      char filename[50];
      sprintf(filename, "turing_feedback-%03d.txt",rep);
      outputfile = fopen(filename, "w");
      while(n1 < outsteps) {
        for (y=0; y<ln; y++){
          fprintf(outputfile, "%g\t", (u[y]));
        }
        for (y=0; y<ln; y++){
          fprintf(outputfile, "%g\t", (pv[y]));
        }
        for (y=0; y<ln; y++){
          fprintf(outputfile, "%g\t", (v[y]));
        }
        for (y=0; y<ln; y++){
          fprintf(outputfile, "%g\t", (w[y]));
        }
        fprintf(outputfile, "\n");

        n1++;
        for (m = 0; m < stepsperout; m++){
          
          for (y=0; y< ln; y++){
            k1u = f(u[y], v[y], w[y]);
            k1v = g(u[y], v[y], w[y]);
            k1w = h(u[y], v[y], w[y]);
            k2u = f(u[y] + k1u * dt / 2, v[y] + k1v * dt / 2, w[y] + k1w * dt / 2);
            k2v = g(u[y] + k1u * dt / 2, v[y] + k1v * dt / 2, w[y] + k1w * dt / 2);
            k2w = h(u[y] + k1u * dt / 2, v[y] + k1v * dt / 2, w[y] + k1w * dt / 2);
            k3u = f(u[y] + k2u * dt / 2, v[y] + k2v * dt / 2, w[y] + k2w * dt / 2);
            k3v = g(u[y] + k2u * dt / 2, v[y] + k2v * dt / 2, w[y] + k2w * dt / 2);
            k3w = h(u[y] + k2u * dt / 2, v[y] + k2v * dt / 2, w[y] + k2w * dt / 2);
            k4u = f(u[y] + k3u * dt, v[y] + k3v * dt, w[y] + k3w * dt);
            k4v = g(u[y] + k3u * dt, v[y] + k3v * dt, w[y] + k3w * dt);
            k4w = h(u[y] + k3u * dt, v[y] + k3v * dt, w[y] + k3w * dt);
            unew[y] = u[y] + (k1u + 2 * k2u + 2 * k3u + k4u) / 6 * dt + Du * (u[(y+1)%ln]+u[(y-1+ln)%ln]-2*u[y])*dx2inv*dt;
            vnew[y] = v[y] + (k1v + 2 * k2v + 2 * k3v + k4v) / 6 * dt + Dv * (v[(y+1)%ln]+v[(y-1+ln)%ln]-2*v[y])*dx2inv*dt;
            wnew[y] = w[y] + (k1w + 2 * k2w + 2 * k3w + k4w) / 6 * dt;
          }
          for (y=0; y< ln; y++){
            uold[y] = u[y];
            u[y] = unew[y];
            v[y] = vnew[y];
            w[y] = wnew[y];          
          }

          u[0] = ub;
          v[0] = vb;
          w[0] = w[1];
          u[ln-1] = ub;
          v[ln-1] = vb;
          w[ln-1] = w[ln-2];
        
          for (y=0; y< ln; y++){
            pv[y] = -((u[y]-uold[y])/dt)/((u[(y+1)%ln]-u[(y-1+ln)%ln])/2.0/dx);
          }
        }

        printf("%d %d\n",rep,n1);
      }
      fclose(outputfile);
    }
  }
  return 0;
}

//file12.c
//Numerical simulation code with Dirichlet boundary conditions for u and v.

#include <stdio.h>
#include <stdlib.h>
#include <time.h>
#include "MT.h"// can be obtained from https://omitakahiro.github.io/random/code/MT.h

//Parallelization
//#ifdef _OPENMP
//#include <omp.h>
//#endif

#define _USE_MATH_DEFINES
#include <math.h>

#define outsteps 1000 //number of outputs

double f(double u, double v, double w){
    return 0.57 * u - v + w - u*u*u;
}

double g(double u, double v, double w){
    return 1.5 * u - 2.03 * v - v*v*v;
}

double h(double u, double v, double w){
    return - u - 0.03 * w - w*w*w;
}

int main(void) {
  #pragma omp parallel
  {
    int rep;
    #pragma omp for
    for(rep=0;rep<1;rep=rep+1) {
        int stepsperout = 620;//time steps per output
        double dt = 0.001;//timestep
        int ln = 32*8+2;//lattice number
        double dx = 0.1;//spatial lattice size
        double dx2inv = 1.0/dx/dx;
        double dxinv = 1.0/dx;
        double s = 1.1867777442770433;//spatial scaling paramter throughout the study
        double Du = 0.03 * s;//diffusion coefficient for u
        double Dv = 1.0 * s;//diffusion coefficient for v
        int n1,m,y,n,x,j;
        
        double ku = 1.0;//boundary produnction term coefficient for U
        double kv = 1.0;//boundary produnction term coefficient for V

        double *u, *v, *w, *uold, *pv;
        double *unew, *vnew, *wnew;
        u = (double*) malloc(sizeof(double) *  ln);
        v = (double*) malloc(sizeof(double) *  ln);
        w = (double*) malloc(sizeof(double) *  ln);
        unew = (double*) malloc(sizeof(double) *  ln);
        vnew = (double*) malloc(sizeof(double) *  ln);
        wnew = (double*) malloc(sizeof(double) *  ln);
        uold = (double*) malloc(sizeof(double) *  ln);
        pv = (double*) malloc(sizeof(double) *  ln);

        srand((unsigned)time(NULL)+rep*100);
        for (n1 = 0; n1 < ln; n1++){
          u[n1] = 50.0 * (0.02 * rand()/RAND_MAX -0.01);
          v[n1] = 50.0 * (0.02 * rand()/RAND_MAX -0.01);
          w[n1] = 50.0 * (0.02 * rand()/RAND_MAX -0.01);
          pv[n1] = 0.0;
        }

        FILE *outputfile1;
        char filename1[50];
        sprintf(filename1, "turing_feedback-single-%03d.txt",rep0);
        outputfile1 = fopen(filename1, "w");

        double k1u, k1v, k1w, k2u, k2v, k2w, k3u, k3v, k3w, k4u, k4v, k4w, u_reaction, v_reaction, w_reaction, u_diffusion, v_diffusion, w_diffusion, l2normdif, l2normold, l2norm;
        double l2normcutoff = 1.0e-10;
        l2normold = 1000.0;
        l2normdif = 1000.0;
        n1 = 0;
        while(n1 < outsteps) {
          for (y=0; y<ln; y++){
            fprintf(outputfile1, "%g\t", (u[y]));
          }
          for (y=0; y<ln; y++){
            fprintf(outputfile1, "%g\t", (pv[y]));
          }
          for (y=0; y<ln; y++){
            fprintf(outputfile1, "%g\t", (v[y]));
          }
          for (y=0; y<ln; y++){
            fprintf(outputfile1, "%g\t", (w[y]));
          }
          fprintf(outputfile1, "\n");

          n1++;
          for (m = 0; m < stepsperout; m++){
            for (y=0; y< ln; y++){
              k1u = f(u[y], v[y], w[y]);
              k1v = g(u[y], v[y], w[y]);
              k1w = h(u[y], v[y], w[y]);
              k2u = f(u[y] + k1u * dt / 2, v[y] + k1v * dt / 2, w[y] + k1w * dt / 2);
              k2v = g(u[y] + k1u * dt / 2, v[y] + k1v * dt / 2, w[y] + k1w * dt / 2);
              k2w = h(u[y] + k1u * dt / 2, v[y] + k1v * dt / 2, w[y] + k1w * dt / 2);
              k3u = f(u[y] + k2u * dt / 2, v[y] + k2v * dt / 2, w[y] + k2w * dt / 2);
              k3v = g(u[y] + k2u * dt / 2, v[y] + k2v * dt / 2, w[y] + k2w * dt / 2);
              k3w = h(u[y] + k2u * dt / 2, v[y] + k2v * dt / 2, w[y] + k2w * dt / 2);
              k4u = f(u[y] + k3u * dt, v[y] + k3v * dt, w[y] + k3w * dt);
              k4v = g(u[y] + k3u * dt, v[y] + k3v * dt, w[y] + k3w * dt);
              k4w = h(u[y] + k3u * dt, v[y] + k3v * dt, w[y] + k3w * dt);
              unew[y] = u[y] + (k1u + 2 * k2u + 2 * k3u + k4u) / 6 * dt + Du * (u[(y+1)%ln]+u[(y-1+ln)%ln]-2*u[y])*dx2inv*dt;
              vnew[y] = v[y] + (k1v + 2 * k2v + 2 * k3v + k4v) / 6 * dt + Dv * (v[(y+1)%ln]+v[(y-1+ln)%ln]-2*v[y])*dx2inv*dt;
              wnew[y] = w[y] + (k1w + 2 * k2w + 2 * k3w + k4w) / 6 * dt;
            }
            unew[1] += dt * ku;
            vnew[1] += dt * kv;
            unew[ln-2] += dt * ku;
            vnew[ln-2] += dt * kv;
            for (y=0; y< ln; y++){
              uold[y] = u[y];
              u[y] = unew[y];
              v[y] = vnew[y];
              w[y] = wnew[y];          
            }
            
            u[0] = u[1];
            v[0] = v[1];
            w[0] = w[1];
            u[ln-1] = u[ln-2];
            v[ln-1] = v[ln-2];
            w[ln-1] = w[ln-2];
            
            for (y=0; y< ln; y++){
              pv[y] = -((u[y]-uold[y])/dt)/((u[(y+1)%ln]-u[(y-1+ln)%ln])/2.0/dx);
            }
          }
          printf("%d %d\n",rep,n1);
        }
        for (y=0; y<ln; y++){
          fprintf(outputfile1, "%g\t", (u[y]));
        }
        for (y=0; y<ln; y++){
          fprintf(outputfile1, "%g\t", (pv[y]));
        }
        for (y=0; y<ln; y++){
          fprintf(outputfile1, "%g\t", (v[y]));
        }
        for (y=0; y<ln; y++){
          fprintf(outputfile1, "%g\t", (w[y]));
        }
        fprintf(outputfile1, "\n");
    }
  }
  return 0;
}
```
